# European ash pangenome reveals widespread structural variation and genetic basis of low ash dieback susceptibility

**DOI:** 10.1101/2025.07.16.665114

**Authors:** Daniel P. Wood, Mohammad Vatanparast, Dario Galanti, Catherine Gudgeon, Katherine Wheeler, Emma Curran, Levi Yant, Richard Whittet, Richard A. Nichols, Richard J. A. Buggs, Laura J. Kelly

## Abstract

European Ash (*Fraxinus excelsior*) is a keystone forest tree species, whose populations are being decimated by ash dieback disease (ADB). Uncovering the genetic basis of low susceptibility to this devastating disease relies on a comprehensive understanding of genetic variation present in *F. excelsior.* A linear reference genome from a single individual cannot contain the total sequence variability within a species, including its genic regions; a pangenome more fully captures total sequence content. In this study, we developed a *F. excelsior* pangenome reference using a new chromosomal-level phased linear reference genome, and *de novo* assemblies and long-read data from a geographically diverse set of fifty *F. excelsior* samples, with particular focus on individuals showing low ADB susceptibility. We identified 362,965 structural variants (SVs) present in more than three individuals, including 174Mb of sequence absent from the linear reference genome (22% of the linear reference size). We demonstrated that failing to explicitly link SVs with gene sequences can lead to substantial overestimation of dispensable genes (those that vary in their presence/absence between individuals) due to variability in the annotation process. Controlling for this reduced the fraction of the genome estimated as dispensable from 35.9% to 8.7%, identifying 3,412 high-confidence dispensable genes, including 141 annotated with gene ontology terms associated with defence response. We used the pangenome to analyse existing genomic data from over 1,200 individuals to identify loci associated with reduced susceptibility to ADB. This revealed 220 single nucleotide polymorphisms (SNPs) showing allele frequency shifts between healthy and highly damaged pools of individuals that are broadly consistent across the mainly UK seed sources sampled, explicitly demonstrating the existence of a shared genetic component to low ADB susceptibility.

## Introduction

High quality reference genomes constructed from single individuals have been used to great effect to assess genetic diversity within species, allowing mapping of short reads from many individuals and discovery of genetic variants^1^. However, this approach provides limited insights into larger structural variants (SVs) between individuals and can lead to some miscalling of single nucleotide polymorphisms (SNPs)^2^. Pangenomes – reference genome data structures incorporating highly contiguous sequences from a representative set of individuals – have revealed the total sequence contained in SVs can exceed the average genome size of a species^3–6^. These SVs frequently encompass genes, with some studies identifying >50% of genes within a species as dispensable (i.e. not present in all individuals^5,7–9^). Pangenomes thus often contain genes that would be entirely missed if using a single reference genome approach^10,11^. Indeed, novel disease resistance genes are frequently identified in pangenomes^12–14^ including in trees^15–17^, suggesting pangenome approaches may be particularly valuable when seeking to detect loci contributing to pathogen resistance. Analysis of existing data using pangenome references has uncovered the basis of missing heritability in some systems^18^ and has enhanced our understanding of the genetic basis of a variety of traits, often adding complementary information to that uncovered by analyses using a linear reference genome^5,19–21^. Thus, to better understand genetic variability within a species and its impact on traits, pangenome references incorporating diverse samples are required^10,22^.

*Fraxinus excelsior* (European ash), a keystone tree species within Europe, is under severe threat from ash dieback disease (ADB) caused by the invasive non-native fungus *Hymenoscyphus fraxineus*^23,24^. Populations of ash are undergoing substantial and rapid mortality^25–27^, with high associated management costs and biodiversity loss^28–30^, attracting substantial public concern^31–34^. A small fraction of trees (as low as 0.5%^35^) remain healthy under *H. fraxineus* inoculum pressure, with quantitative resistance to ADB having a narrow-sense heritability of approximately 0.4^35–39^. Understanding the genetic basis of this trait is important for ascertaining the feasibility of producing new individuals with lower susceptibility^40–44^. Stocks et al. (2019)^42^ identified 192 SNPs significantly associated with low ADB susceptibility when using a p-value threshold of 1 x 10^−13^ in mainly UK populations by mapping short reads to a single reference genome. Studies in other European populations did not find SNPs strongly associated with low ADB susceptibility^43,44^, with one study indicating many of the most significant from Stocks et al.^42^ were at very low frequencies in Austrian populations^45^, raising the possibility of diverse mechanisms of low susceptibility between populations in different parts of the species’ range^42–47^. A pangenome reference for *F. excelsior* that captures variation from different genetic backgrounds would represent an invaluable resource to enable improved understanding of this critically important trait. A recently published “superpangenome” of the *Fraxinus* genus identifies interspecific SVs associated with salt tolerance, suggesting a role for these variants in adaptation to abiotic stressors over millions of years^48^. However, this study does not capture intraspecific SVs in *F. excelsior* that may contribute to low ADB susceptibility.

Here we construct a chromosome level, phased genome assembly for European ash and a graph pangenome using long-read sequencing from a diverse sample of 50 individuals across Europe. This revealed 174Mb of additional sequence captured in SVs, an increase of 22% from the new linear reference genome size from a single individual. By explicitly associating putative dispensable genes with SVs, we identified 3,412 high confidence dispensable genes in *F. excelsior*, including 141 with gene ontology terms associated with defense response. Reanalysis of poolseq data (i.e. sequence data from a mixture of approximately equal amounts of DNA from multiple individuals) from over 1,200 individuals^42^ revealed 220 SNPs with broadly consistent allele frequency shifts between unhealthy and unhealthy pairs of pools across a set of mainly UK seed sources, providing explicit evidence for a shared genetic component to low ADB susceptibility.

## Results

### A high quality linear reference genome for European ash

A phased linear reference genome was produced for a single *F. excelsior* individual, with total sizes of 793 and 787 Mb, N50s of 32.9 and 32.2 Mb, and L90s of 22 for the haplotype 1 (henceforth referred to as “BATG-1.0”) and haplotype 2 assemblies respectively (Supplementary Table 1). *Fraxinus excelsior* has 23 pairs of chromosomes^49^ – the top 23 scaffolds of BATG-1.0 and haplotype 2 contain 93.7% and 94.5% of the total sequence in each assembly, with the 24th scaffold for each of the haplotypes being 2.1% and 1.5% of the size of the 23rd scaffold. Three short scaffolds were identified as potential contaminants (with lengths between 14,626 and 35,091bp), representing 0.009% of the total assembly size – these were removed from downstream analysis. Telomeres were detected at both ends of 19 of the top 23 scaffolds in BATG-1.0 and 17 of the top 23 in haplotype 2. Together these results indicate the assemblies are pseudochromosomal. Unlike other recently published pseudochromosomal assemblies of *F. excelsior*^48,50^, this assembly is not scaffolded using the *F. pennsylvanica* genome assembly^51^. We also assembled and annotated the *F. excelsior* chloroplast genome; this was 155.6kb in length with 130 annotated genes.

98.2% and 98.3% of eudicot BUSCOs were completely assembled in the BATG-1.0 and haplotype 2 genomes; 21% of BUSCOs were duplicated, a similar proportion as in other *F. excelsior* assemblies (Supplementary Table 1) and consistent with the fact that there are several whole genome duplication events in the Oleaceae, the family to which *F. excelsior* belongs^48,52^. 98.3% of PacBio HiFi reads mapped to one or both of the haploid assemblies. The total assembly sizes of 787Mb and 793Mb are smaller than the 1C genome sizes of 877Mb and 840Mb estimated by flow cytometry in Sollars et al.^53^ (2017) and Siljak-Yakovlev et al. (2014)^54^ respectively – different estimates could result from methodological variation or inter-individual genome size differences. Analysing 27bp kmers from the PacBio reads using GenomeScope^55^ gave an estimated genome size range of 823-825.4Mb for this individual (Supplementary Figure 1) – other kmer lengths tested (see Methods) gave very similar size estimates. The coverage of read mapping to both phased assemblies did not show clear peaks of higher depth beyond the main homozygous peak, which would be indicative of collapsed sequences (Supplementary Figure 2). This suggests the remaining difference between estimated genome sizes and assembly length may be due to highly repetitive regions such as centromeres and ribosomal DNA that are challenging to assemble. The assembly sizes are similar to the size of unpublished pseudochromosomal assemblies of *F. excelsior* that do not use *F. pennsylvanica* for scaffolding (i.e. FRAX_001_PL and daFraExce3.hap1.1; see Supplementary Table 1). The LTR Assembly Index (LAI) was 14.85 and 14.62 for BATG-1.0 and haplotype 2 respectively, with 5.35-5.44 intact LTRs/Mb (comparable to other recent *Fraxinus* assemblies), with the top 23 scaffolds in each having average LAIs of 20.74 and 20.62 respectively (Supplementary Table 1) – a score of >10 indicates a reference standard genome^56^.

65.6% of BATG-1.0 consists of repeats. LTR elements occupy 33.0% of the sequence, most of which comprises Ty1/Copia (15.3% of genome sequence) or Gypsy/DIRS1 (15.9% of genome sequence) families, with DNA transposons covering 6.2% of the sequence. Unclassified repeats covered 19.9% of the sequence. The total repetitive fraction is similar to the estimated repeat content of the assembly of another *F. excelsior* individual (69.12%) and assemblies for some other *Fraxinus* species such as *F. pennsylvanica* from the *Fraxinus* superpangenome^48^, but is substantially higher than a separate estimate for *F. pennsylvanica* (48.4%^51^). Olive, also a member of the Oleaceae, has a lower repeat content than *F. excelsior* (56.2%) ^57^, although tomato (a fellow asterid with a similar 1C genome size) is more similar (65%-72%)^58,59^.

Annotation of BATG-1.0 was performed using BRAKER3^60^. Ten runs of BRAKER3 were performed with short-read RNA-seq data and protein orthologue information, and 10 runs with long-read RNA-seq data and protein orthologues. These identified 31,926 genes annotated in every run, with a further 1,736 genes annotated in every run of the short-read annotation only and 1,246 genes annotated in every run of the long-read annotation only. The median gene length of genes exclusively identified by long-reads was 43% longer than those identified only by short reads (Supplementary Figure 3), although different underlying tissue types were used to generate the long-read and short-read RNA-seq data, making these differences challenging to explicitly link to the length of RNA-seq input data. A further 3,075 genes were inconsistently annotated, in either the short– or long-read runs. To reduce the chances of erroneously identifying genes as dispensable in the pangenome, we initially retained all these genes, giving a total of 37,983. This is a similar number to that reported in the BATG-0.5 assembly (38,949)^53^ and the *F. pennsylvanica* assembly (35,470)^51^ but lower than the 43,392 genes reported in the recent Fexc_ONT_UoY *F. excelsior* assembly^50^ and substantially lower than the *Fraxinus* assemblies used in the *Fraxinus* superpangenome (48,389-53,538)^48^ including their *F. excelsior* assembly (50,327). These latter two *F. excelsior* assemblies have more genes annotated than are plausibly missing from BATG-1.0 due to being dispensable (see pangenome annotation below), suggesting differences are likely due to different annotation approaches rather than intraspecific variation in gene content. Using OrthoFinder^61^ to identify orthologues from BATG-1.0 and available annotations for *F. excelsior* assemblies (BATG-0.5, Fexc_ONT_UoY) as well as *F. pennsylvanica*, *Olea europaea*, *Solanum lycopersicum*, *Erythranthe guttata*, *Sesamum indicum* and *Salvia miltiorrhiza* (Supplementary Table 2), 0.6% of BATG-1.0 genes lacked an orthologue in another assembly, compared with 1.1% in BATG-0.5 and 7.3% in Fexc_ONT_UoY. 98.7% of eudicot BUSCO proteins were found in the total BATG-1.0 protein set, suggesting the annotation captures the vast majority of genes in the assembly.

### Structural variant calling and pangenome construction

Low susceptibility to ADB is likely a highly polygenic trait^42,47^, so we prioritised sampling a larger number of individuals to construct the pangenome instead of focusing on generating fewer highly contiguous assemblies, even if this restricts the types of SV that can be analysed. Oxford Nanopore Technologies (ONT) data was generated for 49 individuals from a variety of European provenances, as well as for the reference individual (Figure 1A, Supplementary Table 3). PCA of SNPs called from the 49 individuals revealed little geographic structure (Figure 1B), as was also found to be the case when a similar set of provenances were analysed in earlier work^53^.

**Figure 1.**
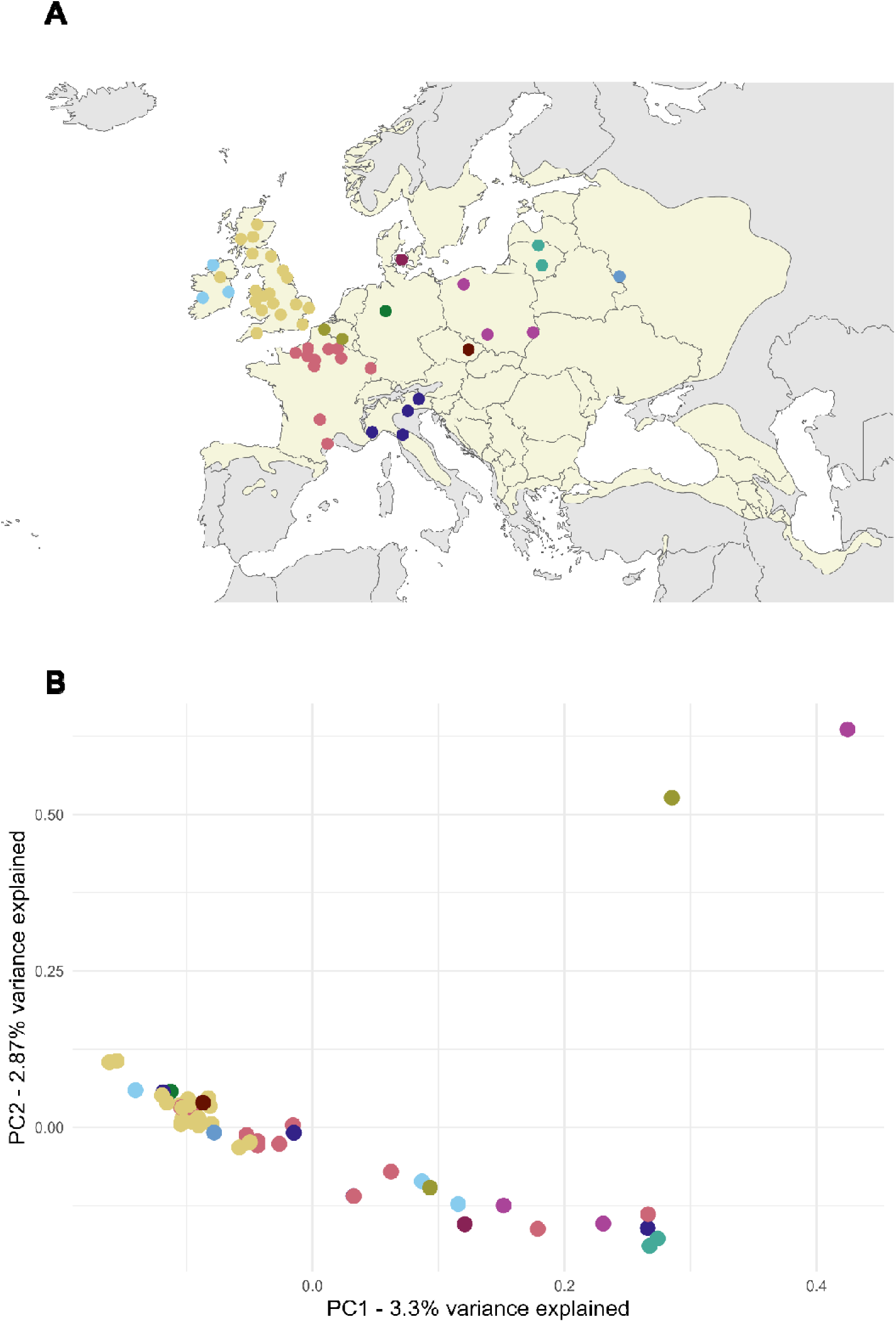
– Geographic origins and genetic structure of samples. A) Map showing location of origin for individuals used to construct the pangenome; see Methods and Supplementary Table 1 for full details. Points represent individuals sampled or approximate sample locations for seedlots from which individuals were grown. Light yellow areas represent the native range of *F. excelsior* using shapefiles from Caudullo et al.^119^. B) Principal components analysis of single nucleotide polymorphism (SNP) calls from long-read sequencing data from individuals sampled. First two principal components are plotted, with the percentage of variation explained by each principal component shown on the axis labels. In Figures A and B, individuals are coloured by country of origin (see Supplementary Table 1).

Heterozygous structural variants called between the two high quality haploid genomes of the reference individual were used to assess the accuracy of calling SVs using lower coverage ONT data from the same individual with a variety of methods. These included mapping the ONT reads to the reference haploid genome and calling SVs using Sniffles2^62^ and cuteSV^63^, *de novo* assembling the ONT reads with Flye^64^, Shasta^65^ or nextDenovo^66^ and calling SVs from contigs mapped against the reference using svim-asm^67^, and the combination of read mapping and assembly mapping approaches (see Methods). This analysis indicated that the read mapping approaches had higher recall of SVs detected using the high quality haploid genomes, while assembly mapping approaches had higher precision, and a combination of read mapping and mapping of the assembly produced by Shasta gave the highest recall (84.4%) and F1 score (77.9%), with a precision of 72.2% (Extended Data Figure 1). For insertions, deletions and tandem duplications, this method also gave the highest F1; for inversions, using svim-asm from a Flye *de novo* assembly gave a higher F1, although the overall precision/recall for inversion and tandem repeats was low (Supplementary Figure 4).The mapping and/or Shasta assembly based approach was therefore used to call SVs for all 50 samples. Within each sample, a median of 41.6% of SVs called with cuteSV and Sniffles2 were also called with svim-asm, and a median of 72.6% of SV called with svim-asm were also called with cuteSV and Sniffles2 (Supplementary Figure 5).

Merging SVs across all individuals identified a total of 737,995 SVs (Figure 2A). Merging samples sequentially resulted in fewer new SVs discovered in the fiftieth sample compared with the first, but the number newly detected did not plateau, indicating that more samples would be required to exhaustively sample SVs in *F. excelsior* (Figure 2B). It is also possible that sample-specific SVs represent sample-specific assembly or mapping errors – to attempt to control for this, only SVs present in more than three individuals were retained for further analysis (Figure 2A), although this may result in rare SVs being missed. This resulted in 362,965 SVs, of which 170,202 were insertions, 189,724 were deletions, 295 were inversions and 2,744 were tandem duplications (Figure 2A; Supplementary Figure 6). The total number of SVs represented 396Mb of variable sequence across the *F. excelsior* individuals, including 174Mb of insertion sequences not present in the linear reference genome (Figure 2C). 99.9% of SVs were on the 23 pseudochromosomal scaffolds. Insertions and deletions were generally more frequent towards the ends of chromosome arms (Figure 2D; Supplementary File 1). 69% of SVs overlapped with an annotated repeat in the reference genome; the distribution of base pairs overlapping different SV types closely reflected the makeup of the genome as a whole (i.e. mostly LTRs and DNA transposons), although rRNAs and tRNAs were relatively unlikely to overlap segregating SVs, likely due to their core role in the cell machinery (Supplementary Figure 7). The set of 362,965 SVs was used to construct a pangenome using vg^68^ which had 30,710,268 nodes and 31,069,469 edges.

**Figure 2.**
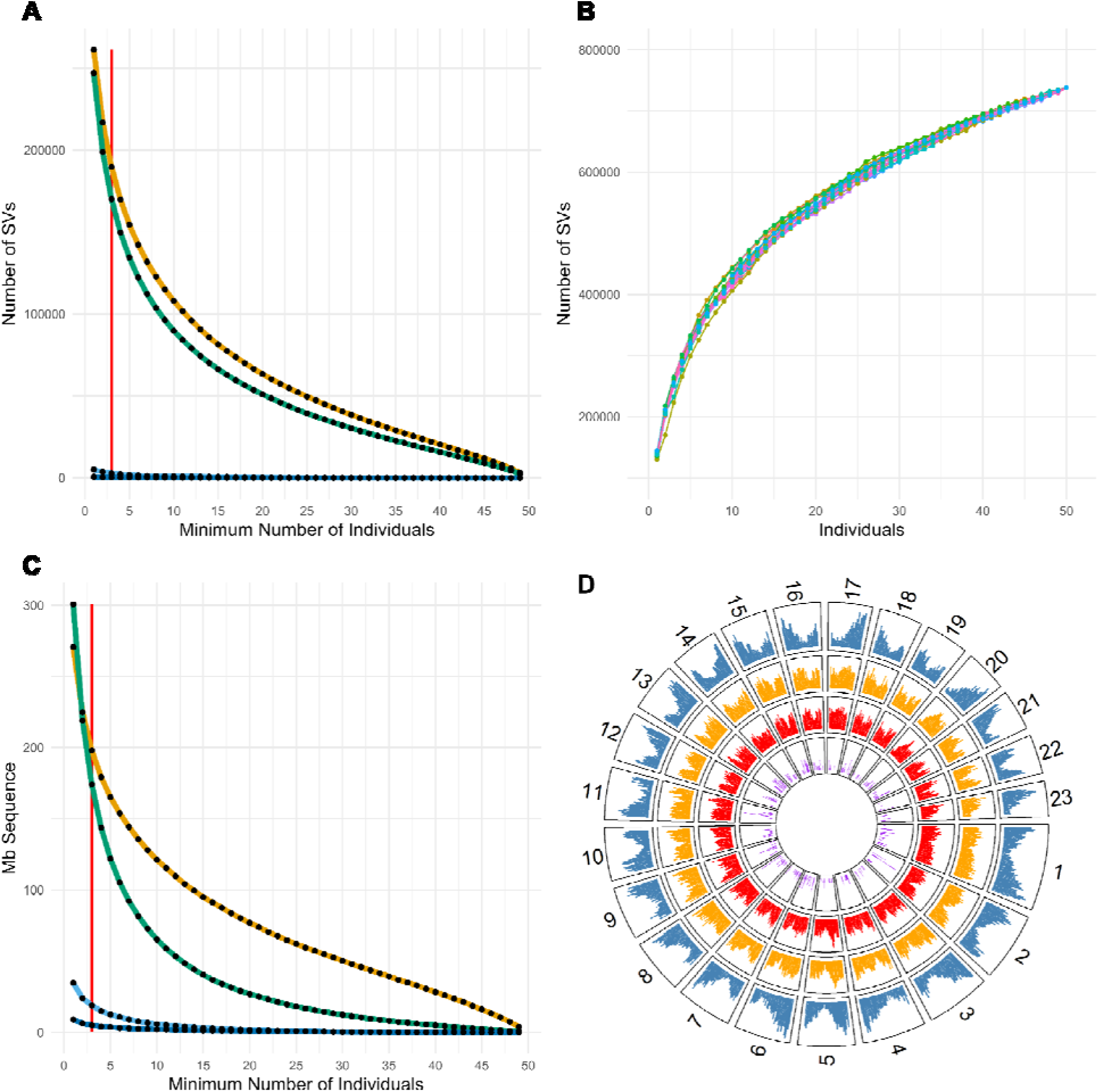
– Structural variant properties. A), C): The total number of SVs (A) and total quantity of sequence (C) of different SV classes, at different minimum numbers of individuals called from long-read data. Colour-coding indicates different types of SVs: insertions (green), deletions (orange), tandem duplications (light blue) and inversions (dark blue). A vertical red line at n = 3 indicates the cutoff for SVs being included in the pangenome. Note: to aid visualisation, separate plots for inversions and tandem repeats are available in Supplementary Figure 6. B) Saturation curves of sequentially merging SVs from different individuals. Each sequential series of SVs was generated by combining individuals in a random order, one at a time, with the total number of SVs in the group being displayed; separate series of random combinations are designated by colour, with points in the same series joined by a line. The X axis represents the number of individuals merged, the Y axis the total number of SVs in those individuals. D) Circular plot of feature densities per chromosome in 100kb windows. The 23 largest scaffolds are labelled 1-23 in descending order of size.Tracks, from outer to inner correspond to density of i) genes (blue), ii) insertions (orange), iii) deletions (red) and iv) inversions (purple). Larger plots for each individual chromosome, also including repeat and tandem duplication density, are presented in Supplementary File 1.

### Pangenome annotation: explicitly linking dispensable genes with SVs

To identify dispensable genes with an explicit link to SVs in the pangenome, a genome sequence was generated for each individual by using the SV genotype calls to modify the BATG-1.0 reference fasta (i.e. deleting deletions, inserting insertion sequence, etc). Genes were *de novo* annotated for each individual genome sequence and consolidated across individuals using OrthoFinder to identify a set of unique gene models. Of the 48,356 genes, 30,992 (64.1%) were present in all individuals including the reference (Figure 3). Of the 10,411 genes absent from the BATG-1.0 genome (i.e. putatively undiscovered dispensable genes), 6,009 did not overlap with an SV in any individual. For those that did overlap with SVs, only 1,474 did so with an F1 score (a measure of how consistently a gene presence or absence was associated with a SV, see Methods) of 0.8 or above (Figure 3, Extended Data Figure 2). As in each case the non-SV sequence is identical to the reference sequence, these putatively dispensable genes may be the result of stochasticity in the annotation process, or the result of SVs elsewhere in the genome altering the mapping of RNA-seq data and thereby reducing evidence to support the gene annotation. Similarly, of the 6,953 genes present in the reference assembly but absent from one or more of the other assemblies, 3,190 did not overlap with a SV in any individual. For those that did overlap with a SV, only 1,938 did so with an F1 score of 0.8 or above (Figure 3, Extended Data Figure 2). Putatively dispensable genes with an F1 score of less than 0.8 were classified as invariant if present in the reference genome, and if absent were removed from the analysis. This brought the total number of genes in the pangenome to 39,419, of which 3,412 were dispensable (8.7%% of the total). This is a substantial reduction in dispensable genes compared with the estimated 35.9% of genes without filtering for those consistently overlapping with SVs (Figure 3). Putatively dispensable genes without a consistent SV association (i.e. those excluded from our final annotation) could be affected by the non-comprehensive RNA-seq dataset (only 72% of individuals in the pangenome were represented in the RNA-seq data) or the particular annotation pipeline used. The final pangenome gene set included 99.0% complete eudicot BUSCOs – slightly more than the 98.7% captured in the BATG-1.0 protein sequences alone.

**Figure 3.**
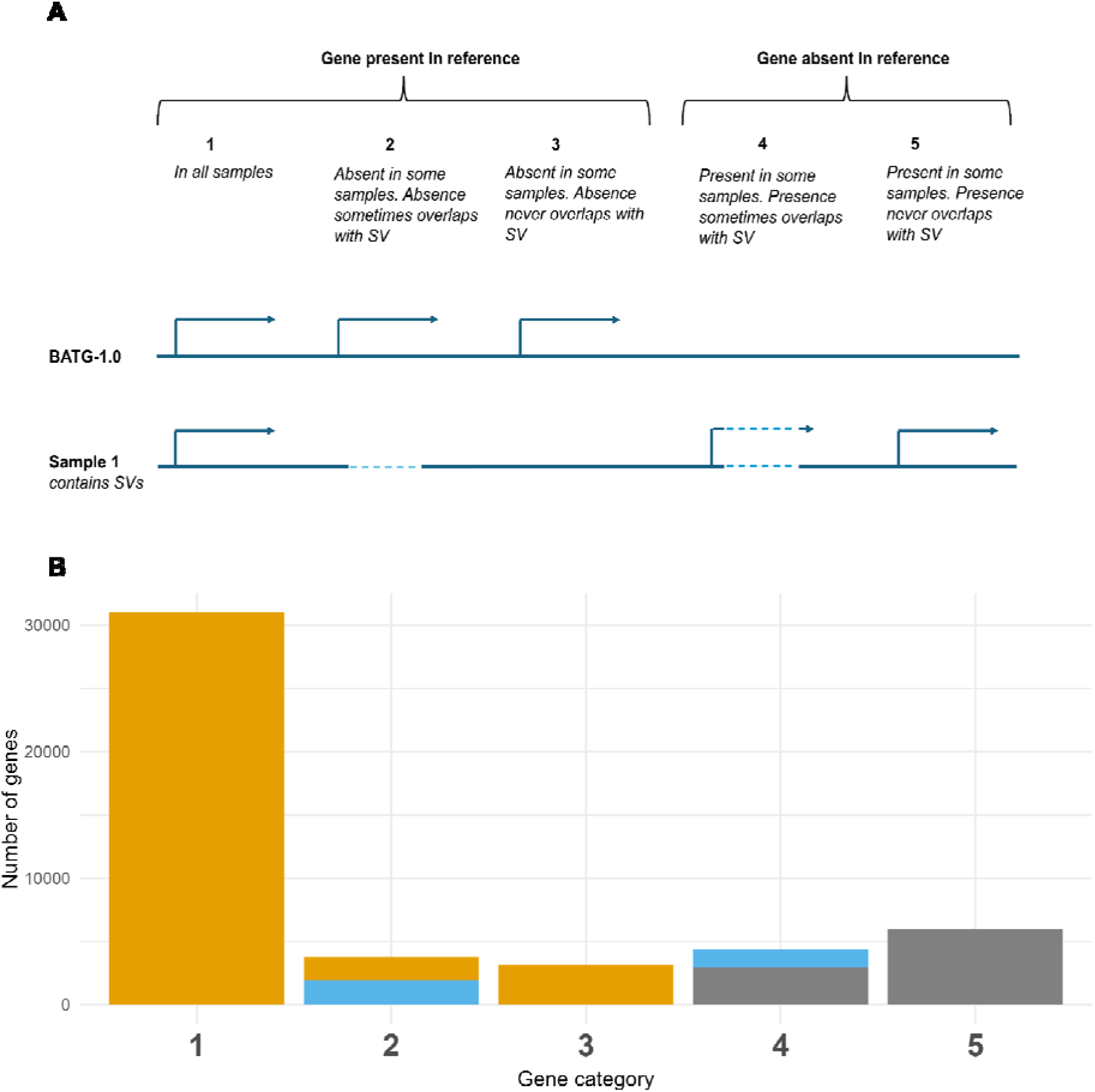
– Consistency of association of putatively dispensable genes with structural variants. A) Illustration of classification of genes in the pangenome based on their presence/absence, and how this corresponds with structural variants (SVs). Genome sequence is depicted as a blue solid line; SVs by light blue dashed lines (representing deleted sequence relative to the reference in this illustration). Genes are depicted by black arrows. For illustration, each gene in the reference sequence (top line) is homologous to the gene immediately below them in the example sample (bottom line). For those present in the BATG-1.0 reference, genes can be present in every other sample (1) or sometimes absent (2, 3). The absence of a gene can either coincide with an SV overlapping the gene in at least one sample (2), or with no overlap with an SV (3). For genes absent from the reference but present in at least one sample, this presence can either coincide with an overlapping SV in at least one sample (4), or never coincide with an overlapping SV (5). B) Barplot of the number of genes in each category in the pangenome, where numbers correspond to those in part A. Following reclassification of genes in each category as invariant (orange), dispensable (blue), or discarded from the analysis (grey) according to how consistently they overlap with SVs based on an F1 score. Genes in category 2 with an F1 score > 0.8 are classified as dispensable (blue); those with a score below that threshold are categorised as invariant. Genes in category 3 are never associated with an SV so are all categorised as invariant. Genes in category 4 with an F1 score > 0.8 are classified as dispensable, those with a score below that are discarded from the analysis. Genes in category 5 are never associated with an SV so are discarded from the analysis. In an approach where genes absent in some individuals are treated as dispensable, all the genes in categories 2-5 would be treated as dispensable genes.

Functional annotation indicated that high confidence dispensable genes overall had fewer annotations than indispensable genes; putatively dispensable genes that were filtered out also had a similar reduction in functional annotations compared with indispensable genes (Figure 4A). This may be a result of these gene subsets being less conserved across species and/or less well annotated. High-confidence dispensable genes were enriched for eight GO terms including “RNA-templated DNA biosynthetic process”, “retrotransposition” and “floral meristem growth” (Supplementary Table 4). Putatively dispensable genes that were filtered out were enriched for 28 terms including “defense response” and “regulation of auxin polar transport” that were not significant in the high-confidence dispensable genes (Supplementary Table 4). However, a direct comparison of the high confidence dispensable genes with the filtered genes provided little evidence of GO terms only enriched in the dispensable genes, or the filtered genes, differing meaningfully in frequency between these two sets (Figure 4B). The 3,412 dispensable genes include those with GO terms indicating a role in response to stress (355), developmental process (341) and defence response (141), with 26 genes linked to a function in “defense response to fungus” specifically (Supplementary Table 4). Dispensable genes with a “defense response” GO term belonged to a variety of protein superfamilies (Supplementary Table 5), including cytochrome P450s (three genes) and flagellin-sensing 2 (FLS2)-related serine/threonine protein kinases (two), with Nucleotide Binding Site, Leucine Rich Repeat (NBS-LLR) superfamily being the most frequently annotated (eight genes). Genes annotated for functions in leaf development (42) and leaf senescence (16) were also included within the dispensable set – variation in Spring leaf flushing and Autumn leaf senescence has been linked to low ADB susceptibility in some studies^38,44,69,70^.

**Figure 4.**
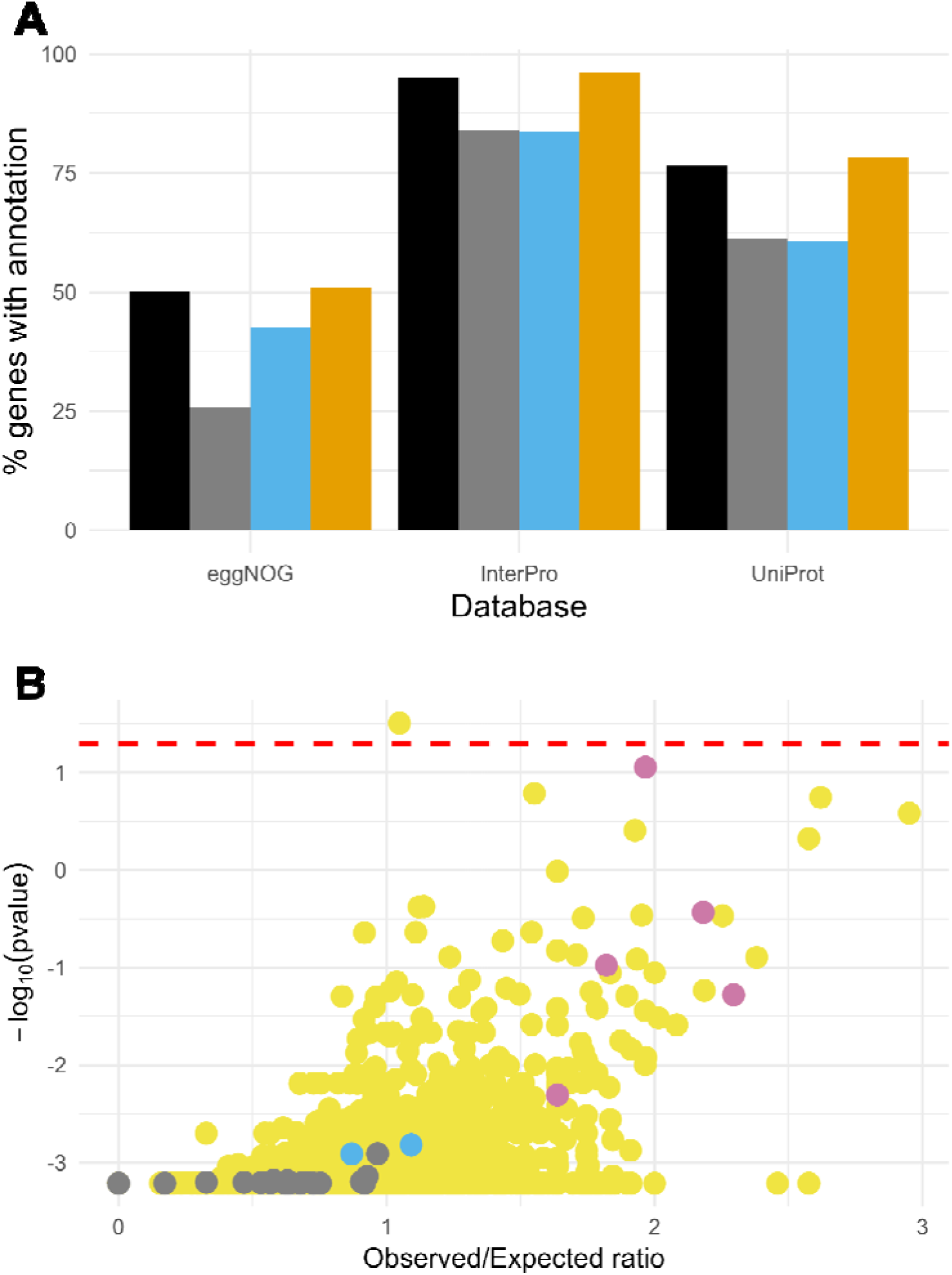
– Annotation of putatively dispensable genes. A) Barplots showing the percentage of genes annotated in the eggNOG, InterPro and UniProt databases, for genes included in the pangenome after filtering (black), genes removed during filtering (grey), included genes classified as dispensable (i.e. the high-confidence dispensable genes; light blue) and included genes classified as invariant (orange). B) Results of Gene Ontology (GO) enrichment analysis of the included dispensable genes, against a background of the included dispensable genes plus the putatively dispensable genes removed during filtering for not being consistently associated with an SV, with points coloured by significance in other GO enrichment analyses. Values along the x-axis represent the ratio of the observed number of genes in the included dispensable genes to the number of expected genes in the total background set. Y-axis values represent the –log_10_ transformed p-values from the GO enrichment test. The red dashed line indicates the significance threshold. Points are coloured according to whether those GO terms were enriched in GO enrichment tests for other sets of genes. Gray points represent GO terms identified as significantly enriched in the genes removed during filtering, against a background of excluded plus included genes in the pangenome. Blue points represent GO terms identified as significantly enriched in genes classified as variable after filtering, compared to a background of all genes included after filtering. Purple points represent GO terms identified as significantly enriched in both the previous comparisons described. The remaining GO terms are coloured in yellow.

### Read mapping, SNP calling and SV calling using the pangenome

We next tested whether using our pangenome reference improved read mapping for short-read sequenced individuals, compared to the BATG-1.0 linear reference sequence. While overall mapping to our pangenome produced little effect on the total number of short reads mapped in proper pairs (Extended Data Figure 3), we observed general improvements based on other metrics: this includes increases in mean mapping quality and alignment score (1.4% and 1.2% on average, respectively Extended Data Figure 3), a reduction in the total number of substitutions (3.0% on average, Extended Data Figure 3), and a large reduction in soft clipped sequence (33.5% on average, Extended Data Figure 3). Overall, this indicates the pangenome improves the accuracy of short-read read mapping. Mapped reads in each case were used to call SNPs – overall numbers of SNPs and quality metrics were similar using both approaches; only 5.6% and 3.4% of SNPs called in the reference-based and pangenome approaches respectively were not called with the other approach. The quality of SNPs that were not detected with both approaches appeared similar to the general background set of SNPs (Supplementary Figures 8,9; Supplementary Table 6). The median distance of unique SNPs from one approach to SNPs from the other approach was <30bp for both callsets, although the reference-based callset included some SNPs >1kb from the nearest SNP in the pangenome-based callsets (Supplementary Figure 9). This indicates that while the improved overall mapping to the pangenome results in some SNP-calling differences, the overall impact on SNP-based analysis is likely to be minimal compared to that associated with using a high-quality linear reference genome.

Examination of allelic depths of homozygous alternate, reference and heterozygous sites showed SVs and SNPs exhibit similar distributions of read support for reference and alternate alleles (Extended Data Figure 4). The minor allele frequency (MAF) distributions and deviation from Hardy-Weinburg equilibrium (HWE) for the two variant types were also similar (Extended Data Figure 5A, B). Patterns of linkage decay with distance were broadly similar in SNP-SNP, SV-SV and SNP-SV comparisons, with slightly faster linkage decay between SVs, and between SVs and SNPs, compared with between SNPs (Extended Data Figure 5C). Together, these suggest SNP and SV calls behave in a similar way for individually sequenced samples.

To investigate the accuracy of allele frequency estimations from short-read poolseq data, we constructed an *in silico* pool with an equivalent number of individuals and depth to the average *in vitro* pool from Stocks et al.^42^. Allele frequencies for SNPs were estimated using Popoolation2^71^. Similar software does not exist for estimating SV allele frequencies in poolseq data using a pangenome. Whilst vg^68^ does not explicitly support poolseq allele frequency estimation, we called SVs for the pools using vg and used the depths of reference vs. alternate alleles to estimate SV frequency in the pools. Correlation between the alternate allele frequencies from the individual based calling and estimates from the *in silico* pool was high for SNPs (slope = 0.94, R^2^ = 0.93). For SVs, whilst the overall correlation was also high (slope = 0.96, R^2^ = 0.85), there were a minority of SVs where the pool-based estimates differed substantially from the individual based calling (Extended Data Figure 6A, B). Poolseq data from Stocks et al.^42^ was mapped to the pangenome, and SNP and SV frequencies analysed. PCA recovered similar population structures from SV and SNP frequencies estimated using these pools (Extended Data Figure 6C, D) as has been found in other studies^68,72^. Together, these results suggest that allele frequencies of most SVs can be captured accurately by mapping poolseq data to a pangenome.

### GWAS with SVs and SNPs

We reanalysed poolseq data from Stocks et al. (2019)^42^ using the pangenome. In that study, for 13 seed sources, a pool of healthy trees (no visible signs of ADB damage) and a pool of unhealthy trees (for which the majority of individuals had a dead/partially dead main stem; 42 trees per pool on average) from field trials heavily affected by ADB were sequenced (see Methods for details). We performed a GWAS for low ADB susceptibility using SV allele frequencies estimated from this data, with a Cochran-Mantel Haenszel (CMH) test^73,74^. This test identifies associations of treatments and outcomes while taking into account stratification – it is a commonly used statistic for identifying associations from paired poolseq data^42,71,75–79^. This identified 531 significant SVs with a p-value cutoff of 1 x 10^−13^ (the same significance threshold as used for the CMH test in Stocks et al.^42^; Extended Data Figure 7A). However, many of these sites showed inconsistent allele frequency estimates from the *in silico* pools vs. individual genotyping as assessed in the above section (261 had a difference in estimated frequency of > 0.2; see Extended Data Figure 7B vs. Extended Data Figure 6B). Whilst some of the 531 significant SVs may be truly associated with low ADB susceptibility, many are likely false positives resulting from errors in the estimation of allele frequencies. These results indicate SV allele frequency estimates from poolseq data are insufficiently accurate for a reliable GWAS.

We next considered the estimated allele frequencies for SNPs called from the Stocks et al.^42^ poolseq data using our pangenome, and used these to perform a separate GWAS for low susceptibility to ADB using the CMH test. This displayed high p-value inflation (Supplementary Figure 10A). 866 SNPs were identified as significant after Bonferroni correction with = 0.05 (uncorrected p-value threshold = 1.52 x 10^−9^), and 404,503 with 5% false discovery rate using the Benjamini-Hochberg procedure. 92 SNPs had a p-value of less than 1 x 10^−13^ (the significance threshold used for the CMH test in Stocks et al.). Wiberg et al. (2017)^75^ demonstrated that heterogeneity in SNP allele frequency shifts among pairs of pools can be identified as a main effect, causing p-value inflation in the CMH test. Deviation from homogeneity, as measured either by a significant Woolf test^80^ p-value (p < 0.05: 53.4% of significant SNPs), or by an I^2^ statistic > 0.25 (73.2% of SNPs)^81^, was common, and was particularly pronounced for SNPs with the lowest CMH test p-values (Supplementary Figure 11). Indeed, many of the SNPs with the lowest p-values (less than 1 x 10^−13^) had high read depth variation between pools (Extended Data Figure 8), consistent with the presence of rare SVs distorting estimates of allele frequency (see Supplementary Note 1 for further discussion).

A quasibinomial generalized linear model (qGLM) test – a GLM for proportion/binomial data including an overdispersion parameter – has been shown to be better able to identify consistent allele frequency shifts across heterogeneous pairs of pools^75^. This analysis did not identify any significant sites, although substantial p-value deflation suggests this test may be overly conservative (Supplementary Figure 12A). Indeed, the lowest p-values displayed consistent allele frequency shifts, but with shifts in allele frequency between healthy and unhealthy pools similar to random subsets of SNPs (Supplementary Note 2, Supplementary Figure 12B-F). Different approaches to modelling the effect of provenance using GLMs either led to a similar lack of significant SNPs, or substantial p-value inflation (Supplementary Note 2, Supplementary Figure 13). This indicates limited evidence for SNPs behaving in a highly consistent manner across pools. We attempted to explicitly assess heterogeneity in allele frequency changes across pairs of pools using the Fixation index (*F_ST_*) or Fisher’s exact test, and considering sites where multiple pairs of pools showed a strongly supported allele frequency shift. However, this proved ineffective for identifying sites shared by subsets of pools (with minimal overlap found even between biological replicates from Stocks et al.^42^), likely due to the inherent noise in individual allele frequency estimates from poolseq data (see Supplementary Note 3).

Despite inflated p-values, the 866 SNPs passing CMH test Bonferroni correction displayed large allele frequency shifts compared to the genome-wide average, also displaying a consistent direction of change in most pairs of pools (Figure 5), suggesting they are likely enriched for SNPs associated with ADB resistance in the majority of seed sources analysed. 220 of these SNPs did not violate the CMH test assumption of homogeneity, or display high standard deviation in read depth potentially resulting from SVs not captured in the pangenome (see Methods), suggesting their low p-values are more likely to represent a true association with tree health status. 34 genes contained one or more of these SNPs, including two encoding pentatricopeptide-repeat containing proteins, ribosomal subunit proteins, and transcription and translation factors (Supplementary Table 5). A further 177 genes were within 10kb of these SNPs (Supplementary Table 5), of which 16 were dispensable. No GO terms were significantly enriched in either the 34 genes overlapping the SNPs, or the 211 genes within 10kb, after correction for multiple testing (Bonferroni correction with = 0.05; uncorrected p-value threshold of 1.53 x 10^−5^; Supplementary Table 4), with “immune response-activating cell surface receptor signaling pathway” and “brassinosteroid metabolic process” among those in the 10 most enriched terms. The set of 211 genes included nine with a “defence response” GO term annotation. These included i) a gene belonged to the NBS-LRR superfamily, ii) a dispensable gene homologous to an *Arabidopsis* globulin family protein, and iii) a phosphatidylinositol-specific phospholipase C homologue that directly overlapped the significant SNPs.

**Figure 5.**
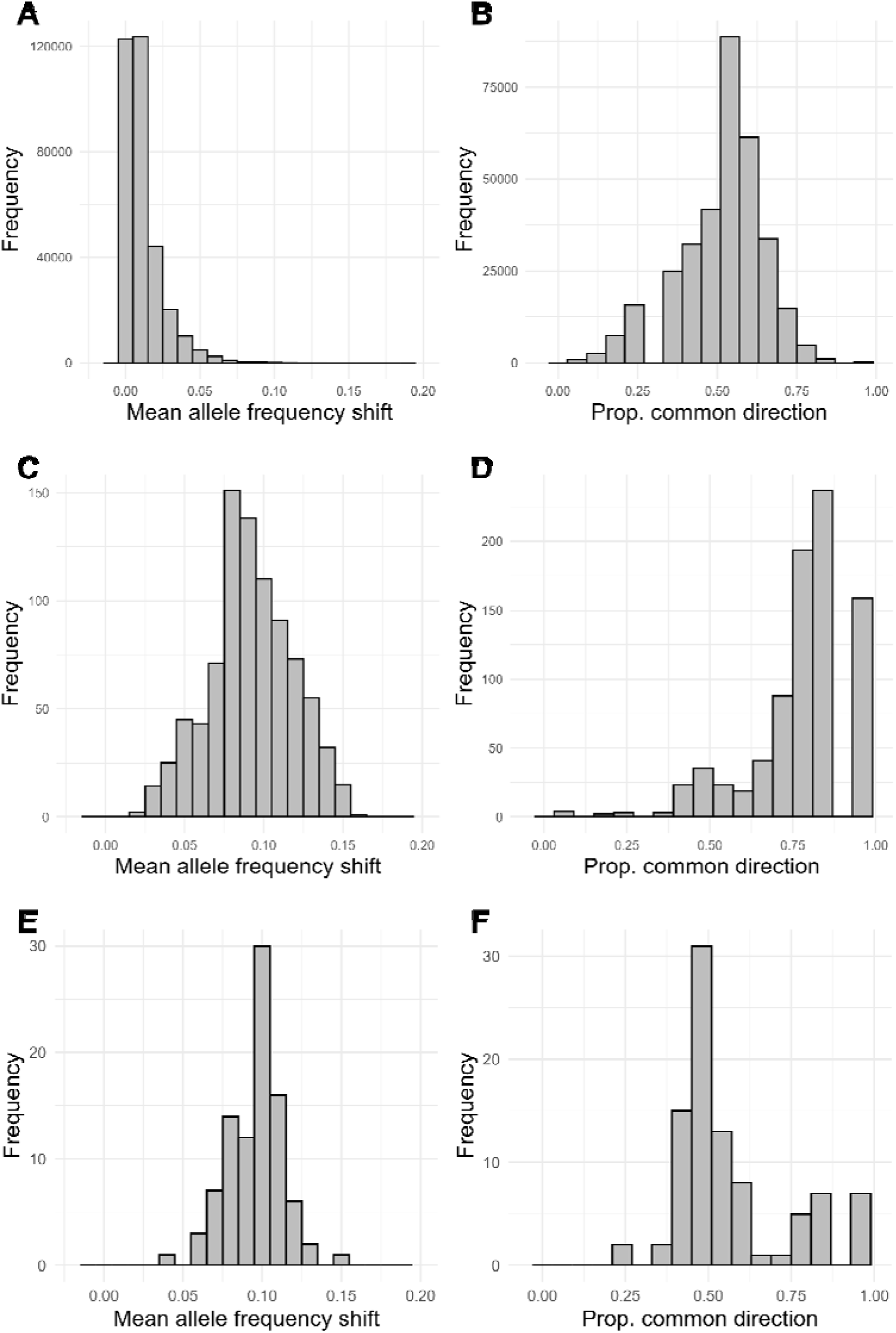
– Shifts in direction between healthy and unhealthy pools in candidate SNPs compared to background SNPs. For a random 1% of SNPs estimated from poolseq data (panels A and B), and SNPs with significant p-values from a Cochran-Mantel Haenszel CMH) test, with Bonferroni correction, LJ = 0.05, (panels C and D), and SNPs with a CMH p-value < 1 x 10^−13^ (panels E and F), i) the mean absolute value of the allele frequency shift between healthy and unhealthy pools (A, C and E), and ii) the proportion of all pairs of pools shifting in the most common direction (B, D and F).

## Discussion

Here we present a new chromosome-level linear reference genome (BATG-1.0) and a pangenome for European ash from a geographically diverse sample of individuals, enriched for those with low susceptibility to ADB. The pangenome of 50 individuals captures 362,965 SVs, representing 396Mb of variable sequence, including 174Mb that is absent from the linear reference genome (equivalent to approximately 22% of its length). This is broadly similar to other recent pangenome studies, although there is substantial variability across these^3,15,82–95^ (Extended Data Figure 9A, 9B; Supplementary Table 7). Our analyses likely underestimate the total SV diversity in *F. excelsior*, as each additional individual contributed novel SVs to the collection. A comparison of SVs called between the chromosome-level haploid assemblies, and ONT data for the same individual, indicates our SV calling approach has a precision of 72.2% (although a recall of 84.4%) compared with using chromosome-level phased assemblies, suggesting some SVs included in the pangenome may be false positives. However, it should be noted that i) as heterozygous sites only have half the read support of homozygous alternate sites it is likely the latter can be detected with higher recall and precision, and ii) the fact that we only included SVs called in more than three samples in the pangenome potentially reduces the number of false positives, although at the cost of excluding rare SVs. Our results highlight the substantial diversity of SVs in European ash, consistent with high SNP genetic diversity^53^, but also underline that more extensive sampling is required to fully capture segregating SVs.

We identified 8.7% of genes in the pangenome as dispensable; this is substantially lower than most other plant pangenomes, which typically report 20-60% of genes or gene families as being variable^3,15,82–93,95^ (Extended Data Figure 9C, Supplementary Table 7). As noted above, our pangenome only contains SVs called in more than three individuals, so potentially misses the rarest dispensable genes. However, most of the difference in the proportion of genes that are dispensable is explained by our filtering of genes not reliably associated with SVs; without this filter, a more typical 35.9% of genes in the pangenome are classified as dispensable (Extended Data Figure 9C, Supplementary Table 7). Despite leading to a dramatic reduction in dispensable genes, we believe this filter is justified as variability in the annotation process has previously been found to underlie putatively dispensable gene sets^92,95^. Indeed, previous pangenome studies have identified “core” genes as being more highly expressed^7,84,87,90,96,97^. This relationship could be due to these genes performing critical biological functions, but also could be the result of more highly expressed genes having more RNA-seq data evidence and being more accurately annotated as a result. Functional groups typically found as enriched among dispensable genes in other studies, such as defense and auxin response^3,7,84,92,94,98^ were enriched in our initial set of putatively dispensable genes, but not following our filtering process. It should however be stressed that there was no clear functional distinction between the final set of dispensable genes and those removed during filtering – a smaller set of genes will inherently have lower power to reliably detect enrichment in GO terms. More comprehensive RNA-seq sampling from tissues/conditions across individuals, and manual curation of repetitive elements, may lead to more genes being annotated – including as dispensable – in future studies. We should also note that we aimed to identify genes specifically within our characterised SVs; in studies where *de novo* assemblies are explicitly incorporated (e.g. all-against-all pangenomes and comparative genomics more broadly) specific testing of DNA sequence differences underlying presence/absence of genes should be performed, as has been suggested elsewhere^99^. Our results emphasise the importance of confirming that putatively dispensable genes are driven by underlying sequence differences, to distinguish them from possible non-biological annotation variability. The genes we identified as dispensable with high confidence include 141 with homology to defence-related genes, one of which was within 10kb of the 220 SNPs associated with low ADB susceptibility, emphasising the importance of pangenomes in understanding the mechanistic basis of disease resistance traits.

Using a pangenome reference allows improved short-read mapping for *F. excelsior*, as has been found for other species^11^, and we were able to successfully call individual SNPs and SVs from individually sequenced samples. However, there was not a clear improvement in SNP calling compared to using the BATG-1.0 linear reference genome. SV genotyping from short-reads is generally more challenging than SNPs^100–102^, and we are not aware of previous studies calling allele frequencies of poolseq data mapped to a pangenome graph. Our results indicate poolseq data can be used to estimate allele frequencies accurately for most SVs (Extended Data Figure 6), but the approach used generates more discrepancies with individual based genotyping than is the case for SNPs. SVs showing these discrepancies were particularly likely to be identified as significant in a GWAS using a CMH test, resulting in many likely false positives. We were unable to identify the cause of these discrepancies (see https://github.com/vgteam/vg/issues/4443). We should emphasise that vg is not designed for analysing poolseq data; the development of dedicated software for estimating SV allele frequencies from poolseq data, equivalent to those available for SNPs, is required in order to make full use of poolseq data in the pangenome era. Nevertheless, some of the SVs identified as significant from the CMH test are potentially involved in low ADB susceptibility; future studies using individually genotyped data could be used to confirm this.

Despite broadly accurate SNP allele frequency estimates from poolseq mapped to a pangenome, our reanalysis of data from Stocks et al (2019)^42^ highlighted known challenges of analysing poolseq data^103^. The commonly used CMH test can assign very low p-values to heterogenous allele frequency shifts among pairs of pools^75^, and uncharacterised SVs can distort allele frequency estimations. A combination of these sources of error meant that despite mapping to a pangenome and using relatively stringent depth filtering, a likely rare, large/high copy number SV produced the lowest p-values in our dataset using this test (Supplementary Note 1). We find a subset of 220 SNPs with significant p-values from the CMH test that do not violate its assumptions of heterogeneity, or display unusual patterns of depth, and shift in frequency between unhealthy and healthy pools in a broadly consistent manner across most seed source zones (Figure 5). We would note the inflation of p-values from the CMH test makes the threshold of inclusion in this group (passing Bonferroni correction with = 0.05) somewhat arbitrary. Nevertheless, these results indicate a shared genetic component to low susceptibility exists across a broad range of provenances. It should be noted that more densely sampled regions in the Stocks et al.^42^ dataset (e.g. Great Britain) may make SNPs consistently shifting in these regions more likely to be identified. Whilst no GO terms were significantly enriched in genes near these SNPs, the “brassinosteroid metabolic process” term was highly ranked – brassinosteroid biosynthesis genes were enriched near SNPs predictive of autumn leaf yellowing in a previous study, a trait which has been associated with low ADB susceptibility^44^. Variation in the frequency of trees with low susceptibility exists between provenances^35,36^, suggesting there may be further geographically restricted genetic variation associated with low ADB susceptibility, although we are not able to accurately characterise this in our dataset. Higher coverage per pool, or individual based sequencing, would allow more accurate characterisation of variants restricted to subsets of provenances.

In Stocks et al (2019)^42^, the consistency of allele frequency shifts, and the potentially confounding effect of SVs, were not explicitly examined. Indeed, the maximum per-pool depth for sites considered was 37.5X the average coverage, rather than the more stringent 2.5× in the present study, increasing the potential for SVs to impact allele frequency estimates. It is likely therefore that the 192 SNPs with p-values lower than 10^−13^ in Stocks et al. are similarly, or more severely, enriched for problematic sites, as those in our study. This may explain the lack of correlation of CMH p-values with estimated effect sizes in genomic prediction (GP) of ADB damage, and the fact that around two thirds of their top 192 GWAS hits are ranked outside the top 2,500 SNPs by GP effect size (Stocks et al. 2019). It may also explain a lack of shared genes between Doonan et al. (2025) and Stocks et al.^44^, although we should note that ADB health scores represent a complex phenotype underpinned by several mechanisms, which may reduce the chances of identifying similar underpinning regions across studies. Nevertheless, a GP model trained on the top 10,000 GWAS loci from Stocks et al. was able to predict tree health status with an accuracy of 67%^42^, despite likely including many loci violating the assumptions of the CMH test. This is consistent with our results which suggest that despite containing false positives, SNPs with significant CMH test p-values after Bonferroni correction are nevertheless enriched with loci that display broadly consistent allele frequency shifts between healthy and unhealthy pools.

Overall, our newly developed pangenome captures a diverse set of SVs from across the geographic range of European ash, including a large quantity of previously uncharacterised sequence and dispensable genes, making it an essential resource for full investigation of the role of both SNPs and SVs in a breadth of ecological traits for this important tree species. After carefully accounting for spurious signals in our GWAS, we detect signals of a shared component of low susceptibility to ADB across UK provenances. Explicitly identifying variants with consistent effects across a range of seed sources may be useful in developing widely-applicable genomic prediction models for low ADB susceptibility, and ultimately informing breeding programs to restore this threatened species.

## Methods

### Reference assembly

To construct a phased, chromosome-level assembly, we sampled a potted graft growing at Royal Botanic Gardens, Kew. The graft was created by Forest Research in 2017 using a rootstock and a scion from *F. excelsior* individuals of unknown provenance. Leaf tissue from this graft was flash frozen in liquid nitrogen in May 2022 and sent to Cantata Bio, CA, USA (formerly Dovetail Genomics) on dry ice, where DNA extractions, library preparation, sequencing and genome assembly was performed as outlined below.

DNA was extracted by Cantata Bio using an in-house CTAB protocol and quantified using Qubit 2.0 Fluorometer (Life Technologies, Carlsbad, CA, USA). The SMRTbell Express Template Prep Kit 2.0 (PacBio, Menlo Park, CA, USA) using the manufacturer recommended protocol was used to construct a PacBio SMRTbell library (∼20kb) for PacBio Sequel. The library was bound to polymerase using the Sequel II Binding Kit 2.0 (PacBio) and loaded onto PacBio Sequel II). Sequencing was performed on PacBio Sequel II 8M SMRT cells, resulting in 66Gb data (approximately 79X coverage based on the prime 1C genome size estimate for *F. excelsior* of 840Mb^54,104^. For the Omni-C libraries, chromatin was fixed in place with formaldehyde in the nucleus and then extracted. Fixed chromatin was digested with DNAse I, chromatin ends were repaired and ligated to a biotinylated bridge adapter followed by proximity ligation of adapter containing ends. After proximity ligation, crosslinks were reversed and the DNA purified. Purified DNA was treated to remove biotin that was not internal to ligated fragments.

Sequencing libraries were generated using NEBNext Ultra enzymes and Illumina-compatible adapters. Biotin-containing fragments were isolated using streptavidin beads before PCR enrichment of each library. The library was sequenced on an Illumina HiSeqX platform to produce approximately 30X sequence coverage in 150bp paired-end reads. The combined Illumina and PacBio data was used to produce an initial assembly with hifiasm v0.15.4-r347^105^ using default parameters. The *de novo* assembly and Dovetail OmniC library reads were used as input data for HiRise^106^ to produce a draft assembly. OmniC library reads were then aligned to the draft input assembly using BWA^107^ and reads with MQ < 50 removed. The mapping information was analyzed using HiRise to produce a likelihood model for genomic distance between read pairs, and the model was used to identify and break putative misjoins, and to score and make joins.

After receiving the assemblies and raw data from Cantata Bio, the haploid genome assemblies were assessed using QUAST v5.2^108^ and BUSCO v5.5.4 using the eudicots_odb10 database^109^. To assess potential contaminants, scaffolds from the most contiguous of the two haploid genomes (BATG-1.0) were searched against the NCBI nt database (downloaded 21/02/23) using blast+ v2.11.0^110^ with settings –max_target_seqs 10 –max_hsps 1 –evalue 1e-25. To assess coverage, the PacBio data was mapped to the haploid genomes using minimap2 v2.5^111^. The combined data was analysed using blobtools v4.0.7^112^. To assess LTR completeness, LTRharvest from genome-tools v1.6.6^113^ and LTR_finder v1.07^114^ were run with default parameters, other than specifying a 85% similarity threshold, length range 100-5000bp, starting/ending motif TG/CA and maximum distance of LTR starting positions of 20,000. LAI was calculated using LTR_retreiver v3.0.4^56^. To estimate genome size using kmers, LVgs v1.00^115^ was run on the PacBio fastq files, with kmer sizes between 17 and 77bp in increments of 10, and results analysed using Genomescope2 v2.1.0^55^

The PacBio data was used to *de novo* assemble a chloroplast genome for *F. excelsior* using Oatk v1.0^116^ using HMMER v.3.1b2 (hmmer.org) with parameter settings –c 30 –k 1001 and a chloroplast gene profile database provided by OatK (embryophyta_pltd). Annotation was performed using GeSeq v2.03^117^ using default settings.

### Long-read RNA extraction and sequencing

Long-read RNA seq data from leaf and cambium tissue was generated to annotate the new genome assembly from the reference individual. Leaf material was collected from the same grafted tree as sampled for the reference assembly (see above) in May 2022 and flash frozen in liquid nitrogen. For cambium tissue, twigs were sampled in January 2023 and the layer of tissue immediately below the bark was isolated using a fresh razor blade before flash freezing in liquid nitrogen. Tissue was stored in a –80 freezer until RNA extraction.

For long-read RNA extraction, a modified version of the Qiagen RNeasy Plant Mini Kit standard protocol was used (Supplementary Note 2). In brief, 25-30mg tissue was used with the standard kit protocol (including the optional DNase step) and eluted in 50µl RNase-free water. 450µl buffer RLT was added to the RNA, and the RNeasy kit protocol was repeated (without the DNase step). RNA was quantified using a Nanodrop 2000 and a Promega QuantiFluor® RNA System Kit using a Quantus fluorometer. Samples were shipped to the Deepseq facility at the University of Nottingham and fragment length determined with an Agilent Tapestation using an RNA Screentape kit.

cDNA libraries were prepared using an Oxford Nanopore PCR-cDNA Sequencing Kit (SQK-PCS109), loaded onto R9.4.1 flowcells and sequenced on a Gridion X5 Mk1 – this was performed once for the cambium library and twice for the leaf library. Runs were performed with high-accuracy basecalling (running on MinKNOW software version 22.12.5). All data reported relates to reads that passed a base calling quality filter (PHRED score >9).

### Sampling for pangenome construction

In sampling individuals for the ash pangenome, we aimed to i) sample a broad range of European populations, with ii) a particular focus on UK diversity by sampling individuals representing different NSZ in the UK^118^ (see https://assets.publishing.service.gov.uk/government/uploads/system/uploads/attachment_data/file/797909/Regions_of_provenance_and_seed_zone_map.pdf). With this approach we aimed to capture a global set of variants from across the species range, whilst increasing our chances of capturing SVs segregating in the mainly UK populations for which short-read data is available from Stocks et al.^42^.

In total, 50 samples were used for long-read sequencing. To maximise the chances of identifying SVs segregating in the poolseq data from Stocks et al.^42^ one individual from each of the 13 NSZ used in the Forest Research (FR) mass screening field trial at North Burlingham, Norfolk, UK (Site 16)^42^ was sequenced, with the exception of NSZ-204 where two were sequenced, as individually sequenced data for this provenance were available from Stocks et al.^42^ The North Burlingham site was planted in 2013 with two-year-old saplings that were grown for non-research purposes in commercial nurseries from seed sources from 15 different provenances. The site contained four complete replicate plots for each seed source, with 5,000 trees per ha (a spacing of 1 m× 2 m). At the time of sampling, the site had high *H. fraxineus* inoculum pressure from naturally established infections, and ADB damage was severe throughout. In 2022 (when trees were eleven years old) individuals were sampled that appeared relatively healthy within each plot, and scored between 3 and 5 on the damage scale of Pliura et al.^37^ (where 3 is a tree with moderate damage from ADB, 4 a tree with slight damage and 5 a tree with no visible signs of damage) to increase the chances of capturing alleles for low ADB susceptibility. Three wild trees of unknown age from seed zones in the UK not covered by the FR trial populations (NSZ-305, NSZ-406, NSZ-402) were also included to further capture UK diversity. To increase the likelihood of including more of the range-wide diversity of structural variation in *F. excelsior*, one individual from each of 29 mostly mainland European provenances was sampled from the Realising Ash’s Potential (RAP) trial (established in 2005), as well as two from provenances in the FRAXIGEN trial (established in 2004), both at Paradise Wood, Earth Trust, Oxfordshire. Finally, leaf material from the individual used for the linear reference genome was also included. Sample information is outlined in Supplementary Table 3, including approximate coordinates of origin where available. A map based on these coordinates is shown in Figure 1A, overlaid with a shapefile of the *F. excelsior* native range from Caudullo et al.^119^. We note our sampling does not comprehensively cover the geographic range of *F. excelsior* – in particular, the far south western, south eastern European, west Asian, and far northern areas of the range are unrepresented in our samples (See Fig. 1A).

Sampling for long-read DNA sequencing was performed in May 2022. For samples collected in the field, intact stems with leaves were excised from individual trees using secateurs or a pole pruner, wrapped in damp tissue paper and sealed in airtight plastic bags before returning to RBG Kew for storage at 4°C. Leaf tissue was removed from the stems and frozen within 48 hours of collection by immersing in dry ice, and then stored at –80°C prior to DNA extraction.

### High molecular weight DNA extraction, and RNA extraction, from pangenome samples

For high molecular weight DNA extraction, a modified CTAB protocol was used^50^ (see Supplementary Note 6 for full protocol). In brief, 300-400mg tissue was disrupted in liquid nitrogen and added to 8ml Carlson buffer with 20µl beta-mercaptoethanol and 40µl Qiagen Proteinase K. This was incubated at 65°C for 1 hour, before adding 16µl Qiagen Rnase A and incubated for a further 15 minutes. 1 volume of SEVAG solution was added, the tube inverted, and centrifuged for 10mins at 4°C before the aqueous phase was removed. This step was then repeated. 0.7 volumes of isopropanol was added and the tube incubated at 4°C for 15mins. The sample was centrifuged at 4°C for 30mins to precipitate the DNA and resuspended in Qiagen buffer AE before being redissolved in 5ml Qiagen Buffer G2 for 15 minutes. A Qiagen Genomic Tip 100/G Kit was used to purify the DNA; the “Isolation of Genomic DNA from Blood, Cultured Cells, Tissue, Yeast or Bacteria using Genomic-tips” protocol from the kit handbook was used, with the modification of using 10ml rather than 7.5ml buffer QC. DNA was eluted in 100µl buffer 1x TE and left to resuspend at room temperature and then at 4°C for up to 3 and 4 days respectively. DNA was quantified using a Quantus Flurometer with QuantiFlour dsDNA kit, and a Nanodrop 2000. Integrity was measured using an Agilent Tapestation with a Genomic DNA kit, and the DNA shipped to the University of York on dry ice to prevent fragmentation during transit, where library preparation was performed as described below.

Short fragment elimination was performed using a Circulomics SRE Standard Kit. Library preparation was performed using the Oxford Nanopore Technologies (ONT) SQKNBD114.24 (Protocol version NBE_9169_v114_revD_15Sep2022-promethion.pdf) library preparation kit, with the following modifications: i) Reaction volumes were proportionately scaled up to 60μl total for end repair and A-tailing, and incubation times increased to 30 minutes each at 20°C and 65°C, ii) Reaction volumes were doubled for the barcode adapter ligation step, which was performed at room temperature for 1 hour. Barcode-ligated samples were bead cleaned separately before pooling at appropriate volumes prior to adapter ligation. Native barcodes were ligated and reads demultiplexed by barcode following the run. Libraries were loaded onto PromethION flowcells, runs commenced on a Gridion with live super accuracy base calling (running on MinKNOW software version 22.10.5) with barcode demultiplexing and adapter trimming. All data reported relates to reads that passed a base calling quality filter (PHRED score >10).

For 35/50 samples with remaining leaf tissue after DNA extraction, RNA from frozen material was extracted for short read sequencing (outlined in Supplementary Table 3). For one additional sample (PG5), fresh leaf tissue was collected in May 2024 and stored in Thermofisher RNALater, and stored at 4°C until RNA extraction in November 2025. The same RNA extraction protocol was used as previously described (Supplementary Note 5). Samples were sent to the Centre for Genomic Research (CGR), Liverpool. Analysis using an Agilent Tapestation did not reveal any evidence of genomic DNA contamination. cDNA libraries were prepared using a New England Biolabs NEBNext polyA selection and Ultra II Directional RNA library preparation kit, and sequenced on half a lane on an Illumina NovaSeq X Plus using 25B chemistry, with paired-end 15-bp reads. Raw data was trimmed by CGR using Cutadapt^120^ v4.5 with option –O 3. Further trimming was performed using Trimmomatic^121^ with settings LEADING:7 TRAILING:7 ILLUMINACLIP:2:30:7 MINLEN:70, resulting in an average of 63 million read pairs per sample after trimming (Supplementary Table 3).

### Long-read DNA data analysis

The simplex ONT DNA sequence data generated was trimmed, firstly using Trimmomatic^121^ with settings LEADING:7 TRAILING:7, followed by Nanofilt v2.8.0^122^ with minimum average read quality of 7 and length 1000. Quality was inspected and summary statistics generated using Nanoplot v1.4.10^123^, FastQC v0.11.9^124^ and multiQC v1.14^125^.

Reads were mapped to BATG-1.0 using minimap2 v2.5^111^, with PCR duplicates and reads with mapping quality (MAPQ) <20 removed using samtools v1.9^107^. Mean depth after filtering was 22.7X (Supplementary Table 3).

To inspect population structure of these samples, SNPs were called individually for all samples other than the reference genome individual using Clair3 v.1.0.5^126^ with model r941_prom_sup_g5014. Gvcfs were merged using glnexus v1.4.1-0-g68e25e5n^127^. Genotypes with DP < 8 or > 60 were set to missing using bcftools v1.16^128^, and sites with >10% missing data or MAF < 0.05 were excluded. Plink2 v2.00a3.3LM^129^ was used to identify unlinked SNPs (using setting –indep-pairwise 50 10 0.1) and calculate a PCA using default settings.

Read-based SV calling was performed using the long-read data generated for all 50 samples using Sniffles2 v2.0.7^62^ and cuteSV v2.0.3^63^, two tools that rank highly across several comparisons of read-based SV calling software^130–137^. Both tools were run with a minimum read support of 5, and cuteSV with the additional parameters – max_cluster_bias_INS 100 –max_cluster_bias_DEL 100 –diff_ratio_merging_INS 0.3 – diff_ratio_merging_DEL 0.3. Breakends/translocations (BND) or sites which were genotyped as homozygous reference were removed.

To complement the read-based SV calling, *de novo* assemblies were produced from the ONT read data. *De novo* assembly from trimmed reads was performed using shasta v0.11.1^65^ with the Nanopore-May2022 config file, Flye v2.9.2-b1786^64^ with default settings, and nextDenovo v.2.5.0^66^ with an estimated genome size of 850Mb. Assembly contiguity was assessed using QUAST. SVs were called using these assemblies by mapping them to BATG-1.0 using minimap2 in asm10 mode. SVs were called with svim-asm v.1.02^67^ using alignments with a minimum MAPQ of 20 and minimum SV size of 50. Haplotype 2 of the linear reference genome was also mapped to BATG-1.0, and SVs called using the above method. SV calls between the two pseudochromosomal haploid genomes were compared to SV calls from the ONT data from the same individual using the various approaches, with SV sets merged using SURVIVOR v1.0.7^138^ using settings 200 2 1 1 1 50 for the intersection of sets and 200 1 1 1 1 50 for the union. A 200bp merge distance yielded similar numbers of SVs called with Sniffles2 and cuteSV to values between 10 and 300bp yielded similar numbers of SVs, with larger or smaller values resulting in fewer SVs. Assuming the SV calls between the two haploid linear reference genomes are a “truth” set, precision, recall and F1 were calculated for the various SV calling methods from the lower coverage ONT data. Based on these comparisons (see Results), SVs for each individual were included if they were called in at least one of i) cuteSV and Sniffles2 ii) svim-asm from mapping the shasta assembly to BATG-1.0.

These SVs were merged across all 50 individuals using SURVIVOR with settings 200 2 1 1 1 50. SVs that overlapped with Ns in the reference were removed. SVs that were called in more than 3/50 individuals were identified using bcftools v1.16 and used for downstream analysis. The overlap between these SVs and the repeat annotation was assessed using the bedtools v2.28.0^139^ window function with parameter –w 1. Consensus sequences for insertions and deletions were searched against the ncbi nt database using blast+ with parameters –max_hsps 1 –evalute 1e^−25^, with the resulting data loaded into blobtools. Based on analysis with blobtools, four sequences were identified as putative contaminants and the associated SVs removed from the vcf. The resulting vcf, as well as the BATG-1.0 fasta was used to construct a pangenome using the vg v1.64.0^68^ construct command with option –v –S –a.

### Linear reference and pangenome annotation

Repeats were identified using Repeatmodeler v2.0.1^140^ in LTRStruct mode. The outputs of this were combined with the custom repeat library used previously to mask the BATG-0.5 assembly^53^, and used to mask the genome with Repeatmasker v4.1.2^141^ with the parameters –wublast –nolow. Telomere sequences were identified using Teloscope v0.0.9^142^ with default settings using the canonical plant telomeric repeat CCCTAAA. To annotate the linear reference genome, short-read RNA-seq data from leaf, root, cambium and flower tissue that had been used to annotate the BATG-0.5 assembly^53^, was downloaded from the European Nucleotide Archive (ENA), as well as RNA seq data from Emerald Ash Borer infested and control phloem tissue^143^, and ADB infested and control petiole tissue^144^ (Supplementary Table 8).Together with the short-read leaf RNA-seq data for the BATG-1.0 individual generated as part of this study, these data were mapped mapped to BATG-1.0 using STAR v2.7.10b with default parameters^145^. The long-read RNA-seq data generated in this study was trimmed firstly using Trimmomatic v0.39^121^ with settings LEADING:7 TRAILING:7, followed by Nanofilt v2.8.0^122^ with minimum average read quality of 7 and minimum length 100. The trimmed data was then mapped to the BATG-1.0 reference genome using minimap2 v2.5^111^. We acknowledge that this RNA-seq data does not provide comprehensive coverage of tissues/timepoints/conditions in the species, and that some genes may therefore be missed. Variability in the gene annotation process could lead to false positives during the identification of variable genes^16,146^, so we repeated the annotation process multiple times and combined the results. BRAKER3^60^ was run 10 times on the softmasked genome using the bam files from the short read data, and protein orthologue data from Viridiplantae from orthodb v11^147^. A version of BRAKER3 capable of using long-reads (singularity build braker3_lr.sif docker://teambraker/braker3:devel) was also run 10 times using the bam files from the long read data and the same protein orthologue set as used with the short read data. To combine the outputs of the runs, the amino acid sequences of the longest transcript per gene (minimum length 30 amino acids) for each run were clustered into orthogroups using Orthofinder v2.5.5^61^. Within each orthogroup, groups of sequences with non-overlapping positions within the BATG-1.0 assembly (likely representing recent duplicates) were separated using the bedtools intersect utility. For each of these groups, the gene model with the longest transcript from the set of 20 BRAKER3 runs was chosen. In some cases, OrthoFinder split genes with largely overlapping coding sequences into separate groups. In cases where two genes with a coding sequence on the same strand overlapped by >50% for one of the genes, only the gene producing the longest protein was retained (for further discussion on clustering see Supplementary Note 7). The features for the selected gene models were combined into a single bed file. The amino acid sequences from the final set of transcripts was used as the input for Orthofinder, along with protein sequences for BATG-0.5^53^, *Fraxinus pennsylvanica*^51^, *Olea europaea* subsp. *europaea*^148^, *Solanum lycopersicum*^149^, *Erythranthe guttata*^150^, *Sesamum indicum*^151^ and *Salvia miltiorrhiza*^152^ (Supplementary Table 2).

To annotate the pangenome, sequences and locations of structural variants were used to modify the sequence of the BATG-1.0 fasta, which was then annotated. As most SVs were defined from read mapping to BATG-1.0 rather than from the individual *de novo* assemblies, this approach maximises the opportunity for novel genes/gene deletions to be explicitly linked to underlying SVs. The steps were as follows (see Extended Data Figure 10 for an explanatory diagram): for each individual, a non-overlapping set of SVs detected in that individual was extracted from the combined vcf of SVs used to build the pangenome. Where SVs overlapped, insertions were retained in priority to other types of SVs, and longer SVs were retained in priority to shorter ones, as these may be more likely to capture coding sequences absent in BATG-1.0 This process resulted in 2,394 deletions, 96 insertions and 21 inversions being left unrepresented in any of the samples (0.7% of SVs). Tandem duplicates (0.5% of SVs), which were not included in the graph pangenome, were also excluded. Based on these sets of SVs, sequence was then inserted, deleted, or inverted in the reference fasta (the SV sequence in the combined vcf of SVs used to build the pangenome was used, so the same sequence was used to represent a given SV in different individuals) to produce a transformed fasta for each individual in turn. The transformed fasta was then annotated for each individual. Repeats were annotated and softmasked as described above. Long and short read RNA-seq data from the reference individual were mapped to each transformed fasta using the same methods as for the reference sequence. For the 35 non-reference individuals where short-read RNA-seq data was generated as part of this study, leaf RNA-seq data from each individual was also mapped to the corresponding transformed fasta and used in downstream annotation. A single run of the short-read BRAKER3 protocol, and a single run of the long-read BRAKER3 protocol was performed as detailed above. For these two runs for each individual, the Orthofinder approach as outlined previously was used to consolidate the outputs of BRAKER3 into a single set of proteins. Orthofinder was then run again for each set of proteins across the 50 individuals, as well as the proteins from the consensus annotation of BATG-1.0. Non-overlapping groups of sequences were separated using bedtools as before. For each gene, bedtools intersect was used to identify SVs that overlapped the gene sequence. Novel genes not present in the BATG-1.0 genome should overlap with an SV, as should deletions of genes present in the reference genome. For each novel gene that overlapped a SV at least once (i.e. in at least one sample, including the reference individual sequenced with long reads), an F1 score was calculated to identify the strength of the association between the new gene and the SV: 2 × N_Gene_SV / (2 × N_Gene_SV + N_Gene_NSV + N_NGene_SV) where N_Gene_SV is the number of samples with the gene and the SV, N_Gene_NSV is the number of samples with the gene but without the SV, and N_NGene_SV is the number of samples without the gene but with the SV. For novel genes overlapping more than one SV, the SV with the highest F1 score was retained as potentially causal for the novel gene. An equivalent calculation was performed for genes present in the BATG-1.0 genome but absent in at least one sample, where an SV overlapped with the annotation coordinates for the gene in the reference. To account for genes potentially being associated with multiple SVs, we also calculated an F1 score of gene presence/absence with all overlapping SVs – if this scored higher than any individual SV, this score was taken forward. Genes with an F1 score of 0.8 or above were retained in the final pangenome annotation as being dispensable; for those with an F1 score of < 0.8, those absent in the BATG-1.0 genome were removed, and those present in the BATG-1.0 genome were classified as invariant. Genes that did not overlap an SV in any sample were classified as invariant if they were annotated in BATG-1.0 and were absent in some other samples, and discarded if they were absent in BATG-1.0 but present in some other samples. To assess the impact of OrthoFinder potentially oversplitting genes, in a separate analysis, orthogroups containing at least one pair of genes with a coding sequence on the same strand overlapping by more than 50% were merged, and SV/Gene F1 scores recalculated. The results for this were qualitatively similar to the unmerged dataset, so the latter was used subsequently – for more discussion see Supplementary Note 7.

The complete set of putatively dispensable genes (i.e. before removing those with an F1 score <0.8) was functionally annotated. Protein sequences were searched against Araport 11 representative gene peptides (downloaded 14/09/2022), the NCBI SR database (downloaded 20/08/24), and UniProt/SwissProt database (downloaded 24/07/24) using DIAMOND v2.1.9.163^153^ in blastp mode. Interproscan v5.69.101.0^154^ with default settings was used to search for protein domains. GO terms were assigned using eggNOG-mapper v2.1.12^155^ by using DIAMOND blastp to search the eggNOG database. GO enrichment was performed using topGO v2.62.0^156^ using annot = annFun.gene2GO, nodeSize = 10 and the exact Fisher test with Bonferroni correction. OrthoFinder was run with the complete set of potential pangenome protein sequences using the same species and parameters described above.

### SNP and SV calling from short read data from individuals and pools

To test the impact of using the pangenome on read mapping accuracy, short-read data from 42 individuals separately whole genome sequenced in Stocks et al.^42^ (this number selected to reflect the average number of individuals in a pool in that study, see generation of *in silico* pools below) was mapped using vg giraffe^68^ to either i) the full pangenome or ii) the BATG-1.0 genome converted into pangenome format using vg construct. The resulting gam files were converted to bam files in the BATG-1.0 coordinate system using vg surject. Mapping statistics were calculated using samtools. For each mapping approach, SNPs were called using bcftools v1.19 call and calls merged across individuals. Genotypes with DP <5 or >50 were set to missing, then sites with >10% missing individuals or a MAF of <0.05 were removed.

For SV calling from these individuals, vg pack was used with parameter –Q 5, and SVs called using vg call with the vcf used to construct the pangenome as a reference. For SNP calling, bams were produced using vg surject. SNPs were then called using bcftools v1.19 call. SNPs and SVs were merged across individuals, and calls with DP < 5 or >50 were set to missing, then sites with >10% missing individuals or a MAF of <0.05 were removed. To produce similar numbers of SNPs and SVs for linkage analysis, 1% of SNPs were randomly sampled using vcflib v1.0.3^157^ function vcfrandomsample. SNP and SV data were combined using bcftools v1.19 concat option. Pairwise linkage between markers was calculated using plink v1.9.-170906^158^, with parameters –ld-window-kb 1000 and –ld-window 99999. Linkage decay with distance between SVs alone, between SNPs and SVs, and between SNPs alone were analysed separately.

To identify the accuracy of SNP and SV calling from poolseq data using the pangenome, read files from the 42 individually sequenced samples (which each contained approximately the same number of reads, equating to approximately 22X 1C genome coverage) were combined, and 10% of read pairs randomly subsampled with seqkit v2.8.0^159^. This corresponds to approximately 2.2X coverage per individual, the average per-individual coverage of the poolseq data in Stocks et al.^42^. SNPs and SVs were called from this *in silico* pool using the approaches above, but with sites being set to missing if they had a DP < 50 or DP > 200. Allele frequencies for SNPs/SVs were calculated as the number of reads supporting the alternate allele divided by the total number of reads. For the merged individually called SNPs/SVs, the correlation between the alternate allele frequency in the individually called samples and the estimated allele frequency for the *in silico* pool was tested with a linear model in R v4.3.1^160^.

### GWAS for low susceptibility to ash dieback from poolseq data

A poolseq dataset from Stocks et al.^42^ was used to perform a GWAS for low susceptibility to ADB. In brief, this data was generated as follows: leaf samples were taken from four FR mass screening trial sites in England (including the North Burlingham site where individuals were sampled for constructing the pangenome), from six year old ash trees from a total of 12 different seed sources (10 from UK NSZ, one from Ireland [IRL-DON] and one from Germany [DEU]), as well as a group from a breeding seedling orchard (sourced in the UK) planted by the Future Trees Trust (FTT-SO). For each of the 13 sources, a pool of healthy trees (no visible damage) and a pool of unhealthy trees (main stem dead, or partially dead) – 42 trees per pool on average – were sequenced with the Illumina HiSeq X platform in 150bp paired-end reads. Tree health was assessed based on a tree health score designed to assess the overall level of damage from ADB – it should be noted that low susceptibility to ADB is a complex trait, which may not be associated with long-term tree health, and may be underpinned by multiple underlying mechanisms. Two seed sources had a second pair of pools, produced as biological replicates, for a total of 30 pools (excluding one technical replicate). See Stocks et al.^42^ for full details. The trimmed, paired short read data from ENA (accession ERA1732352, Submitted Files) was used.

Here, these reads were mapped to the pangenome and read depths for the reference or alternate paths were extracted from the vcf for each pool to produce estimates of SV allele frequencies, as described in the previous section. The SV allele frequencies were then merged across pools. For SNPs, the output of vg giraffe was converted to linear reference coordinates using vg surject, with PCR duplicates and reads with MAPQ < 20 removed using samtools v1.9^161^. SNPs were called from the resulting bam files using samtools mpileup with –d 8000 and the mpileup2sync function from Popoolation2^71^ v1.201 with –min-qual 20 to produce a .sync file. Sites were removed if there was a third most frequent allele with a total depth higher than 15. The frequencies of major alleles for i) SVs and ii) a random 1% of SNPs were used to perform PCA using the R prcomp function.

To identify SV and SNP allele frequency changes associated with ADB damage, Cochran-Mantel Haenszel (CMH) tests were performed using the cmh-test.pl function from Popoolation2, with parameters min_count 15, min_coverage 40 and max_coverage 200. For sites with a significant result (p < 1 x 10^−13^), the Woolf test was used from the vcd R package^162^ to test for significant deviation from homogeneity across groups using the alternate/reference read counts for each pool. To quantify heterogeneity in allele frequency shifts, Cochran’s Q^73^ and I^281^ were calculated.

Cochran’s Q was calculated as 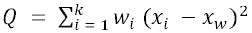 from Cochran (1954)^73^, with x_l_ being the estimated effect size from pool i, x_w_ the weighted mean effect size estimate across all k pairs of pools, and W_l_ being the inverse of the variance estimated for each pair. For the effect sizes, we used natural logarithm transformed of individual odds ratios [i.e. ln(ad/bc)], where a = alt reads in healthy, b = ref reads in healthy, c = alt reads in unhealthy, d = ref reads in unhealthy) as recommended in Woolf (1955)^80^ and performed in other studies^163^. The variance in this measure was estimated as 1/a + 1/b + 1/c + 1/d, as specified in Woolf (1955)^80^. The natural logarithm transformed overall CMH test odds ratio is used as x. These values were used to calculate Q. I^2^ was calculated from these as I^2^ = (Q – df) / Q, where df is the degrees of freedom; in this case, the number of pairs of pools minus one. For SNPs with p < 0.05 from the CMH test after Bonferroni correction, SNPs with a standard deviation of depth of >15, indicative of potentially uncaptured SVs that may interfere with allele frequency estimation (see Results, Supplementary Note 1), were removed, along with SNPs that showed deviation from homogenous odds ratios (violating an assumption of the CMH test) – namely, SNPs with either an I^2^ value of <0.25, or a Woolf test p-value of <0.05. 220 of the SNPs classified as significant after Bonferroni correction remained after this filtering. Genes from the pangenome that overlapped these SNPs, or were within 10kb, were identified using bedtools. For those with annotated GO terms, enrichment was performed as described above for dispensable vs. indispensable genes.

An alternative statistical test for GWAS, a quasibinomial generalized linear model (qGLM), was run using the poolFreqDiffTest_QGLM.py script from the PoolFreqDiff repository^75^, using parameters –mincnt 15 –minc 40 –maxc 200 –n 45. Different methods of accounting for the effect of provenance were investigated, see Supplementary Note 2 for further details. Fixation index (F_st_) and Fisher’s exact test were also calculated in sliding windows for pairs of pools – see Supplementary Note 3 for further details.

## Acknowledgements

We would like to thank i) Genomics Laboratory, Bioscience Technology Facility, University of York, ii) Deep Seq: Next Generation Sequencing Facility, University of Nottingham, iii) Cantata Bio LLC, California for their contribution to sequencing and genome assembly, iv) Earth Trust and other landowners for facilitating access for sampling. We would also like to thank Mike Charters and Owen Blake at RBG Kew, Domen Finzgar and Liz Richardson at Forest Research, Jo Clark at Future Trees Trust, and Paul Hill at the Earth Trust for assistance with sample acquisition, and Laszlo Csiba and the Plant Health and Adaptation team at Kew for valuable advice on this work. We are grateful to Andrea Harper and Sara Franco Ortega for early access to their DNA extraction protocol. This research utilised Queen Mary’s Apocrita HPC facility, supported by QMUL Research-IT (http://doi.org/10.5281/zenodo.438045). This work was funded by the UK Government Department for Environment Food & Rural Affairs through a Centre for Forest Protection grant (no. CFP2208) to LJK, DPW and RW. The results of this project LL2317 were obtained with the financial contribution of the Ministry of Education, Youth and Sports as part of the targeted support of the ERC CZ program, through the contribution of LY. DG and LJK received support from the Natural Environment Research Council [grant number NE/X015351/1 to LJK]. Figure 1 “contains, or is based on, information supplied by the Forestry Commission. © Crown copyright and database right [2009] Ordnance Survey [100021242]”

## Data availability

Source Data are provided with this paper. Sequencing data is available on NCBI under BioProjects PRJNA906474 (long-read DNA sequence data), PRJNA942609 (long-read RNA sequencing data) and PRJNA1390666 (short-read RNA sequencing data). Annotation files for the BATG-1.0 genome and pangenome will be made available on Zenodo.

## Code availability

Code needed to reproduce the analysis will be made available at https://github.com/danielwood1992/Ash_pangenome.

## Extended Data

**Extended Data Figure 1.**
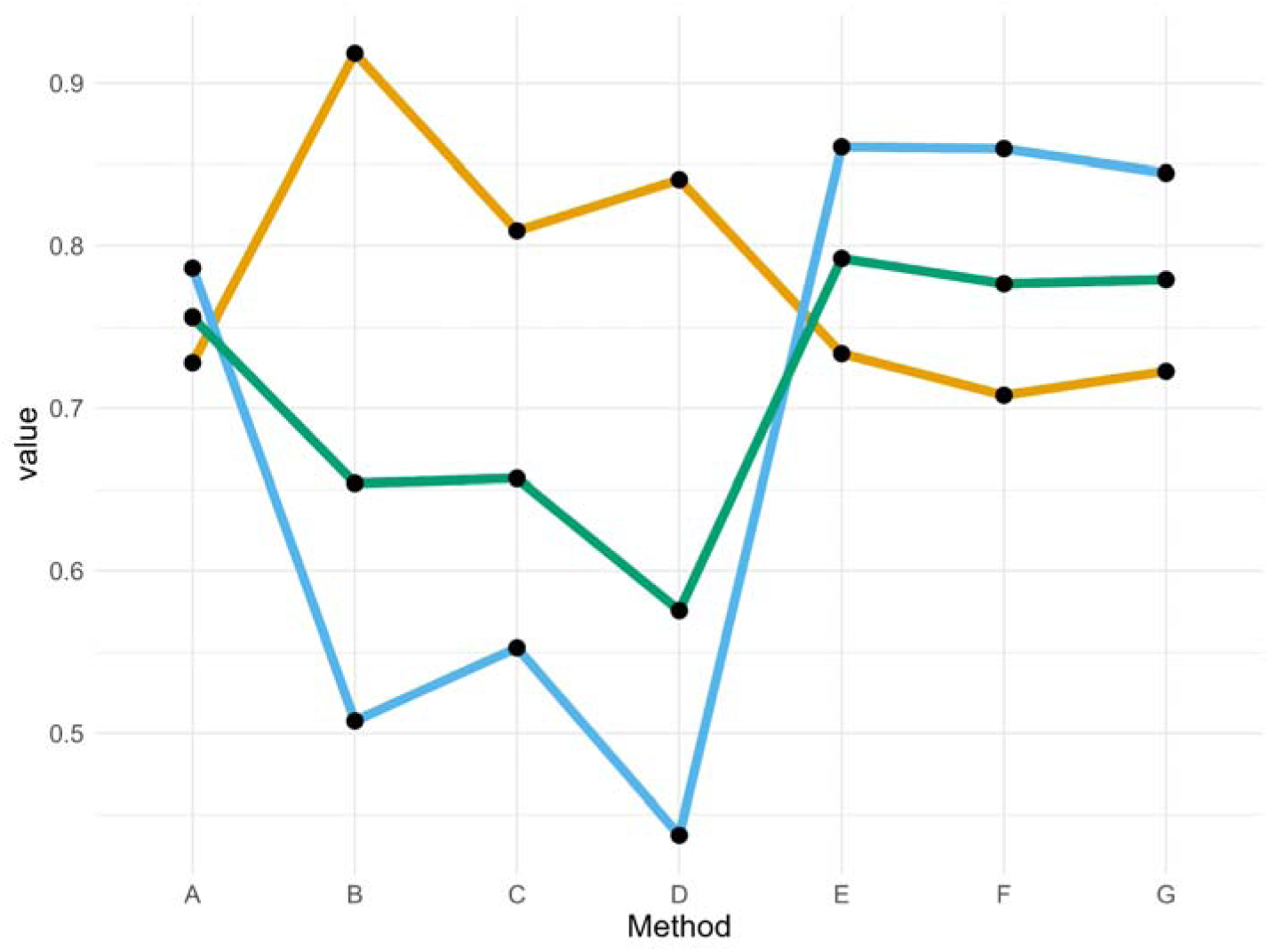
– Precision, recall and F1 scores for different SV calling methods. Values of the precision (orange), recall (blue) and F1 score (green) of SV calling for different methods. Values were based on comparing SVs called using the long-read sequencing data from the reference individual, mapped to BATG-1.0 either as individual reads or assembled contigs, with the “truth” set comprising variants called by mapping the second haplotype of the reference genome to BATG-1.0 and calling SVs with svim-asm. The methods considered were: A) SVs called by both cuteSV and Sniffles2; B) SVs called by svim-asm with a mapping of the *de novo* assembly produced by shasta; C) SVs called by svim-asm with a mapping the *de novo* assembly produced by Flye; D) SVs called by svim-asm with a mapping of *de novo* assembly produced by nextDeNovo; E) SVs called by methods A and/or B; F) SVs called by methods A and/or C; G) SVs called by methods A and/or D.

**Extended Data Figure 2.**
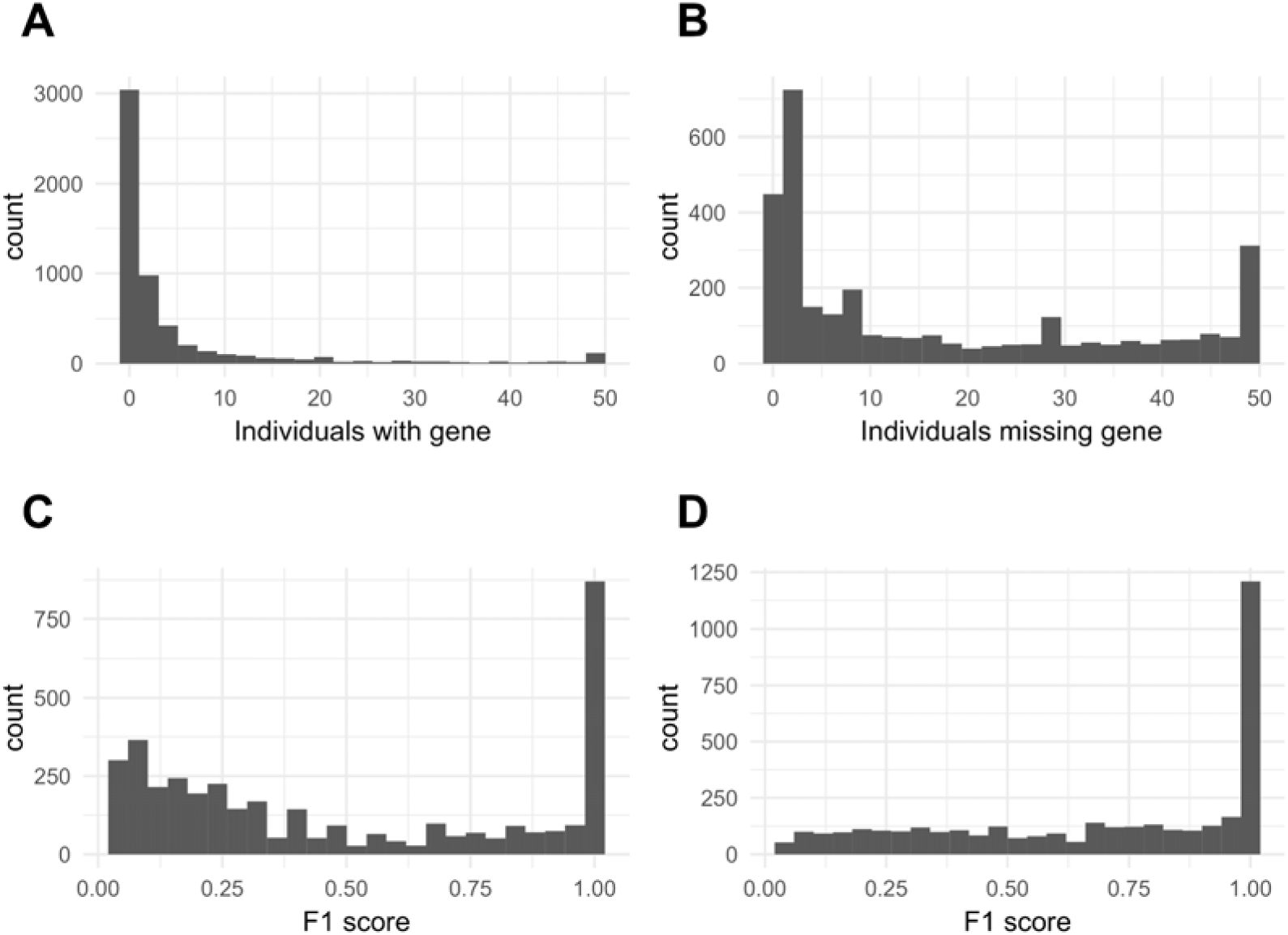
– F1 scores for putatively variable genes association with SVs Histograms showing. A) for genes absent from the linear reference annotation but present in at least one other individual, but not overlapping with an SV, the number of individuals where this gene was present; B) for genes present in the linear reference annotation but absent from at least one other individual, but not overlapping with an SV the number of individuals where the gene was absent, C) for genes absent from the linear reference annotation but present in at least one individual with an SV overlapping the gene location, the best F1 score of the SVs overlapping the gene, D) for genes present in the linear reference individual but absent from at least one individual where an SV overlaps with the gene sequence, the best F1 score of the SVs overlapping the gene.

**Extended Data Figure 3.**
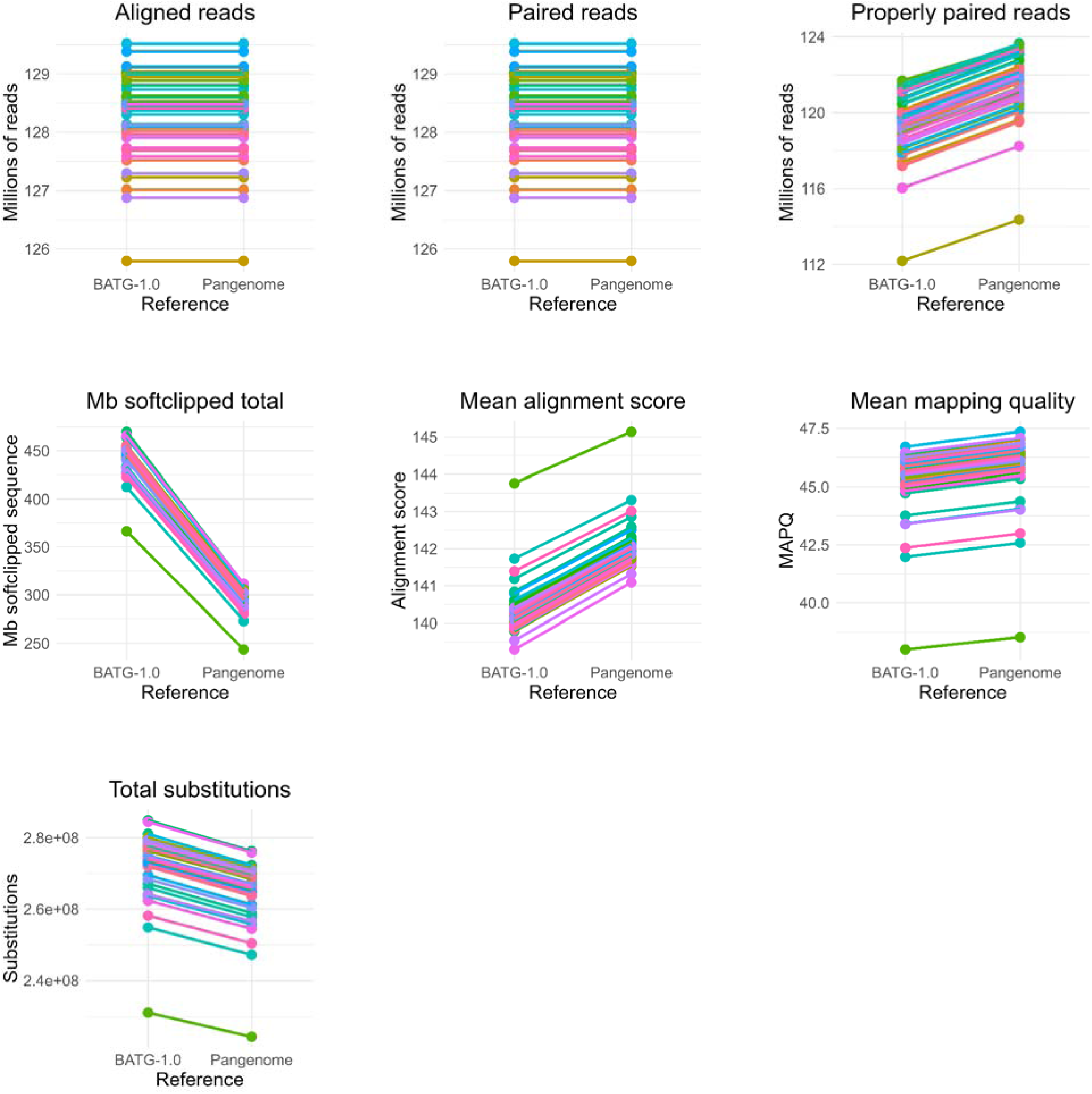
– Read mapping statistics for the linear reference vs. the pangenome. Read mapping statistics for short reads from 42 individuals sequenced in Stocks et al. (2019), mapped using giraffe either to the linear reference genome in pangenome format (BATG-1.0), or to the full pangenome (Pangenome). Each individual is represented by a different colour – lines connect values in the reference-only “pangenome” to the full pangenome in the individual panels. The panels outline i) the total number of aligned reads, ii) the total number of paired reads, iii) the total number of properly paired reads, iv) the number of megabases of softclipped sequences, v) the mean alignment score, vi) the mean mapping quality and vii) the total number of substitutions, as estimated by the vg stats function.

**Extended Data Figure 4.**
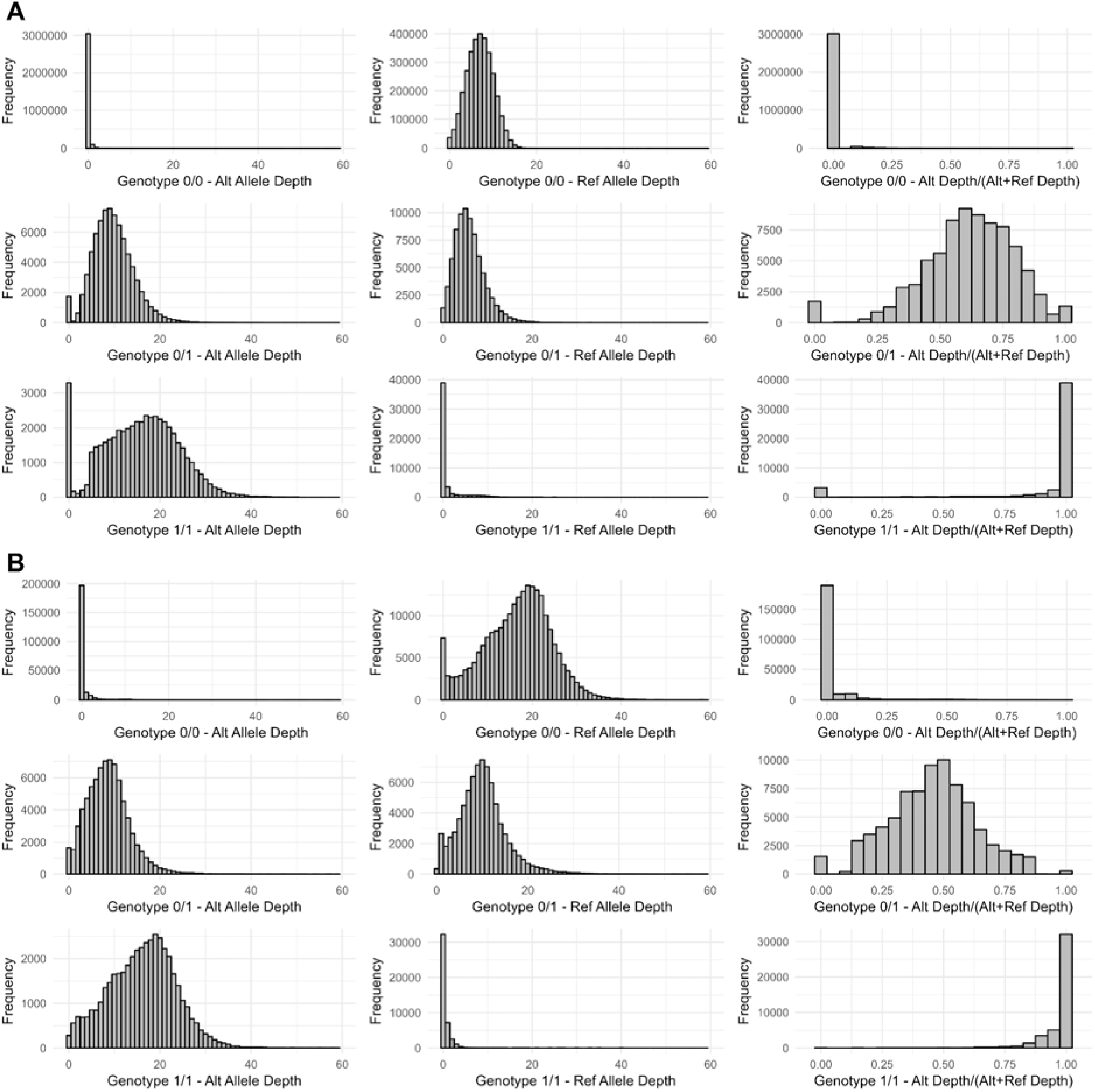
– Allele depths for SNP and SV calls from data for an example individual. Histograms outlining allelic depths for A) SNPs and B) SVs called from an example individual – Individual 164 from Stocks et al. (2019). The panels indicate – for homozygous reference calls (top row), heterozygote calls (middle row) and homozygous alternate calls (bottom row) – the alternate allele depth (first column), reference allele depth (second column) and the alternate allele depth as a proportion of the total allelic depth (last column).

**Extended Data Figure 5.**
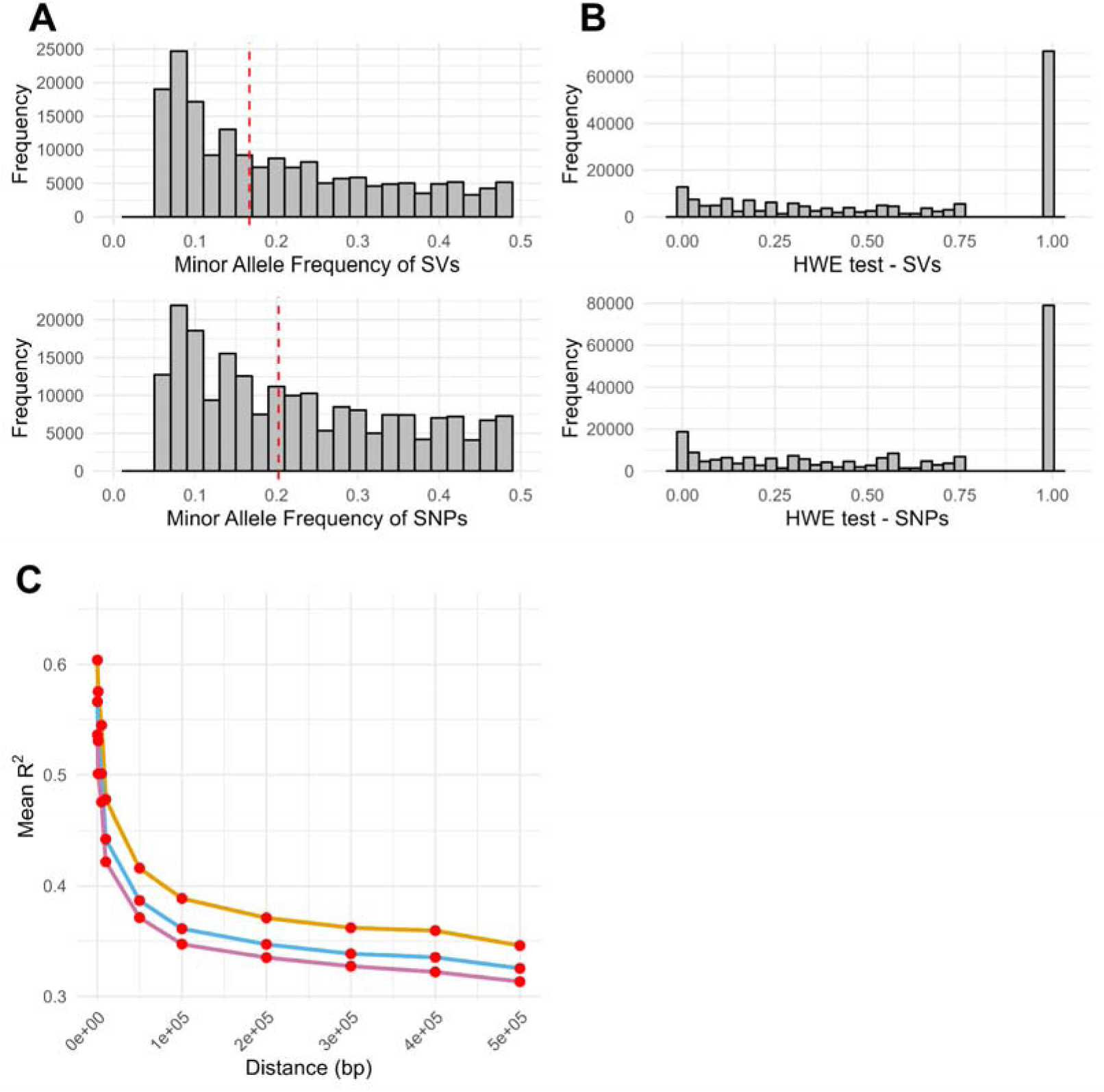
– Statistics of SNP and SV calls combined across individuals. A) Histograms of minor allele frequencies for SNPs (top panel) and SVs (bottom panel) called from a set of individually sequenced ash trees from Stocks et al. (2019). Red vertical dotted line indicates the median value of minor allele frequency for both datasets. B) For the same dataset, estimates of the probability of deviation from Hardy-Weinburg equilibrium (HWE) calculated by bcftools. C) Estimates of linkage disequilibrium decay by distance between SNPs (orange line), SVs (purple line) and between SNPs and SVs (blue line). Pairwise linkage disequilibrium was estimated using plink2, with mean R^2^ values in predefined bins (red points) estimated in R.

**Extended Data Figure 6.**
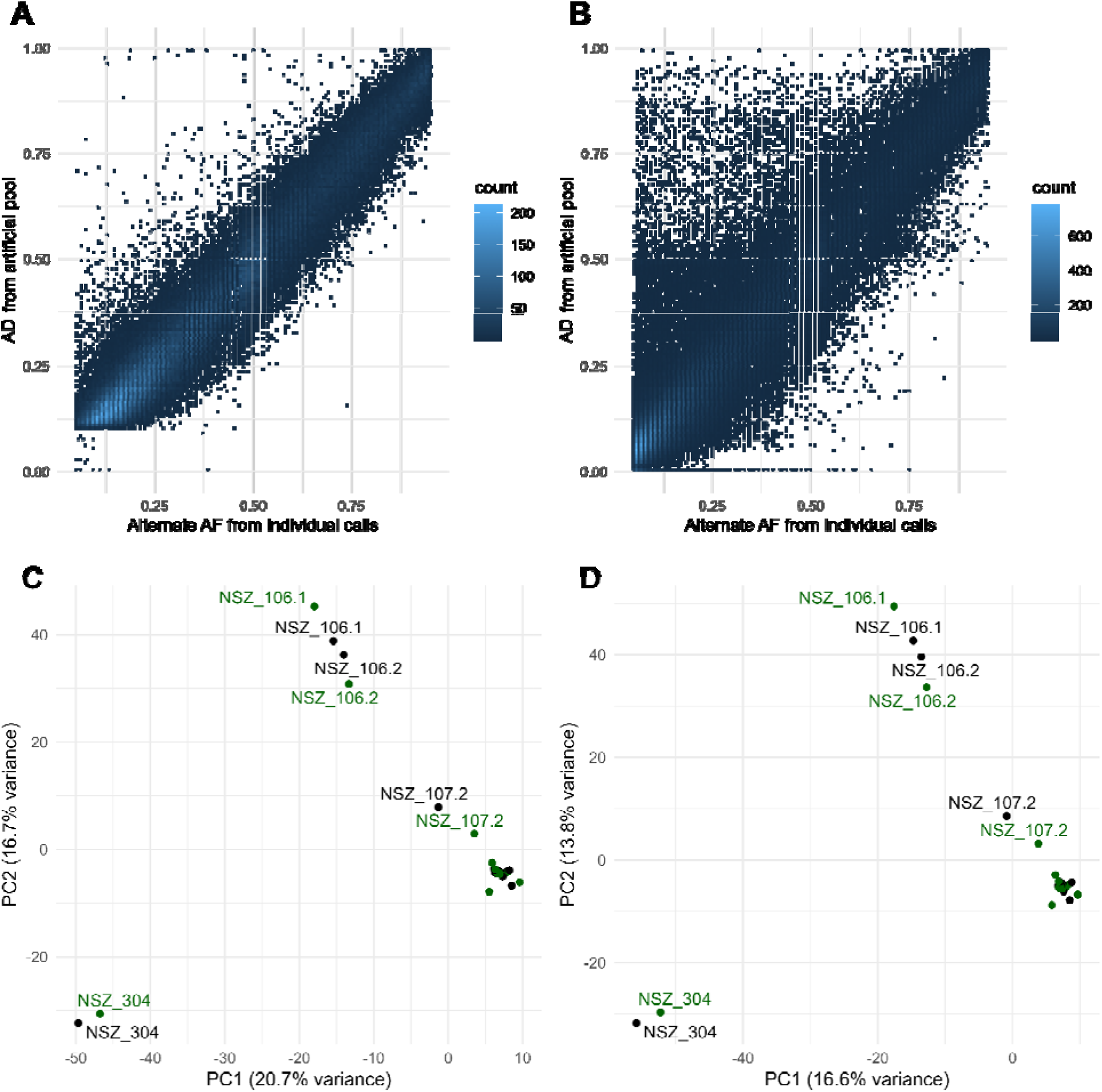
– SV and SNP calling from Poolseq data. A), B) Heatmaps of alternate allele frequencies for SNPs (A) and SVs (B) estimated from individual calls (X axis) from a subset of individually sequenced ash trees from Stocks et al. (2019) vs. an allele frequency estimate from *in silico* pools generated by randomly downsampling reads for these individuals, and measuring the allelic depth (AD; Y axis). Colour scale represents the number of sites in a square. C), D) Plots of the first two principal components (PCs) from principal components analysis from allele frequencies estimated for SNPs (C) and SVs (D) from the poolseq data from Stocks et al. (2019)^42^; axis labels represent the percentage of total variance explained by the PCs. Point labels indicate pool provenances from Stocks et al. (2019) for a subset of pools (i.e. those outside the main dense cluster). “.1” and “.2” represent biological replicate pools. Colours correspond to pool health scores – healthy (green) and unhealthy (black).

**Extended Data Figure 7.**
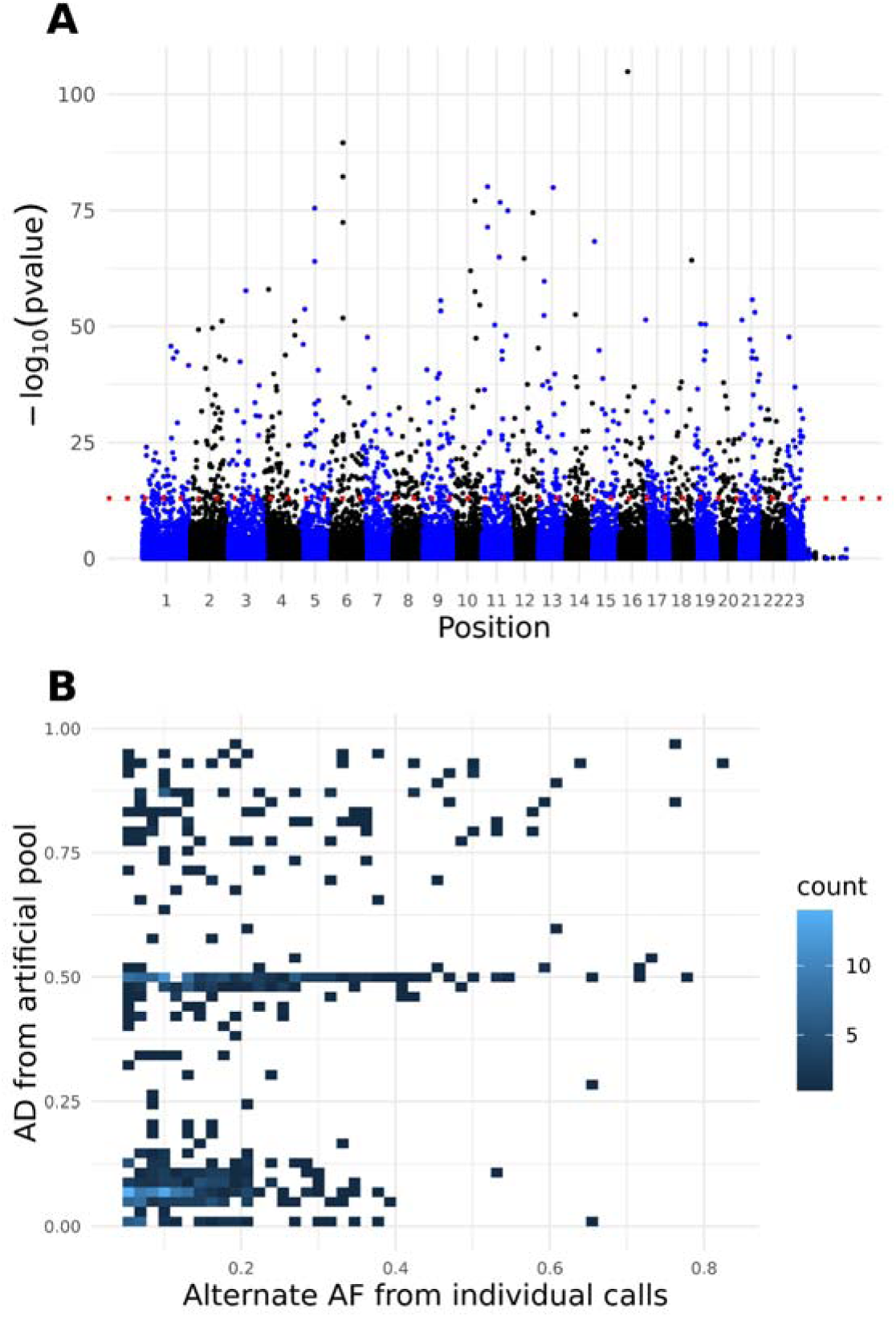
– GWAS using SV allele frequencies estimated from poolseq data. A) Manhattan plot for SVs called from poolseq data from Stocks et al. 2019, identifying association with ash dieback disease damage. Y axis values represent the –log_10_ p-values of a Cochran-Mantel Haenszel (CMH) test (red horizontal line corresponds to p = 1 x 10^−13^). Sites are ordered by position along scaffolds in the BATG-1.0 genome assembly, with alternating black and blue colours representing the different scaffolds. Scaffolds are listed in order of size, with the largest on the left and smallest on the right. B) For SVs identified as significant in the CMH test, heatmap of alternate allele frequencies estimated from individual calls (X axis) from a subset of individually sequenced ash trees from Stocks et al. vs. an allele frequency estimate from *in silico* pools generated by randomly down sampling reads for these individuals and measuring the allelic depth (AD; Y axis). Colour scale represents the number of sites in a square.

**Extended Data Figure 8.**
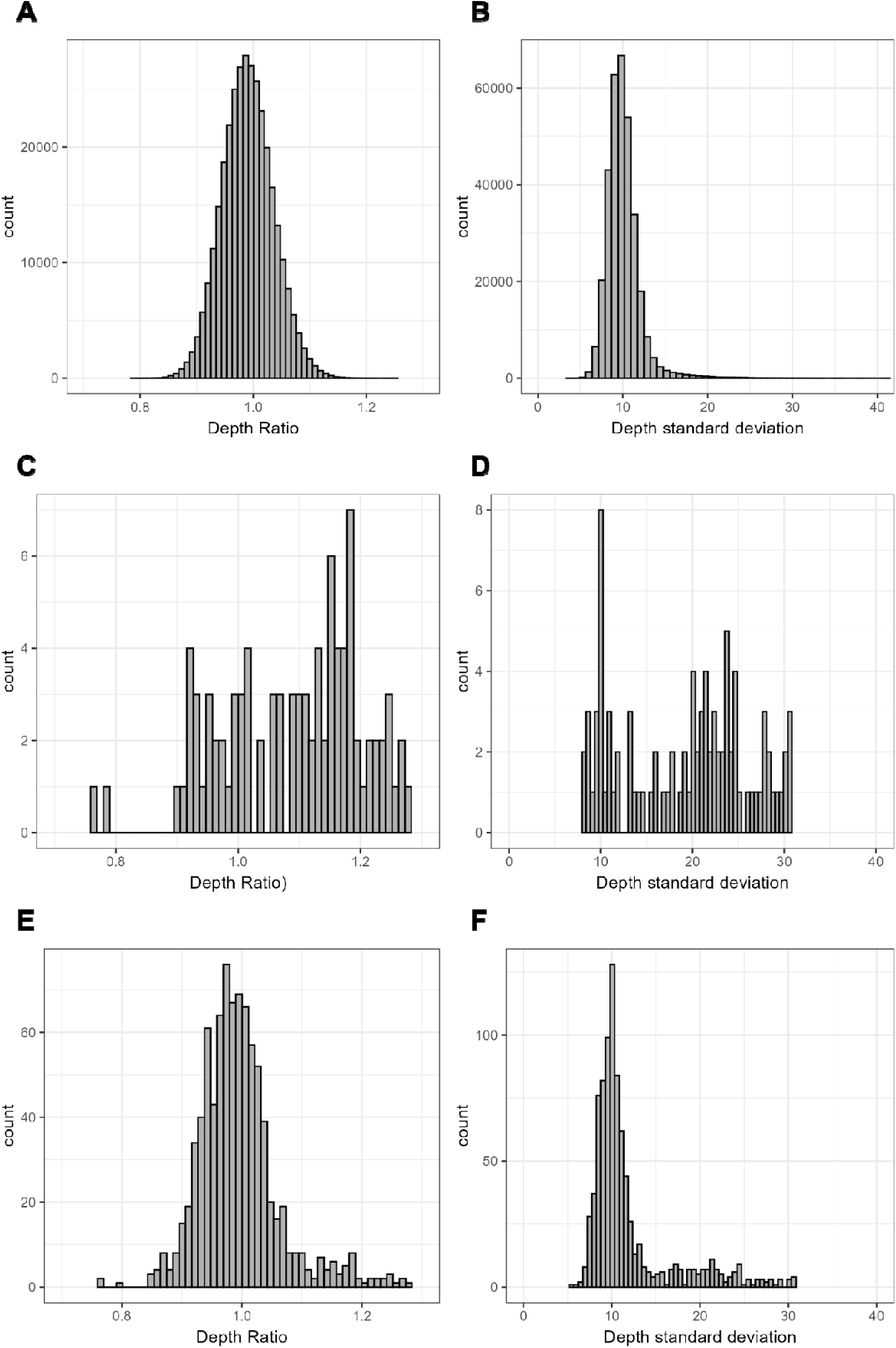
– Depth statistics for SNPs with significant p-values from Cochran-Mantel-Haenszel tests. Histograms showing, for different sets of SNP sites called from the Stocks et al. 2019 poolseq data mapped to the pangenome, i) the ratio of the mean depth of healthy pools vs. unhealthy pools (panels A, C, E), and ii) the standard deviation of read depths across all the pools. Panels indicate sets of SNPs – i) a random 1% of total SNPs (A, B), ii) 92 SNPs with a p-value < 10^−13^ (C, D) and iii) 866 SNPs with a p-value passing Bonferroni correction with LJ = 0.05 (E,F).

**Extended Data Figure 9.**
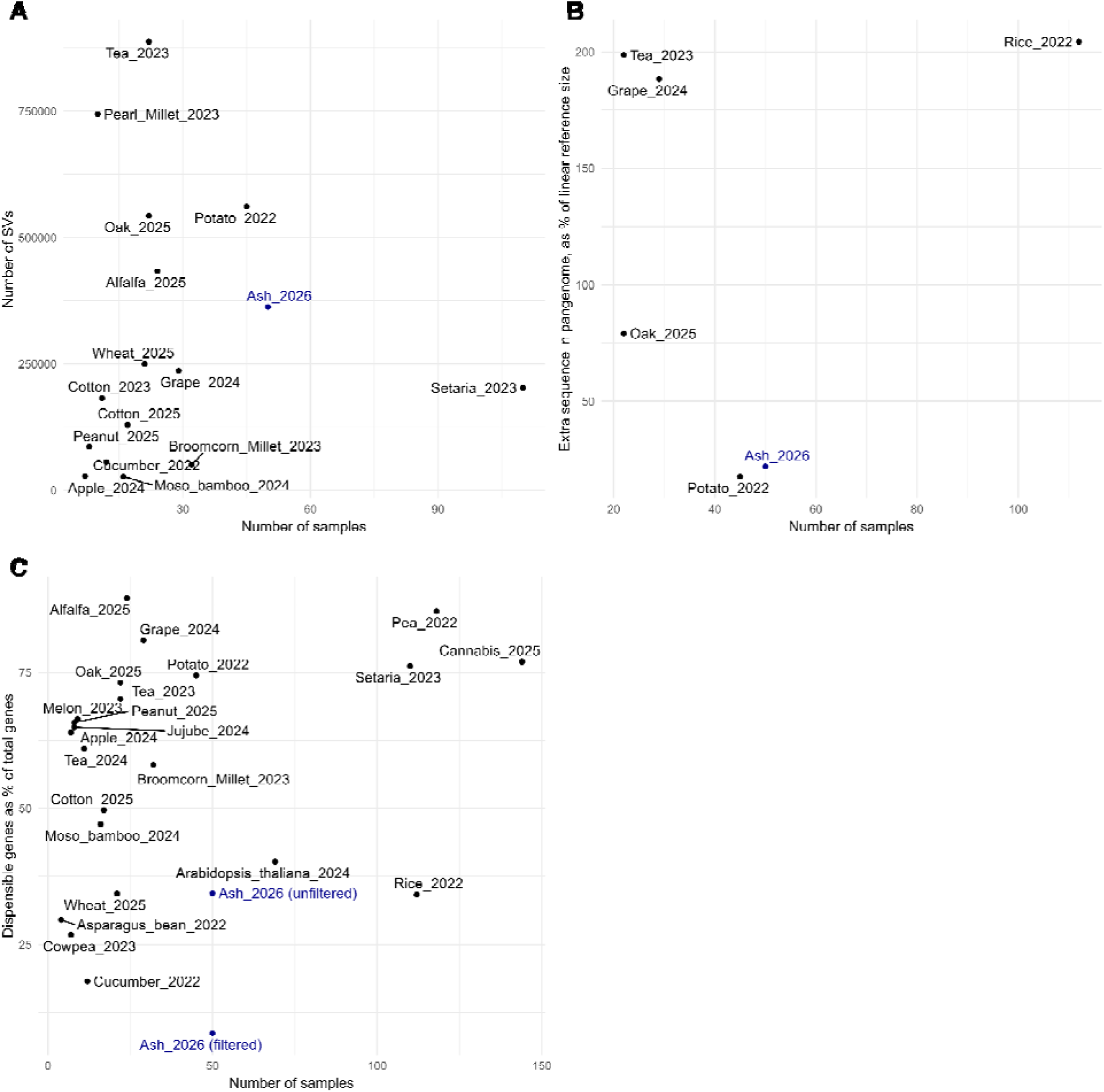
– Comparison of key statistics with other recent pangenome studies. Results from the present study compared with a selection of plant pangenome studies published since 2022; pangenome studies using primarily short-read data, or incorporating multiple species, were excluded. Labels correspond to study organism and publication date, detailed in Supplementary Table 7. Number of samples included in the pangenome is on the X axis. Y axis numbers correspond to total number of SVs discovered (A), additional sequence included in the pangenome, as a percentage of the reference genome size of that species (B), or the number dispensable genes as a proportion of the total number of genes in the pangenome. Results from the present study are in blue; in panel C, Ash_2026 (unfiltered) refers to the proportion of dispensable genes prior to filtering, Ash_2026 (filtered) refers to the proportion after filtering for consistent association with SVs – see Results for further details.

**Extended Data Figure 10.**
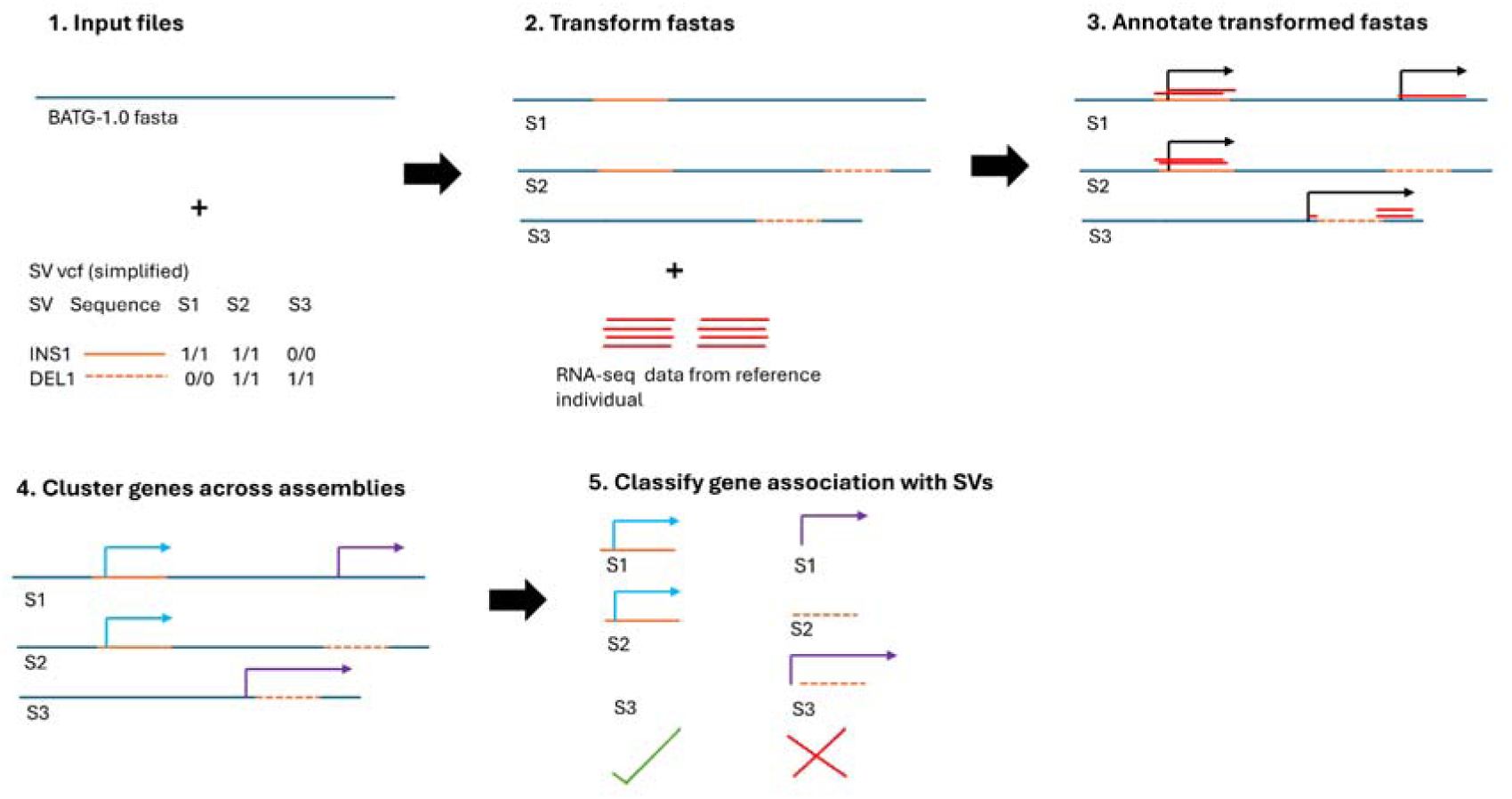
– Explanatory diagram for the pangenome annotation. Explanatory diagram for annotation of the pangenome – see Methods section “Linear reference and pangenome annotation” for full description. For the pangenome annotation, the BATG-1.0 linear reference was transformed using the vcf file of SV calls from each of the 50 ONT samples (1 – S1, S2 and S3 correspond to 3 example samples). For each sample, calls from the vcf used to transform the BATG-1.0 fasta; where an insertion was called, the consensus insertion sequence was inserted at the appropriate position, for deletions the relevant sequence from the BATG-1.0 fasta was deleted. This resulted in a transformed fasta for each sample (2). Using the RNA-seq data from the reference individual, each of the transformed fasta files was annotated (3). Based on sequence homology and position in the genome, genes were clustered across each of the transformed fasta files (4). For each gene cluster, the strength of association of genes with each SV was assessed (5): genes that were consistently associated with an SV were kept as variable genes, whereas those not showing a consistent pattern with an overlapping SV were not.

