## Supplementary Information for "European ash pangenome reveals widespread structural variation and genetic basis of low ash dieback susceptibility"

Contains:

1. Supplementary Notes 1-7
2. Supplementary Figures 1-17
3. Captions for Supplementary Tables 1-6
4. Supplementary File Captions
5. References

### Supplementary Note 1 – Rare SVs distort allele frequency estimates, leading to low CMH test p-values

We investigated whether SNPs with highly significant CMH test p-values may be strongly differentiated in subsets of pools, despite the limited evidence for SNPs associated with ADB low susceptibility that are widely shared across the seed sources. We first considered the 92 SNPs with a p-value of 1 x 10^-13^ (the significance threshold used for the CMH test in Stocks et al. (2019)^1^ - 55/92 of these were in a single 1.3kb region on chromosome 15 (Supplementary Figure 14). The high density of CMH outlier SNPs, combined with raised depths in a subset of pools (Supplementary Figure 15), led us to suspect this region represented a structural variant (SV) not captured in our pangenome, but which may be strongly associated with low ADB susceptibility in a subset of provenances. However, mapping 692 individually sequenced samples to this region using bwa v0.7.17^2^ (150 individually sequenced samples from Stocks et al. 2019^1^, displayed in Supplementary Figure 16B; 542 individuals from Metheringham et al. 2025^3^, not displayed) identified two samples - one from Stocks et al., one from Metheringham et al. - with large increases in depth (29-33X that of the flanking region compared to other samples (Supplementary Figure 16B), which was recapitulated when these two samples were mapped to the full pangenome (Supplementary Figure 16C, D). This suggests the presence in some individuals of large or high copy number SV with high similarity to the 1.3kb region identified in our GWAS. A single individual with this large or high-copy SV could result in a pool with approximately the observed depth increase, giving a highly distorted estimate of SNP allele frequency. The standard deviation of depth, and to a lesser extent the depth of healthy pools relative to unhealthy pools in the top 92 SNPs often exceeded the genome-wide average of these statistics (Extended Data Figure 8) - this was less frequent in the 866 SNPs as a whole that are significant after Bonferroni correction, suggesting this issue is likely to be rare and disproportionately impacting the SNPs with the most extreme p-values from the CMH test (Extended Data Figure 8).

### Supplementary Note 2 - GWAS analysis

Wiberg et al. (2017)^4^ identified a quasibinomial test as an alternative for detecting consistent shifts in allele frequencies. Using this test found no significant SNPs (either with Bonferonni correction with ɑ = 0.05, or p < 10^-13^ ) associated with low susceptibility to ADB, with the lowest p-value being 4.4x10^-7^. There was some evidence of p-value deflation (SV lambda = 0.58; SNP lambda = 0.62; Supplementary Figure 12)**,** suggesting the test could be overly conservative. The top 1000 SNPs showed more consistent shifts than for significant CMH SNPs, but also showed mean allele frequency shift magnitudes that were very similar to a randomly selected background set of SNPs (Supplementary Figure 12B-F). This suggests that the most consistent allele frequency shifts across the pools have a very small average allele frequency shift; in a large enough selection of SNPs, some of these sites would be expected to occur by chance. The qGLM test using the poolFreqDiffTest_QGLM.py script from the PoolFreqDiff repository^4^ by default does not include an effect for pool stratification – including a fixed effect for provenance resulted in substantial p-value inflation (Supplementary Figure 13). Including fixed effects for the first two principal components of the PCA from poolseq SNP allele frequencies (Extended Data Figure 6C) resulted in more modest p-value inflation (Supplementary Figure 13B), identifying a single significant SNP with Bonferonni correction with ɑ = 0.05, but none with p < 10^-13^. Using a binomial GLM with a random effect of provenance, or a binomial GLM and a random effect of provenance and observation (the latter allowing for more overdispersion) resulted in extreme p-value inflation (Supplementary Figure 13C, 13D).

### Supplementary Note 3 – Per pool differentiation estimates

To more explicitly test allele frequency shifts within subsets of provenances, we calculated i) Fisher’s exact test using the fisher-test.pl function in Popoolation2, and ii) F_st_, in 500bp/100bp sliding windows using the fst-sliding.pl function in Popoolation2, for each pair of healthy vs. unhealthy pools. We then looked at windows shared across multiple pairs of pools, either by multiple pairs sharing i) a 500bp/100bp sliding window containing a SNP passing a significance threshold for the Fisher’s exact test SNPs (p < 0.05 after Bonferroni correction for the number of sliding windows), ii) a window in the top 1% of F_st_ value for each pair of pools. It should be noted that absolute F_st_ may lack power to detect small-effect SNPs diverging over very short timeframes, but regions in the top 1% of values may nevertheless be candidates for differentiated regions.

168,378 windows contained a significant Fisher’s exact test SNP in at least one pair of pools - however, only 3,937 of these were shared across more than one pair. For both pairs of biological replicate pools from Stocks et al. 2019, the overlap was very low - only 0.2% of all windows identified across the two pairs of biological replicates were identified as outliers in both pairs. Whilst 101,334 windows were found to have Fst values in the top 1% of values in more than one pool, only 5.1% and 4.9% of the outliers for replicates for NSZ-106 and NSZ-107 that were common to more than one pool, were shared with their respective biological replicate. The clustering of biological replicates in a PCA of SNP allele frequency (Extended Data Figure 6) indicates that in aggregate, these clearly data contain biological signal - but high confidence allele frequency shifts at individual sites are difficult to distinguish from stochastic fluctuations in individual read counts inherent to poolseq data. We therefore do not believe it is possible to detect more geographically restricted genetic variation associated with low ADB susceptibility using the present dataset.

### Supplementary Note 4 – Effect of using the BATG-0.5 reference genome

To compare how the results of the SNP-based CMH test would be affected by using the pangenome, the GWAS was repeated using the BATG-0.5 reference genome, which was the first published reference assembly for *F. excelsior* and has been used for previous GWAS. Reads from the poolseq files were mapped to the assembly using bwa 0.7.17. Samtools v1.9^5^ was used to remove PCR duplicates and reads with MAPQ < 20. From these bam files, the same approach as for the pangenome based analysis was used to create .sync files and perform CMH tests. To compare these results with those generated from the pangenome, a chain file between BATG-0.5 and BATG-1.0 was created using nf-lo v1.8.0^6^ using settings –distance medium –aligner minimap2. Using the –annotation option, the positions of the significant SNPs from the BATG-0.5 GWAS were estimated in the new assembly. This resulted in 116 SNPs with p < 10^-13^. A liftover of these 116 SNPs from BATG-0.5 identified 91 of the 92 sites detected via the pangenome-based GWAS. Reads mapping to significant sites in the BATG-0.5 reference genome were mapped to the pangenome; these mapped to 82 of the 92 significant sites identified from the GWAS using the pangenome. Similarly, reads mapping to significant sites in the pangenome were mapped to BATG-0.5, covering 77 of the 116 significant sites. Only 17 SNPs identified as significant from mapping to the pangenome corresponded to the 192 SNPs in BATG-0.5 found as significant in Stocks et al.^1^, as identified by the liftover, despite an equivalent position in BATG-1.0 being identified for all 192 sites from BATG-0.5. The above analysis indicates this is unlikely to be due to the different reference genomes used but is instead likely the result of different choices of parameters in the two studies. For example, in this study, sites were removed if a pool had a depth > 200, whereas in Stocks et al.^1^ this value was 3,000. By contrast, if sites overlapped with a repeat sequence in Stocks et al.^1^ they were removed, whereas in this study they were retained.

### Supplementary Note 5 - RNA extraction Protocol

High integrity RNA extraction from fresh/flash frozen *F. excelsior* leaves and cambium, modified from the Qiagen RNeasy Plant Mini Kit Protocol. Outputs; 10-20ug RNA, >100ng/ul, 260/280 and 260/230 >1.8, RIN > 7, ND/QT ratio < 2. <2 hours for 2 samples.

**Materials**

- 25-30mg fresh / flash frozen leaf tissue
- QIAGEN RNeasy Plant Mini Kit
- B-mercaptoethanol
- Qiagen RNase-free DNase Set
- Liquid Nitrogen
- Mortar and pestle

**Method**

1. Before starting, prepare buffers RLT and RPE as outlined in the Rneasy Plant Mini Kit quick start protocol. Note that each sample will require 2x the volume of buffers/reagents as outlined in the kit handbook. Prepare Dnase aliquots as outlined in the kit handbook.
2. Disrupt 25-30mg tissue in liquid nitrogen and add to a 2ml centrifuge tube. Do not allow the tissue to thaw.
3. Add 450ul Buffer RLT and vortex.
4. Transfer the lysate to a QIAshredder spin column (lilac) placed in a 2 ml collection tube. Centrifuge for 2 min at full speed. Transfer the supernatant of the flow-through to a new microcentrifuge tube (not supplied) without disturbing the cell-debris pellet.
5. Add 0.5 volume of ethanol (96–100%) to the cleared lysate and mix immediately by pipetting.
6. Transfer the sample (usually 650 μl), with any precipitate, to an RNeasy Mini spin column (pink) in a 2 ml collection tube (supplied). Close the lid and centrifuge for 15 s at ≥8000 x g (≥10,000 rpm). Discard the flow-through.
7. Add 350ul Buffer RW1 to the RNeasy spin column.
8. Close the lid and centrifuge for 15s at ≥8000 x g.
9. Add 10ul DNase I stock solution to 70ul Buffer RD. Mix by inversion.
10. Add the mixture directly to the RNeasy spin column, incubate at room temperature for 15mins.
11. Add 350ul Buffer RW1 to the RNeasy spin column. Close the lid and centrifuge for 15s at ≥8000 x g. Discard the flow-through.
12. Add 500 μl Buffer RPE to the RNeasy spin column. Close the lid, and centrifuge for 15 s at ≥8000 x g. Discard the flow-through.
13. Add 500 μl Buffer RPE to the RNeasy spin column. Close the lid, and centrifuge for 2 min at ≥8000 x g.
14. Place the spin column in a new 2ml tube. Centrifuge at 1min to dry the membrane.
15. Add 450ul Buffer RLT to a new centrifuge tube and place the spin column in this tube.
16. Add 50ul Rnase free water directly to the spin column membrane. Close the lid, and centrifuge for 1min at >8000g to elute the RNA.
17. Discard the spin column.
18. Using the flow through as the lysate, repeat the procedure without the Dnase step, using new spin/shredder columns (i.e. Steps 4-7, 11-14). Then transfer the spin column to a new empty 2ml tube and repeat Step 16 to elute the RNA. Store at -80*C.

### Supplementary Note 6 - DNA extraction Protocol

High molecular weight gDNA extraction from fresh/flash frozen *F. excelsior* leaves. Modified from protocol outlined in Franco Ortega et al.^6^. Outputs; 8-35ug DNA, >100ng/ul, 260/280 and 260/230 >1.8, peak fragment size > 60kb, ND/QT ratio < 2. Including sample disruption and QC, this usually took 2 days for 12 samples.

**Materials**

- 300-400mg fresh / flash frozen leaf tissue (~3 “large” leaflets to 10 “small” leaflets)
- QIAGEN Genomic-tip 100/G and associated Qiagen buffers
- Carlson buffer, pre-warmed to 65° C:
  - 100 mM Tris-HCl, pH 9.5
  - 2% CTAB
  - 1.4 M NaCl
  - 1% PEG 8000
  - 20 mM EDTA
  - To ensure all CTAB is dissolved, stir the Carlson buffer overnight
- B-mercaptoethanol
- Chroloform
- Isopropanol
- Proteinase K (20 mg/ml)
- RNase A (100 mg/ml)
- Liquid Nitrogen
- Mortar and pestle
- Vortex mixer
- 50 ml Falcon tubes
- Centrifuge capable of taking 50 ml tubes
- Water baths at 65° C, 50° C and 55° C

**Method**

Note: use wide-bore pipette tips throughout. Gently invert to mix; do not vortex after step 3.

1. Transfer 8ml of Carlson buffer to a 50 ml Falcon tube. In a fume hood, add 20 µl ß-mercaptoethanol to the Carlson buffer, mix by vortexing and pre-warm to 65° C in a water bath. Immediately before use, add 100 µl proteinase K.
2. Pre-cool the mortar and pestle with liquid nitrogen until both are at -80° C. This keeps the sample as a fine powder during grinding and will prevent re-activation of intracellular DNases. Pour ~30 ml of liquid nitrogen into the mortar and add 300-400 mg leaves. When the liquid nitrogen has evaporated, grind the tissue for approximately 30 seconds, to a flour-like consistency. Keep the sample at the bottom of the mortar as much as possible. Add another ~30 ml of liquid nitrogen and repeat grinding for approximately 30 seconds. Perform 2 - 4 cycles of grinding in total. Note: making sure the sample does not thaw, you can place foil over the mortar containing the sample and store at -80 - then you can do the downstream steps simultaneously (I did up to 12).
3. Transfer the frozen powdered tissue to the tube with the pre-warmed Carlson buffer and vortex the tube for 5 sec. Immediately transfer the tube to a 65° C water bath and incubate for 1 hour, mixing the sample by inversion at regular intervals (every 10 mins or so).
4. Add 40 µl of RNase A (100mg/ml), mix by inversion and incubate for a further 15 minutes.
5. Let the sample cool to room temperature, then add 1 volume of chloroform. Mix the sample by inversion.
6. Centrifuge the sample at 5500 g for 10 mins at 4° C.
7. Carefully transfer the top (aqueous) phase to a new 50 ml Falcon tube, without disturbing the interphase.
8. Repeat the chloroform wash steps 5-7 above.
9. Add 0.7 volumes of isopropanol to the top phase and mix thoroughly by inverting the tube 10 times. Place the tube at 4°C for 15 mins.
10. Centrifuge the sample at 5500 g for 30 mins at 4° C. Carefully discard the supernatant, without disturbing the pellet.
11. (Optional) If not proceeding to g-tip clean up, the pellet can be resuspended in an appropriate volume low TE (250ul Qiagen Buffer AE) and stored at 4*C overnight. Alternatively, proceed to g-tip clean up as below.
12. Carefully dissolve DNA pellet at 5 ml of G2 buffer. Place the sample in a 50° C water bath for 15 mins, mixing occasionally until the pellet dissolves.
13. Equilibrate a QIAGEN Genomic-tip 100/G column with 4 ml of Buffer QBT.
14. Apply your fully dissolved DNA in G2 buffer to the equilibrated QIAGEN Genomic-tip 100/G column. Allow the DNA to enter the resin by gravity flow.
15. Wash the QIAGEN Genomic-tip 100/G with 10 ml of Buffer QC. Wait until all the buffer flows through the resin and repeat the wash with another 10 ml of Buffer QC. (Note; this is more than in the standard protocol, so a Genomic Buffer Kit will not have enough QC for 25 samples).
16. Place the QIAGEN Genomic-tip 100/G over a clean 50 ml Falcon tube and elute the genomic DNA with 5 ml of Buffer QF, pre-warmed to 55°C. Allow the eluate to cool down room temperature.
17. Precipitate the DNA by adding 0.7 volumes (3.5 ml) of room temperature isopropanol to the eluted DNA and mix by inverting the tube several times. Incubate at room temperature for 15 min.
18. Centrifuge at 5500 g for 30 mins at 4° C and carefully remove the supernatant. The pellet will likely be highly fragmented, the supernatant will need to very carefully pipetted off.
19. Wash the centrifuged DNA pellet with 4 ml of cold 70% ethanol. Invert the tube several times to disturb the pellet, and centrifuge at 5500 g for 10 min at 4°C.
20. Carefully remove the supernatant without disturbing the pellet. The pellet will likely be highly fragmented, the supernatant will need to very carefully pipetted off.
21. Air dry the pellet for 10 min and resuspend the DNA in 100 μl of 1x TE buffer pH 8.
22. For high concentrations/quantities of DNA (>100ng/ul), the DNA will take a long time to resuspend. Leaving for 3 days at room temperature and 4 days at 4°C (highly variable readings on a spectrophotometer from a single sample suggests it has not fully resuspended).

### Supplementary Note 7 – Pangenome annotation clustering

To avoid missing genes due to stochasticity in the BRAKER3 pipeline, we ran 10 runs of the pipeline using short read RNA-seq data, and 10 runs using the long read RNA-seq data. Across each run, genes were clustered using OrthoFinder^7^, with each non-overlapping orthogroup cluster treated as a separate gene. 38,766 non-overlapping clusters were initially detected, with 35,524 containing exactly 20 sequences (i.e. annotated in every run), or 10 sequences (i.e. annotated in every long or short read run). This corresponded well with the number of genes in assemblies such as BATG-0.5^8^. However, 91 pairs of genes had the same start/end positions in the genome, with manual inspection identifying short deletions of particular exons likely being responsible, suggesting in some cases OrthoFinder was likely over-splitting genes in some cases. Any approach to clustering genes across different annotations has an inherent chance of clustering sequences together that should be separate genes, and also clustering sequences as two separate genes when they should be the same gene. To investigate the potential impact of this clustering, we identified cases where a coding sequence overlapped more than 50% with coding sequence from another gene on the same strand, and kept the longest protein coding gene. This resulted in a reduction of the number of genes from 38,766 (untrimmed BATG-1.0 gene set) to 37,983 (trimmed BATG-1.0 gene set). Whilst this potentially discards some real genes, we were concerned that downstream clustering of genes in the pangenome with oversplit genes may have led to oversplitting of pangenome genes as well, with potential downstream consequences on our estimation of the reliability of our dispensable gene assignments.

Running OrthoFinder on proteins sequences from the pangenome samples, along with both these gene sets, resulted in very similar numbers of genes - 48,324 non-overlapping Orthogroups when using the BATG-1.0 untrimmed gene set, and 48,356 when using the trimmed gene set. Further merging these gene sets, if any two genes on the same strand overlapped in their coding sequence by more than 50%, resulted in 45,508 genes when using the untrimmed BATG-1.0 sequences, and 45,114 genes when using the trimmed BATG-1.0 sequences. In some cases where the pangenome genes were merged, genes that did not overlap at all were merged, as they were connected by a series of overlapping intermediate genes – with 100 annotation files, the risk of erroneous gene fusions potentially increases.

We ran the analysis to detect the relationship between SVs and gene presence for each of the four combinations of analysis (untrimmed or trimmed BATG-1.0 gene set, merged or unmerged pangenome gene set) – the overall proportions of dispensable genes, and the generally low concordance of variable genes with SVs, was very similar for each combination of approaches (Supplementary Figure 17), although proportion of dispensable genes was slightly higher using the unmerged pangenome set (8.7% and 8.6% for the trimmed and untrimmed BATG-1.0 sequences, respectively) compared to the merged pangenome gene set (7.2% and 7.2% for the trimmed and untrimmed BATG-1.0 sequences, respectively). We judged that the trimmed BATG-1.0 and untrimmed pangenome datasets appeared to strike the best balance between avoiding undersplitting and oversplitting. Regardless of the exact approach of merging genes across annotation runs, the overall conclusion that many putatively dispensable genes are not reliably associated with sequence differences caused by SVs, remains the same.

### Supplementary Figures

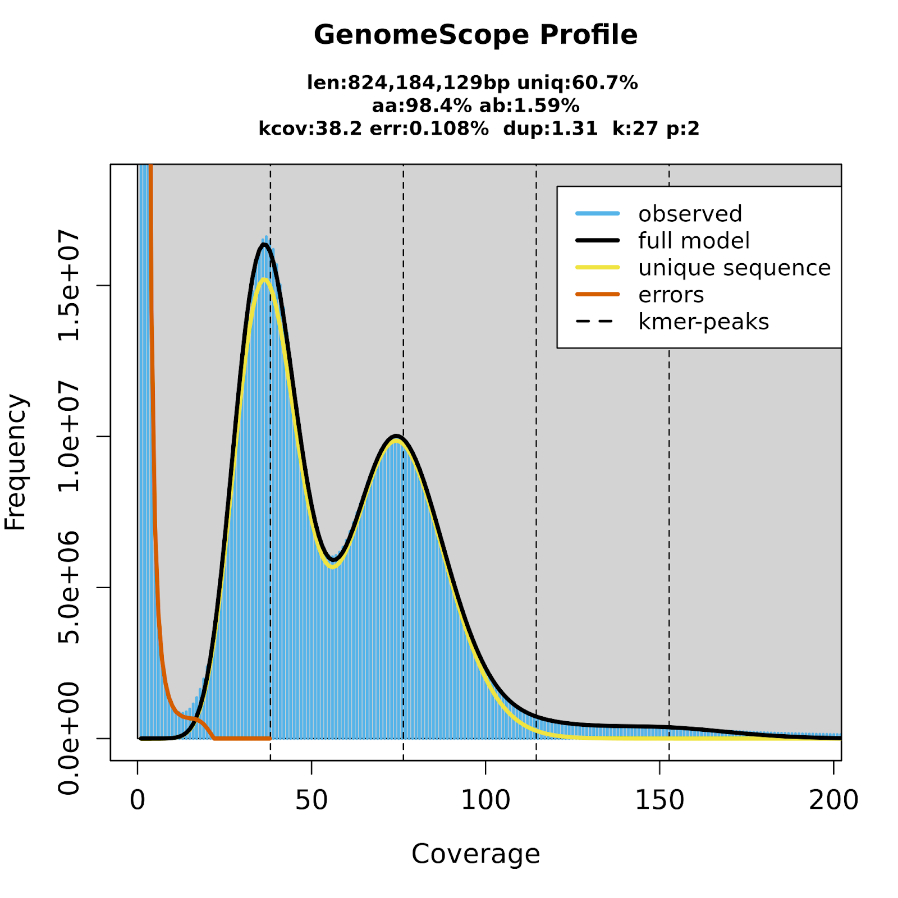

**Supplementary Figure 1: kmer coverage profile for BATG-1.0 PacBio HiFi reads**

Frequencies and coverage of 27bp kmers from PacBio HiFi data for the BATG-1.0 individual using Genomescope.

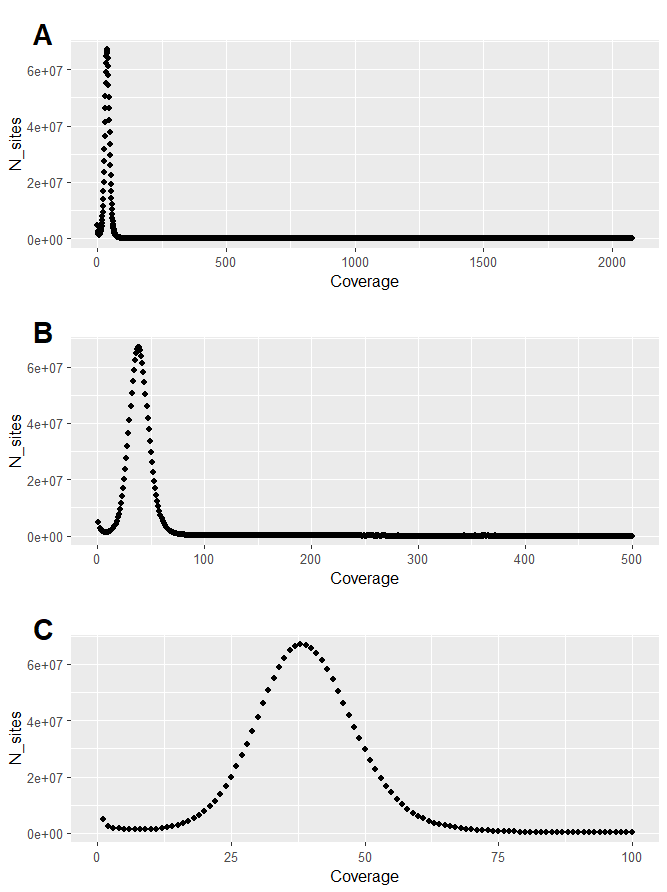

**Supplementary Figure 2 – coverage profiles of phased linear reference genome**

Coverage profiles of PacBio long-reads used to construct the phased assemblies, mapped to a concatenated fasta file of both haplotypes, with per-site coverage on the X axis, and the number of sites displaying this coverage on the Y axis. For legibility, the same data are displayed with X axis cutoffs of A) 2086 (maximum per-site coverage value was 2076), B) 500 and C) 100.

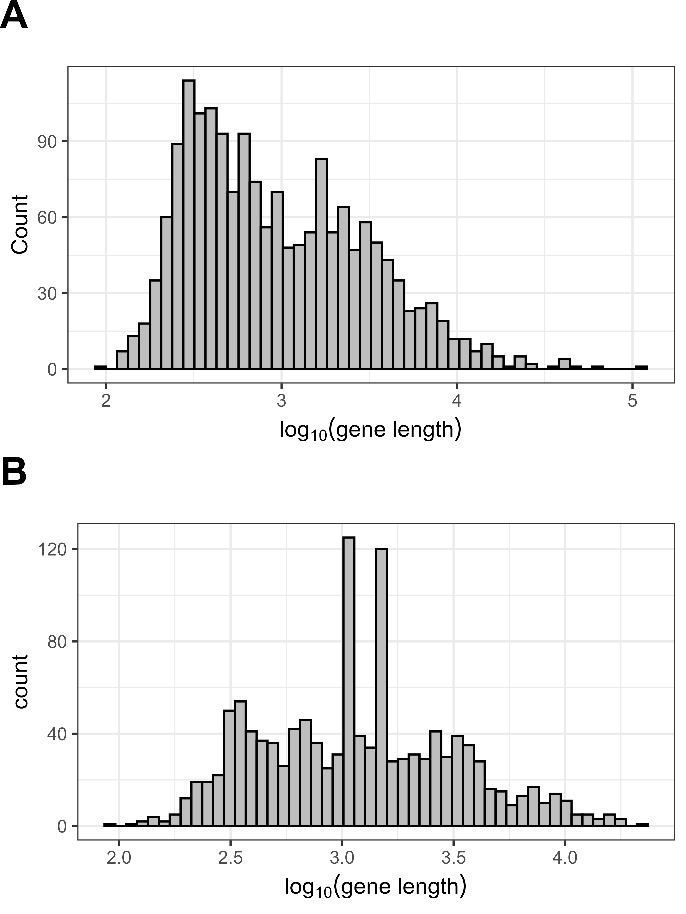

**Supplementary Figure 3 – length distributions for genes identified exclusively with one RNA-seq data type**

Histograms of log_10_ transformed gene length for genes in BATG-1.0 identified exclusively by short-read RNA seq data or long-read RNA seq data. A) Lengths for genes identified in all short-read BRAKER3 runs, but no long-read BRAKER3 runs, and B) genes identified in all long-read BRAKER3 runs, but no short-read BRAKER3 runs.

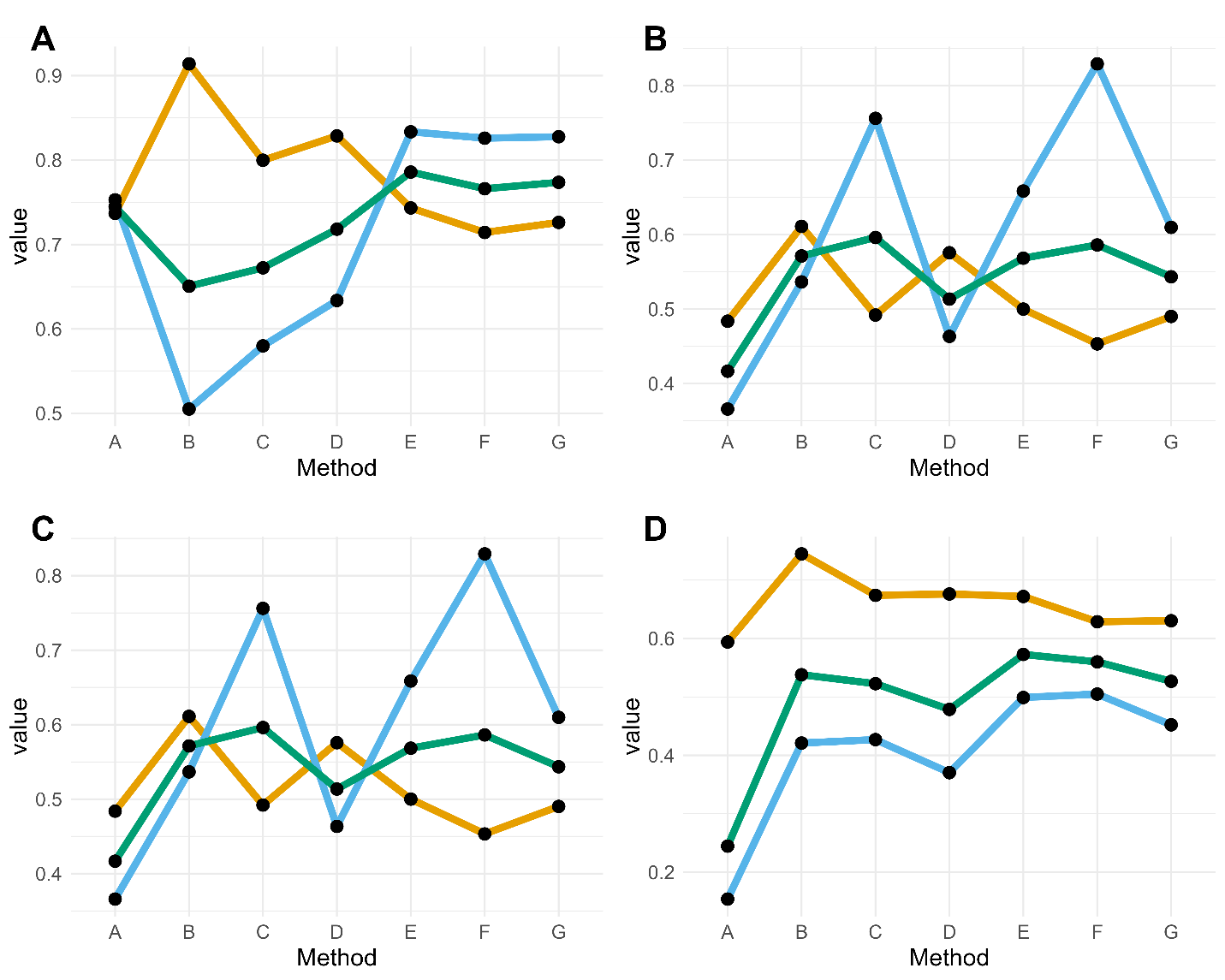

**Supplementary Figure 4 – Class specific precision/recall/F1 of structural variants in BATG-1.0**

Values of the precision (orange), recall (blue) and F1 score (green) of structural variant (SV) calling for different methods. Values were based on comparing SVs called using the long-read sequencing data from the reference individual, mapped to BATG-1.0 either as individual reads or assembled contigs, with the “truth” set comprising variants called by mapping the second haplotype of the reference genome to BATG-1.0 and calling SVs with svim-asm. The methods considered were: A) SVs called by both cuteSV and Sniffles2; B) SVs called by svim-asm with a mapping of the *de novo* assembly produced by shasta; C) SVs called by svim-asm with a mapping the *de novo* assembly produced by Flye;  D) SVs called by svim-asm with a mapping of *de novo* assembly produced by nextDeNovo;  E) SVs called by methods 1 and/or 2; F) SVs called by methods 1 and/or 3; G) SVs called by methods 1 and/or 4. Each panel denotes these values for A) Insertions, B) Deletions, C) Inversions and D) Tandem Duplications.

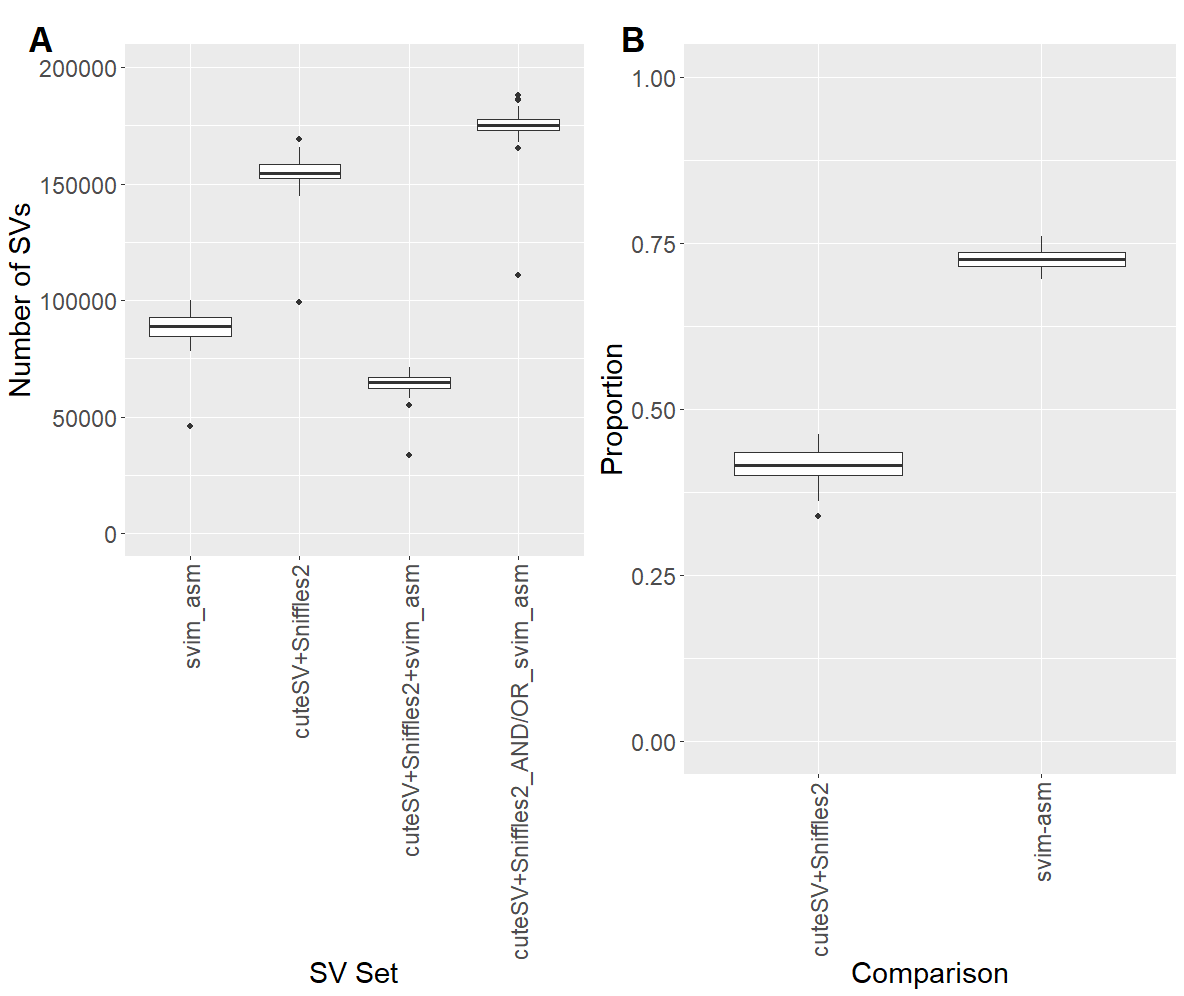

**Supplementary Figure 5 – structural variant caller concordance across different individuals**

A: Boxplots showing, for each of the methods outlined on the X axis, the total number of structural variants (SVs) called per sample. Methods are either i) SV called by svim-asm, ii) SVs called by both cuteSV and Sniffles2, iii) SVs called by cuteSV, Sniffles2 and svim-asm, iv) SVs called by either cuteSV and Sniffles2, and/or SVs called by svim-asm. B: Boxplots showing, for each sample, the proportion of total SVs called per sample called by i) cuteSV and Sniffles2, and ii) svim-asm, for the set of SVs called by either/both of these methods

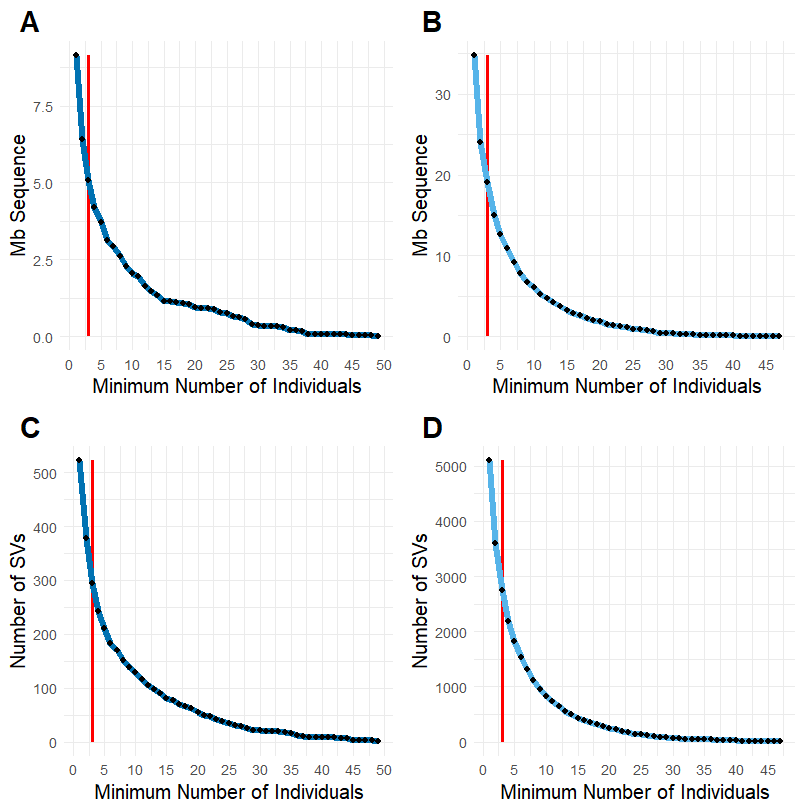

**Supplementary Figure 6 – Inversion and tandem duplicate frequencies**

The same data as Figures 2A and 2B are plotted for inversions and tandem duplicates separately to aid visualisation. The total quantity of sequence (A, B) and total number of SVs (C, D) of different SV classes, at different minimum numbers of individuals called from long-read data. Data for inversions are plotted in panels A and C, data for tandem duplications in panels B and D A vertical red line at n = 3 indicates the cutoff for SVs being included in the pangenome.

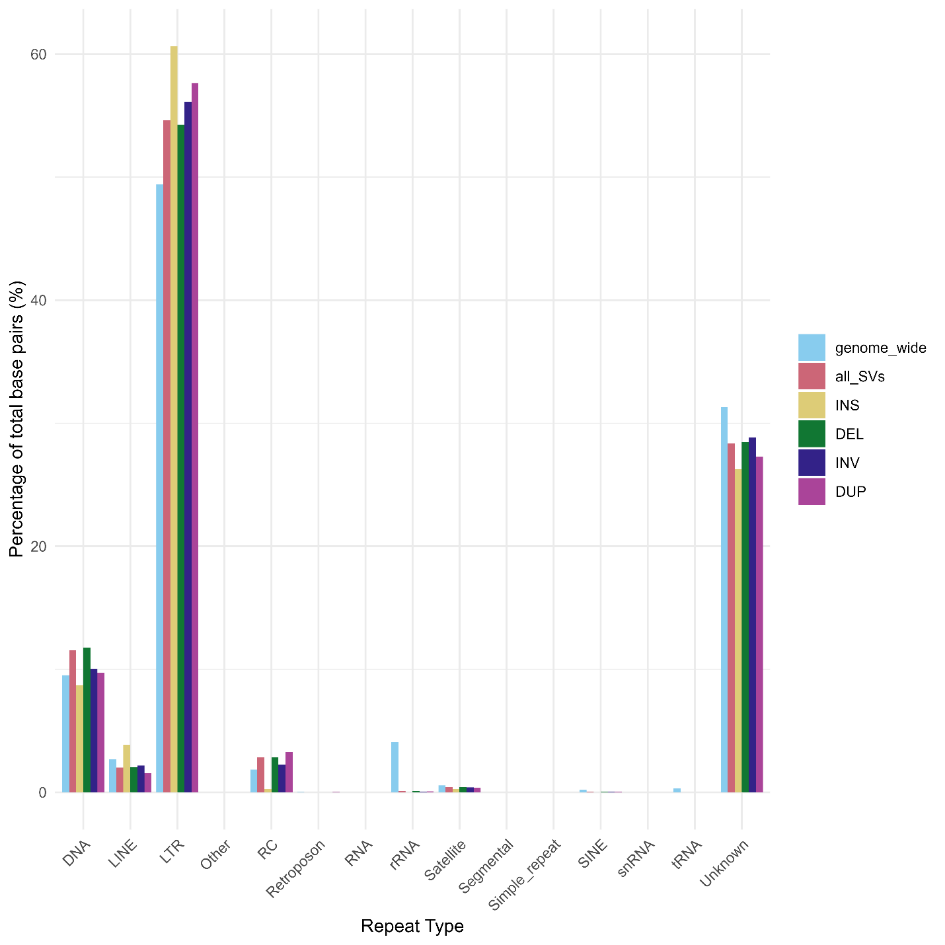

**Supplementary Figure 7 – Classes of repeats overlapping structural variants.**

For i) the genome as a whole, ii) all classes of structural variants, iii) insertions (INS), iv) deletions (DEL), v) inversions (INV) and tandem duplications (DUP), the percentage of base pairs overlapping with various classes of repetitive element are plotted, grouped by repetitive element. These are i) DNA transposons, ii) long interspersed nuclear elements (LINEs), iii) miscellaneous classes of element (“Other”), iv) rolling-circles (RC), v) retrotransposons, vi) RNAs, vii) rRNAs, viii) Satellites, ix) segemental duplications, x) simple repeats, xi) short interspersed nuclear elements (SINEs), xii) tRNAs, and xiii) unclassified repeats (“Unknown”).

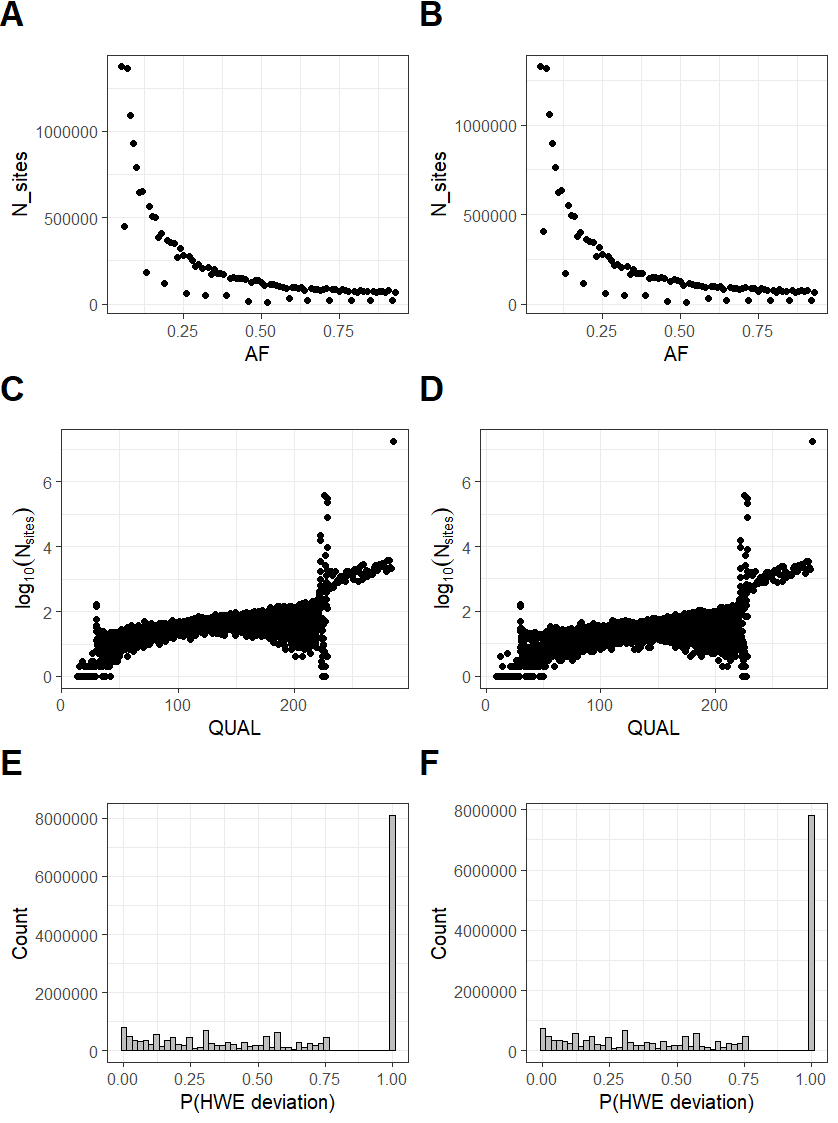

**Supplementary Figure 8 – SNP calls from linear reference genome vs. pangenome**

For SNPs called using linear reference genome (column 1: panels A, C, E) and the full pangenome (column 2, panels B, D, F), i) the frequency of SNPs with varying levels of allele frequency (AF; panels A + B), ii) the log_10_(frequency) of QUAL scores (panels C and D), and iii) the frequency of sites with a given probability of deviating from Hardy-Weinburg equilibrium (HWE; panels E and F).

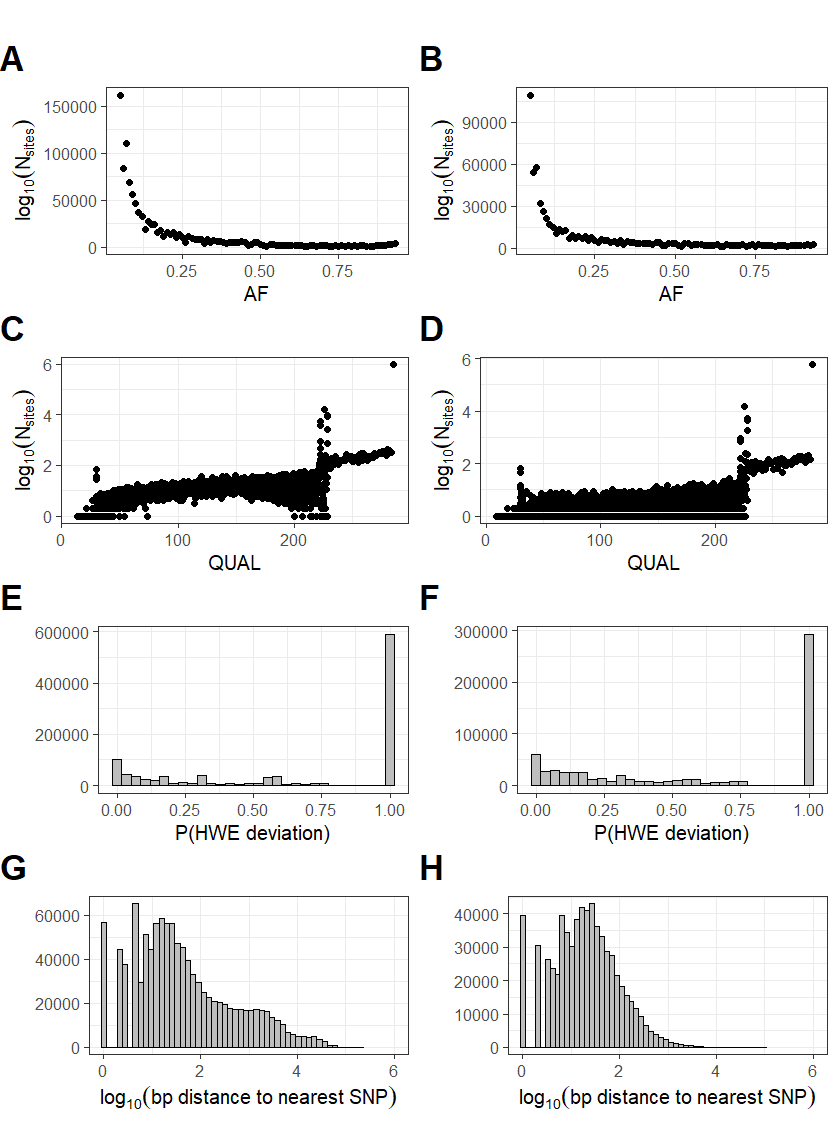

**Supplementary Figure 9 – SNPs called exclusively from linear reference genome vs. exclusively from pangenome**

For SNPs called exclusively when using linear reference genome (column 1, panels A, C, E and G), and called exclusively when using the pangenome (column 2, panels B, D, F and H), i) the frequency of SNPs with varying levels of allele frequency (AF; panels A and B), ii) the log_10_(frequency) of QUAL scores (panels C and D), and iii) the frequency of sites with a given probability of deviating from Hardy-Weinburg equilibrium (HWE; panels E and F). The fourth panels indicate the log_10_(basepair distance) of each SNP, to the closest SNP in the alternate callset (panels G and H).

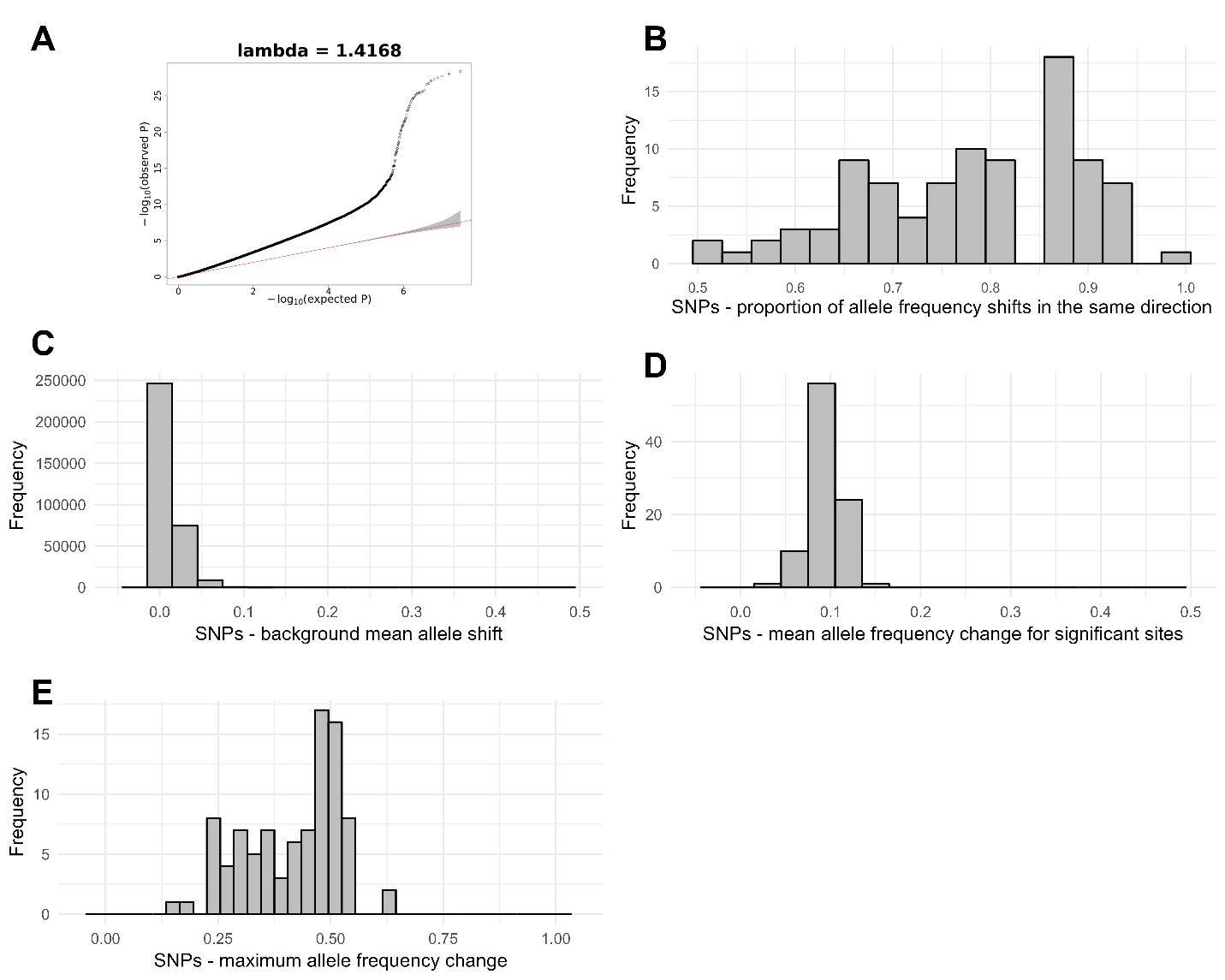

**Supplementary Figure 10 - Cochran-Mantel-Haenszel test statistics for SNPs**

A) quantile-quantile plots for -log_10_(p-values) from Cochran-Mantel Hahn (CMH) tests from SNP allele frequencies, as implemented in the GWASTools package. Genomic inflation factor (lambda) is indicated at the top of each plot. B) Histogram indicating the proportion of allele frequency shifts occurring in the most frequent direction between healthy and unhealthy pools, excluding pairs of pools where the allele frequency shift is 0. C) For a random 1% of SNPs considered in the analysis, histogram of the mean absolute value of allele frequency shift between unhealthy and healthy pools. D + E) For SNPs identified as significant using the CMH test, histogram of the D) mean and E) maximum absolute value of allele frequency shift between unhealthy and healthy pools.

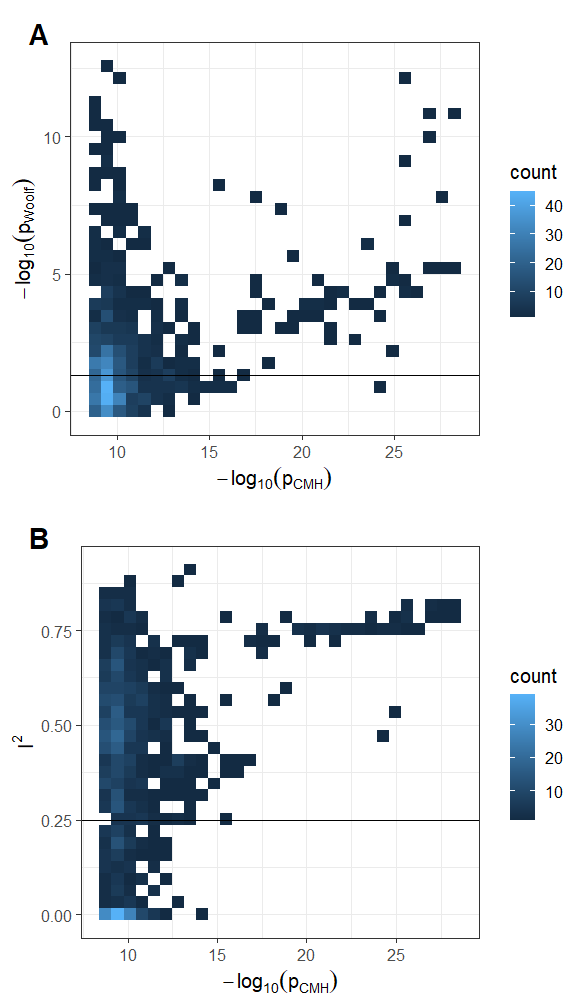

**Supplementary Figure 11 – heterogeneity tests for SNPs with significant p-values from Cochran-Mantel Haenszel test**

For the 866 SNPs with significant p-values from the Cochran-Mantel-Haenszel (CMH) test) after Bonferroni correction (ɑ = 0.05) , -log_10_(p-values) from the CMH test are plotted on a heatmap against A) -log_10_(p-values) from a Woolf test, and B) values of the I^2^ statistic. Horizontal lines represent thresholds for significant heterogeneity; p < 0.05 for the Woolf test, and I^2^ > 0.25.

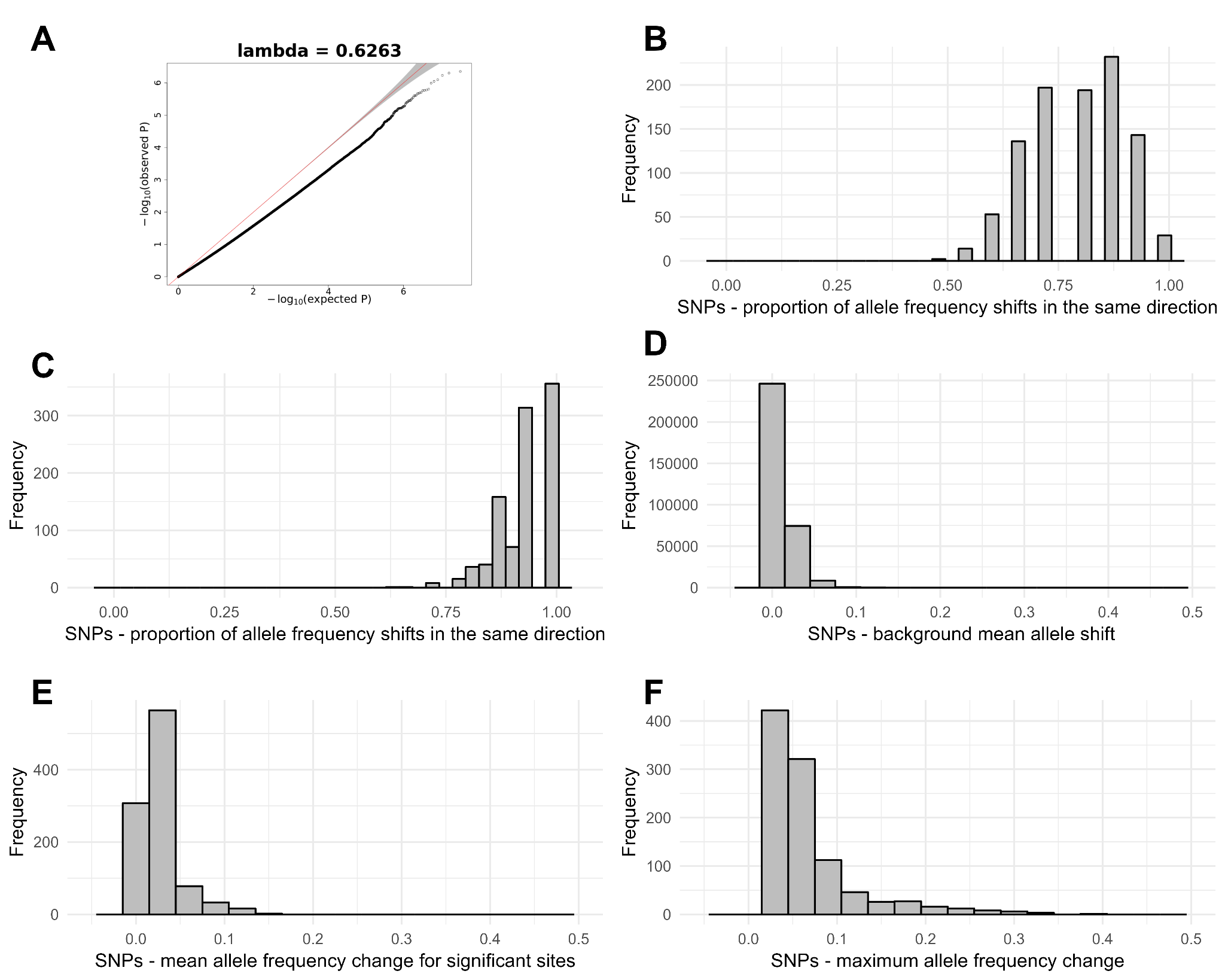

**Supplementary Figure 12 - quasi-binomial generalized linear model test statistics for SNPs**

A) quantile-quantile plots for -log_10_(p-values) from quasibinomial generalized-linear model (qGLM) tests from SNP allele frequencies. Genomic inflation factor (lambda) is indicated at the top of each plot. B) Histogram indicating the proportion of allele frequency shifts occurring in the most frequent direction between healthy and unhealthy pools, including pairs of pools where the allele frequency shift is 0. C) Histogram indicating the proportion of allele frequency shifts occurring in the most frequent direction between healthy and unhealthy pools, excluding pairs of pools where the allele frequency shift is 0 D) For a random 1% of SNPs considered in the analysis, histogram of the mean absolute value of allele frequency shift between unhealthy and healthy pools. E + F) For the 1,000 SNPs with the smallest p-values (none of which were significant), histograms of the E) mean and F) maximum absolute value of allele frequency shift between unhealthy and healthy pools.

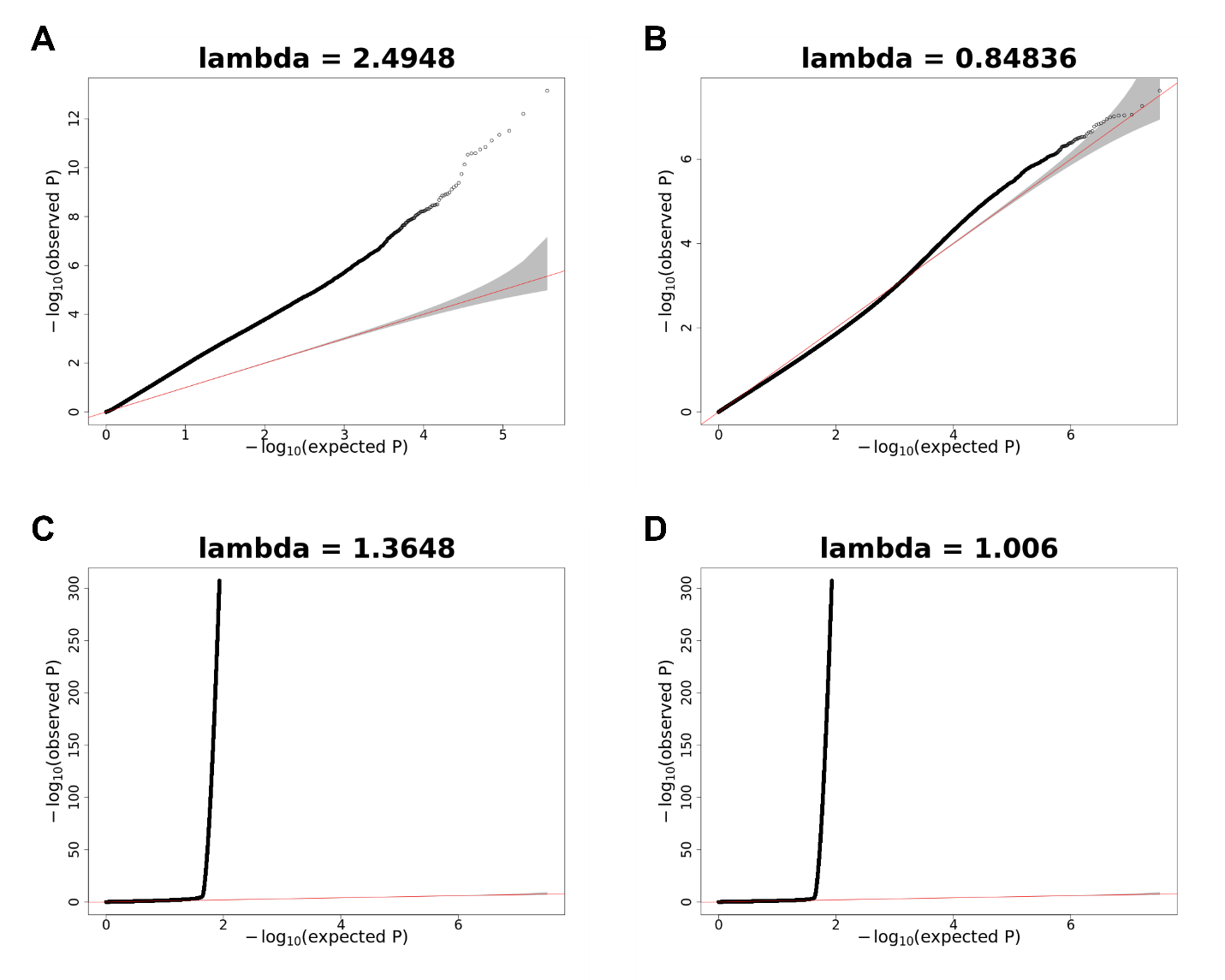

**Supplementary Figure 13 – quantile-quantile plots for quasibinomial generalized linear model tests**

Quantile-quantile plots for -log_10_(p-values) from generalized-linear model (GLM) tests from SNP allele frequencies. Genomic inflation factor (lambda) is indicated at the top of each plot. Panels indicate A) using seed source as a fixed effect in a quasibinomial GLM, B) using the first two principal components from principal component analysis of SNPs from the poolseq data as a fixed effect in a quasibinomial GLM, C) Using seed source as a random effect in a binomial GLM, and D) Using seed source and individual pool identity as random effects in a binomial GLM.

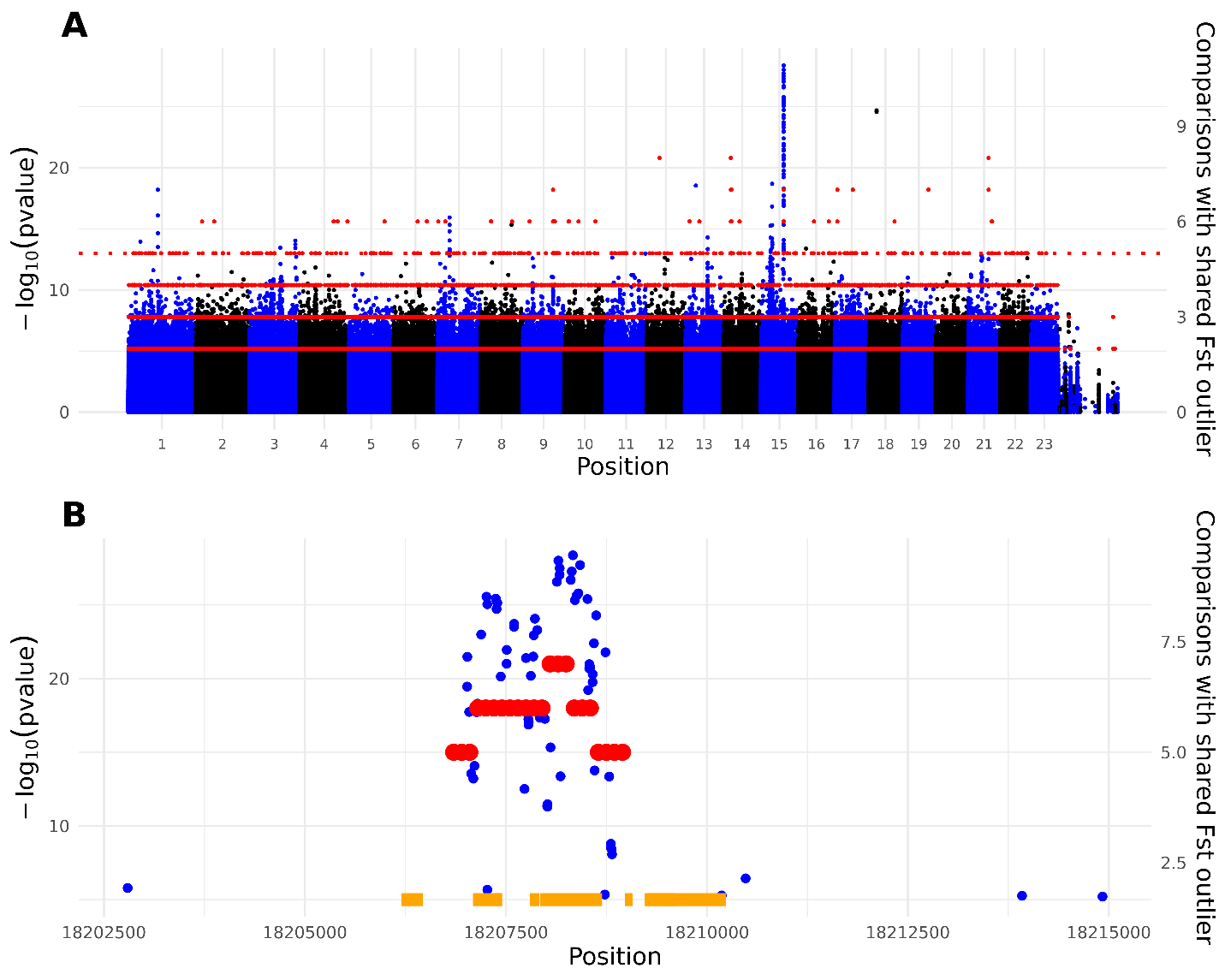

**Supplementary Figure 14 – Putative outlier region on chromosome 15**

A) Association of SNPs called from poolseq data from Stocks et al. (2019), with ash dieback disease damage. Sites are ordered by position along scaffolds in the BATG-1.0 genome assembly. Labels for the 23 largest scaffolds are shown. Black and blue coloured points represent the -log_10_ p-values of a Cochran-Mantel Haenszel (CMH) test, with alternating colours representing the different scaffolds and values indicated by the left hand Y axis. Red points indicate the number of 500bp sliding windows sharing an F_st_ window in the top 1% of values for each chromosome, shared by a minimum number of comparisons as indicated on the right hand Y axis (i.e. 5 corresponds to 5 comparisons having an Fst value in the top 1%; axes are scaled so that 5 on the right hand axis corresponds to -log_10_ (*p*) = 13 on the left hand axis). B) Figure A, zoomed in on the region on chromosome 15 representing a cluster of 55/92 CMH test values < 1 x 10^-13^. Orange bars represent annotated repeats overlapping the region.

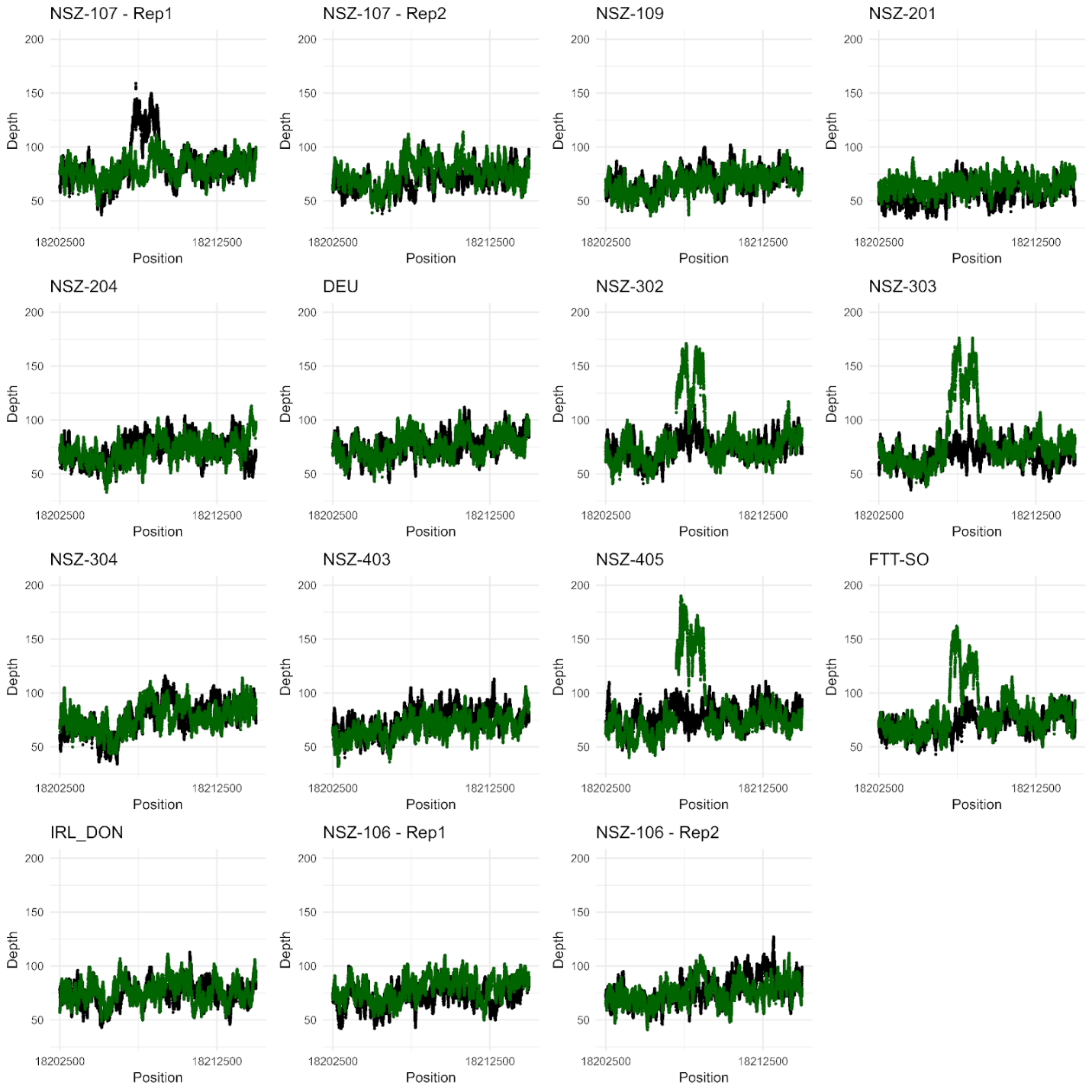

**Supplementary Figure 15 – Read depths across chromosome 15 outlier region**

Total read depths of reads for each of the 15 pairs of pools, surrounding the region with overlapping significant CMH SNPs and F_st_ outlier windows on Chromosome 15. Green points represent depths in the healthy pools; black the unhealthy pools. Biological replicates from NSZ-107 and NSZ-106 are plotted separately.

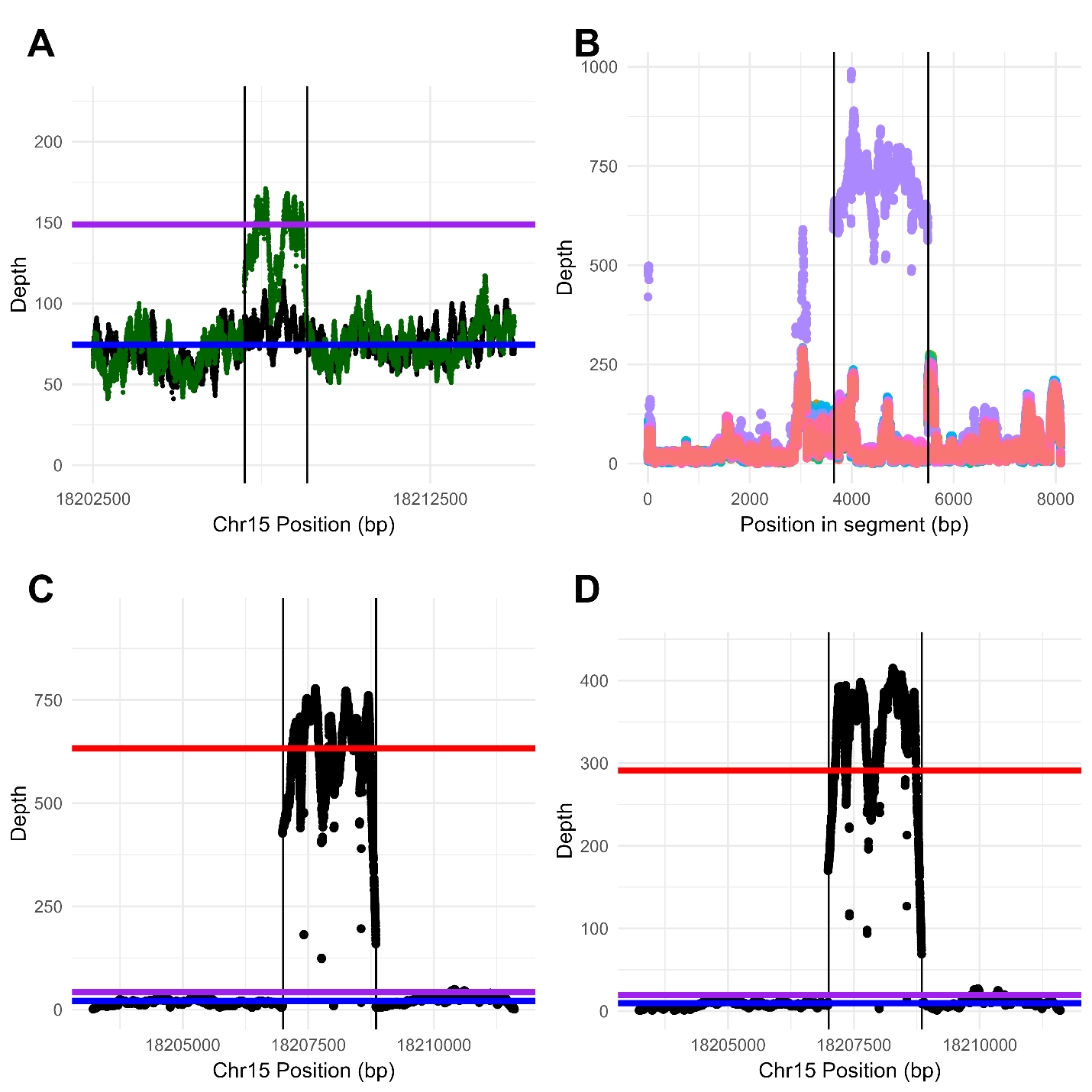

**Supplementary Figure 16 – Read depths across chromosome 15 outlier region from individually sequenced samples**

Read depth from reads mapped to the pangenome in the region of the chromosome 15 outlier region for NSZ-302 poolseq data from healthy pool (green) and unhealthy pool (black). B) Read depths from reads from 150 individually sequenced samples from Stocks et al. 2019 mapped to a segment corresponding to the high depth region in A, and 3kb upstream and downstream of this. Depth for each individual is presented in a different colour. Panels C and D represent read depths from Individual 58 from Stocks et al. (2019) (C), and Individual S178C from Metheringham et al. (2025) (D). Vertical lines represent the boundary of the outlier region. Horizontal lines in panels A, C and D represent multiple of the median depth of the flanking region outside the outlier region, for each panel (+/- 3kb) – blue = 1X, purple = 2X, red = 30X.

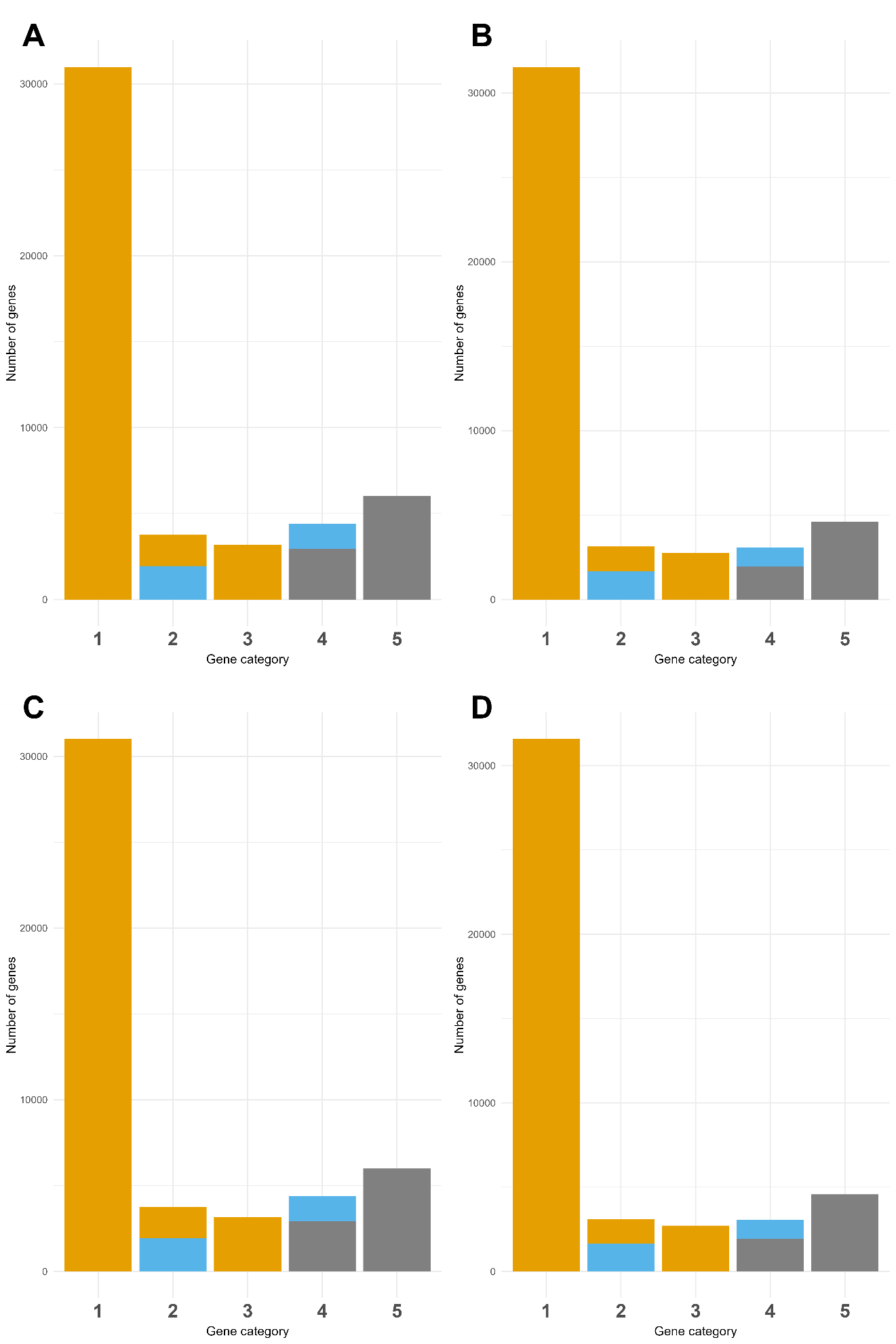

**Supplementary Figure 17 – The impact of different annotation merging approaches on genes classified as dispensable or indispensable**

Barplot of the number of genes in each category in the pangenome, where numbers correspond to genes present in the BATG-1.0 reference (1, 2 and 3) and in every other sample (1), or absent in some samples and overlapping with an SV (2), or (3) absent in some samples, but none of these overlap with an SV, or genes that are absent from the BATG-1.0 reference (4,5) and either overlap with an SV (4) or never overlap with an SV (5). Bar colours are based on reclassification of genes in each category as invariant (orange), dispensable (blue), or discarded from the analysis (grey) according to how consistently they overlap with SVs based on an F1 score. Genes in category 2 with an F1 score > 0.8 are classified as dispensable (blue); those with a score below that threshold are categorised as invariant. Genes in category 3 are never associated with an SV so are all categorised as invariant. Genes in category 4 with an F1 score > 0.8 are classified as dispensable, those with a score <= 0.8 are discarded from the analysis. Genes in category 5 are never associated with an SV so are discarded from the analysis. In an approach where genes absent in some individuals are treated as dispensable, all the genes in categories 2-5 would be treated as dispensable genes. Panels correspond to analysis performed where i) genes in BATG-1.0 that overlapped by >50% in a coding sequence on the same strand had the shortest gene removed (A and B), or not (C and D), and ii) Orthogroups producing during the annotation of the pangenome grouped together if any two members had a >50% of a coding sequence overlapping on the same strand (B and D), or not (A and C).

### Supplementary Table captions

### Supplementary Table 1 - *Fraxinus excelsior* genome assembly statistics

Tab 1.1: details of the assembly statistics for *Fraxinus excelsior* from the present and other studies, including both hapotypes from the phased linear reference genome in this study. Row names refer to i) the assembly name, ii) the source of the assembly, where applicable giving the database name and the identifier, iii) the study associated with each assembly, where applicable, iv) the size of the assembly in Mb, v) the number of contigs, vi) the N50 of the assembly in Mb (50% of the assembly is in fragments this size or larger), vii) the L90 for each assembly (90% of the sequence is present in this many contigs), ix) the number of Ns per 100 kilobases, x) the BUSCO completeness score from the eudicots_odb10 dataset, in the format “% of total complete BUSCOs identified [% single copy, % duplicated], % of fragmented BUSCOs, % missing”, xi) the Long Terminal Repeat (LTR) Assembly Index (LAI) score, xii) the number of intact LTRs, and xiii) the number of intact LTRs per megabase. Tab 1.2: For BATG-1.0 and haplotype 2, LAI score per scaffold for top 23 scaffolds by size.

### Supplementary Table 2 - Reference protein sequences downloaded for Orthofinder

Details the protein sequence data downloaded for species/assemblies used in Orthofinder analysis to identify homology of BATG-1.0/pangenome sequences. Column headings refer to i) scientific name of species used ii) common name iii) database identifier; NCBI identifiers for all except *Fraxinus excelsior* which was downloaded from the Zenodo database associated with the Sollars et al. 2017 publication, iv) the database location where the protein sequences were downloaded and v) the doi of the associated publication for each dataset.

### Supplementary Table 3 - Sample information for long-read sequencing.

Sample information for the 50 biological samples sequenced with Oxford Nanopore Technologies (ONT) long-reads and used to construct the pangenome. Column headings refer to i) sample name, ii) provenance name for the samples, iii) material source - the RAP (Realising Ash’s Potential) Trial; Forest Research (FR) Mass Screening Trial; or the FRAXIGEN Trial; wild collected samples, or the if the sample was from the reference genome plant at Royal Botanic Gardens, Kew, iv) the latitude and v) longitude of the origin of the sample - some of these coordinates are approximate, details of how these were obtained/derived are given in column vi) location information, vii) the collection date of the sample, viii) the number of read pairs for samples that had short-read RNA-seq data generated (NA = no data generated), ix) the mean DNA coverage from ONT dats (assuming a genome size of 840Mb) after read mapping to BATG-1.0 (see Methods).

### Supplementary Table 4 - GO enrichment tables

Details of the results of GO term enrichment analysis of gene sets relative to a background set, using topGO. Column headings refer to i) GO identifier, ii) GO term description, iii) Background - the number of genes in the background set annotated with this GO term, iv) Observed - the number of genes in the set of interest with this annotation, v) Expected - the number of genes expected in the set of interest assuming no enrichment, vi) classicFisher - the unadjusted p-value of the classic Fisher test for enrichment, vii) adjusted - the p-value adjusted using Bonferroni correction. Tabs refer to the different gene sets/background gene sets tested; 4.1 - enrichment of genes identified as dispensable after filtering, against a background of all genes included in the filtered pangenome (dispensable plus indispensable), 4.2 the putatively dispensable genes excluded during filtering for not being consistently associated with an SV, against a background of all the genes included and excluded, 4.3 the truly dispensable genes, against a background of the truly dispensable genes plus the putatively dispensable genes excluded during filtering for not being consistently associated with an SV, 4.4 genes within 10kb of the 220 SNPs associated with low ADB susceptibility, against a background of all genes included in the pangenome, 4.5 genes overlapping the 220 SNPs associated with low ADB susceptibility, against a background of all genes included in the pangenome.

### Supplementary Table 5 – Functional annotation of key subsets of genes

Spreadsheet of functional annotation of key subsets of genes, each subset in a different tab as follows – i) high-confidence dispensable genes (an SV-Gene F1 association score > 0.8; see Results; tab 5.1) annotated with a “defense response” GO term or a child term of this GO term (GO:0006952), ii) genes that contained one or more of the 220 SNPs associated with low ADB susceptibility (tab 5.2), genes within 10kb of the 220 SNPs associated with low ADB susceptibility (tab 5.3), and the subset of these genes annotated with a “defense response” GO term, as defined above (tab 5.4). Columns indicate i) gene name, ii) the bitscore and iv) p-value of the top hit in the UniProt/SwissProt database for each gene, v) the bitscore and vi) p-value of the top hit in the NCBI NR database for each gene, vi) the PANTHER superfamily classification code and vii) description for each gene, vii) the InterPro family classification code and viii) description for each gene, ix) the bitscore and x) p-value of the best hit from a DIAMOND blastp search of the genes against the Araport 11 representative gene peptides, and xi) the name of the best hit.

### Supplementary Table 6 – SNP calling statistics using linear reference genome vs. pangenome

|  | Linear reference | Pangenome | Linear reference only | Pangenome only |
| --- | --- | --- | --- | --- |
| Total SNPs called | 18,955,565 | 18,540,348 | 1,057,326 | 642,020 |
| Transition/Transversion Ratio | 2.2 | 2.18 | 2.5 | 2.13 |
| Mean proportion missing | 1.32% | 1.26% | 3.30% | 3.90% |
| Mean AF | 0.269 | 0.273 | 0.187 | 0.231 |
| % sites with QUAL < 284.5 (maximum) | 6.44% | 6.15% | 8.40% | 6.52% |
| % sites with p(HWE) < 1 | 57.30% | 57.80% | 44.10% | 54.30% |

Table outlining SNP calling statistics for reads from 42 short-read sequenced samples, called i) using a the BATG-1.0 linear reference genome, ii) using the pangenome, iii) called only using the BATG-1.0 linear reference genome, and iv) called only using the pangenome. Statistics include a) the total SNPs called, b) the transition/transversion ratio, c) the mean proportion missing, d) the mean allele frequency of SNPs called, e) the percentage of sites with the SNP quality score being less than the maximum value (284.5), and e) the percentage of sites with a p(deviation from HWE) < 1.

### Supplementary Table 7 - Comparison of key statistics with other recent pangenome studies.

Results from the present study are compared with those from plant pangenome studies published since 2022 (non-exhaustive) - pangenome studies using primarily short-read data, or incorporating multiple species, were excluded. Columns give i) the name of the study used in the labels for Figure S10, comprising the common organism name and publication year, ii) the doi of the study, iii) the first author, iv) the publication date, v) the study organism, vi) N_samples, the number of samples used, vii) N_SVs, the number of SVs detected, where provided, viii) Extra sequence (Mb), the additional sequence in megabases incorporated as part of the pangenome, where provided, ix) the linear reference genome size, where provided, x) Extra/Ref, the proportion of additional sequence as a function of the reference genome size, xi) the number of core (non-dispensable) genes reported, xii) the total number of genes reported, xiii), the number of dispensable genes (non-core) as a proportion of the total number of genes. Columns vi) vs. vii), vi) vs.x) and vi) vs. xiii) are plotted in Extended Data Figure 9 panels A, B and C respectively. Ash_2026 (unfiltered) refers to the proportion of dispensable genes prior to filtering in the present study, Ash_2026 (filtered) refers to the proportion after filtering for consistent association with SVs - see Results for further details.

### Supplementary Table 8 – RNA-seq data used to annotate BATG-1.0 assembly

Details of RNA-seq data used to annotate linear reference assembly BATG-1.0. Columns represent i) the NCBI/ENA accession data was downloaded from, where applicable, ii) the RNA sequence length (long or short reads), iii) the study associated with the data, where applicable, iv) the genotype of the individual, v) the tissue type, vi) experimental treatment (where applicable) and vii) notes.

### Supplementary File Captions

**Supplementary File 1.zip**

File containing genomic feature plots for each of the top 23 scaffolds by size.

For each of the 23 images contained in the zipped folder, chromosome number and scaffold are given at the top. Each panel represents a feature plotted by density per 100kb (Y axis), along the chromosome (x axis). Panels are, from top to bottom, i) genes (blue), ii) insertions (orange), iii) deletions (red), iv) inversions (purple), v) tandem duplications (green) and vi) repeats (brown).
