## Supplementary figures and images for "European ash pangenome reveals widespread structural variation and genetic basis of low ash dieback susceptibility"

### Chromosome_1.png

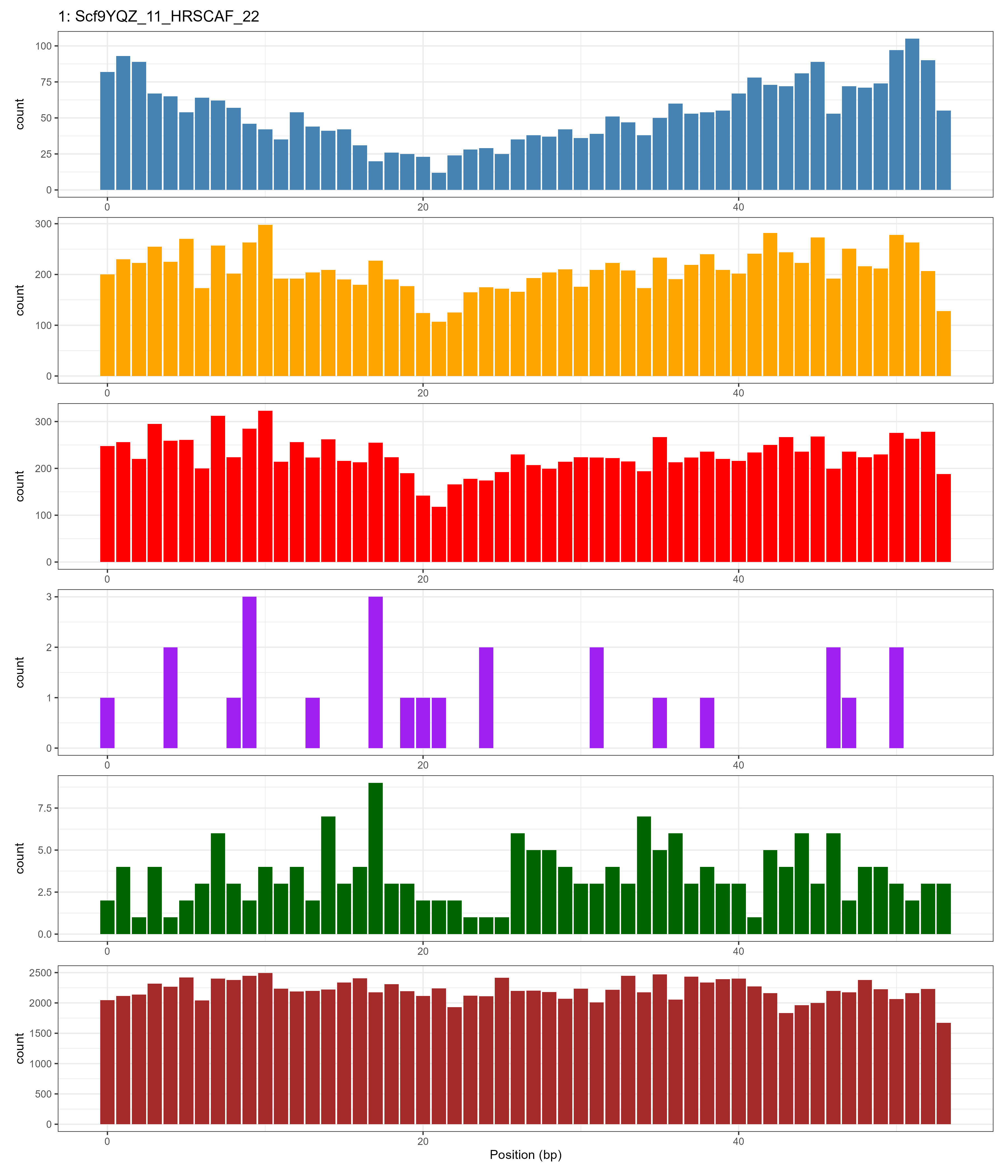

### Chromosome_2.png

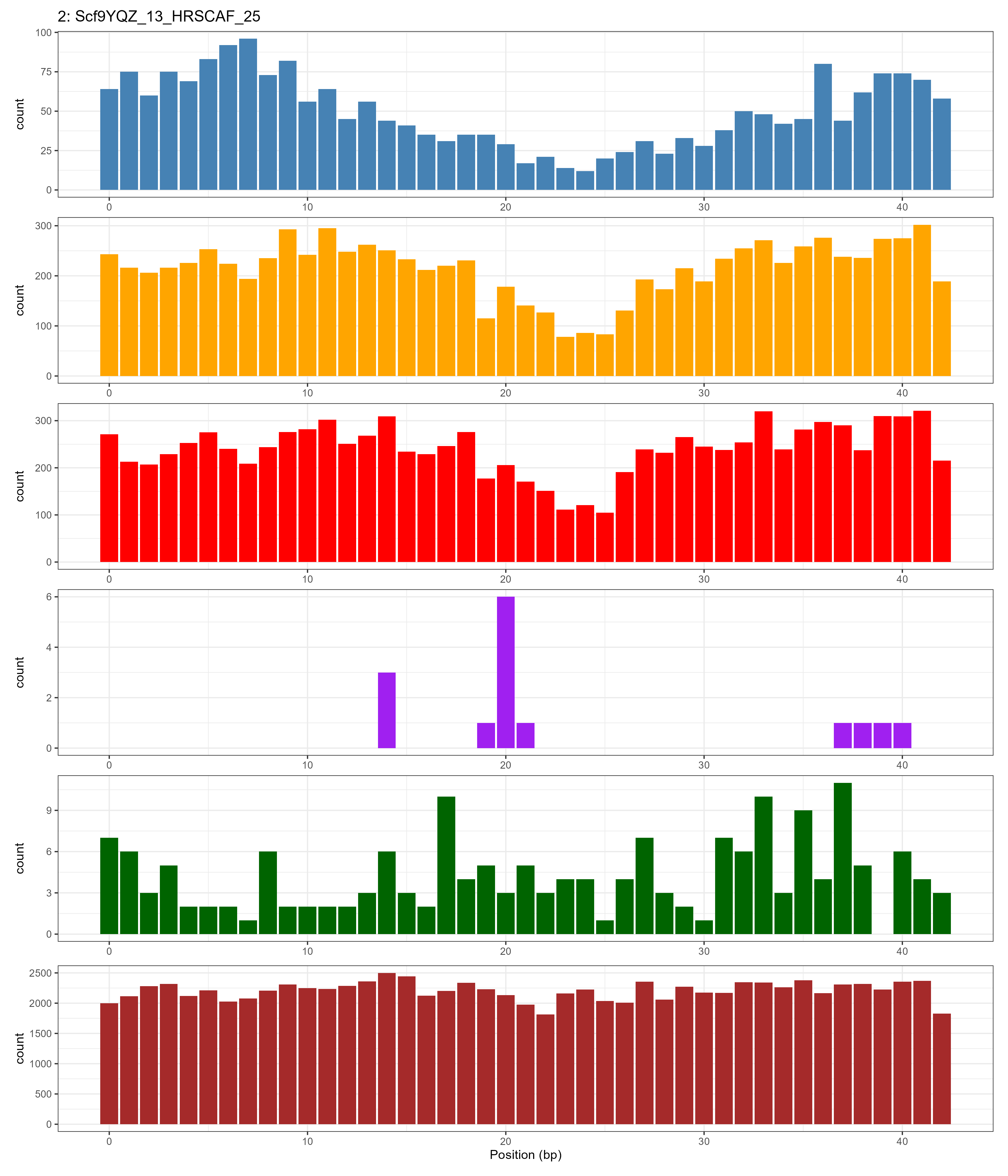

### Chromosome_3.png

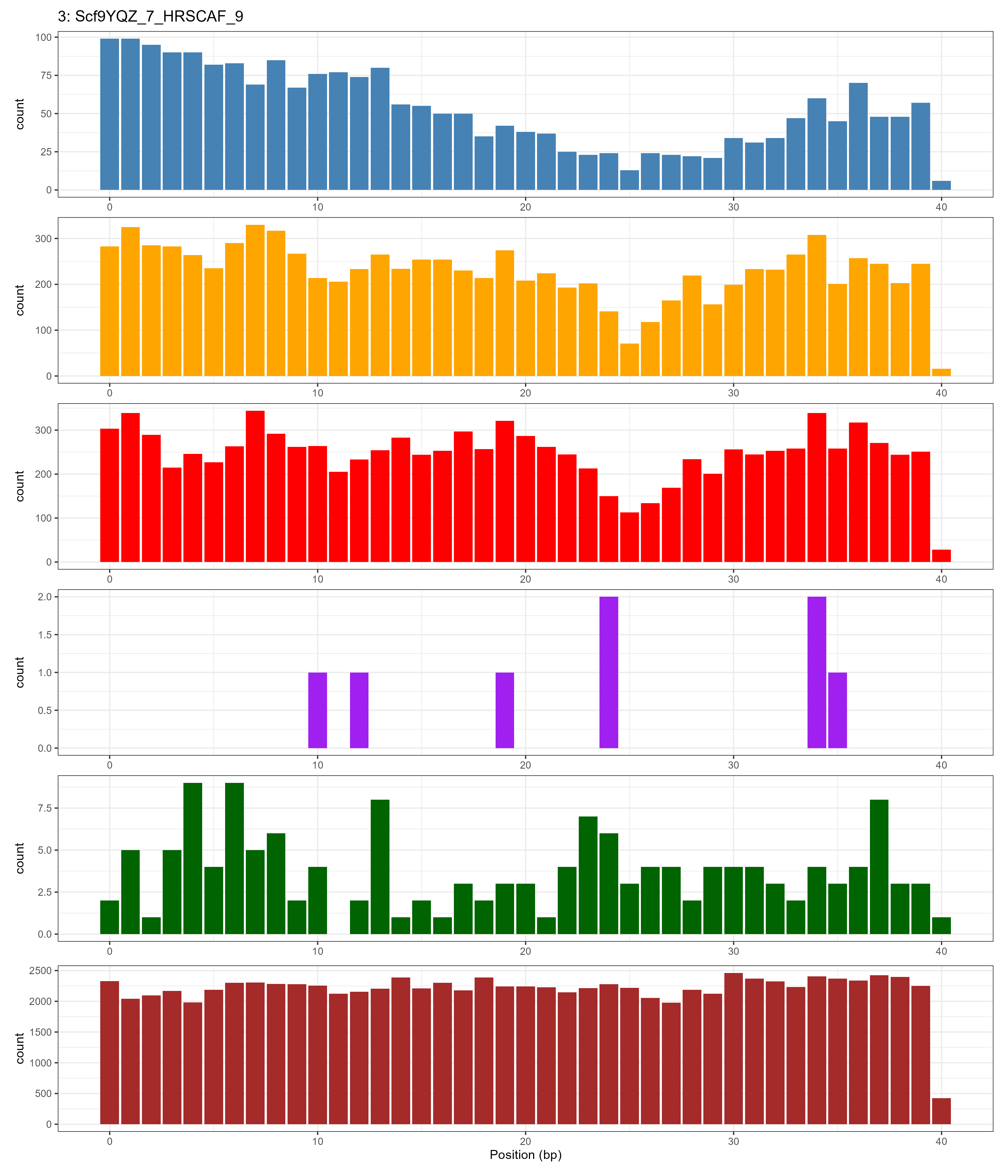

### Chromosome_4.png

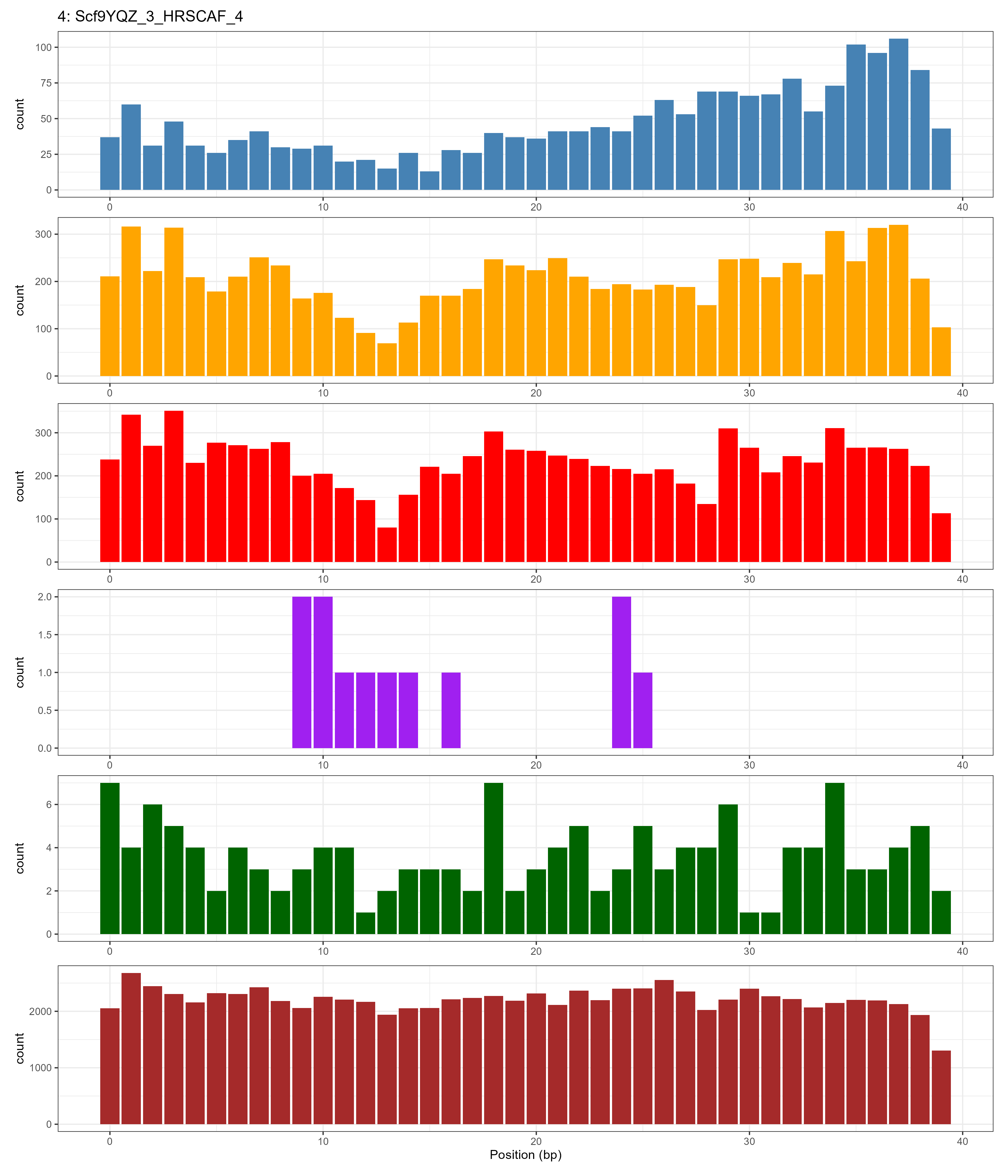

### Chromosome_5.png

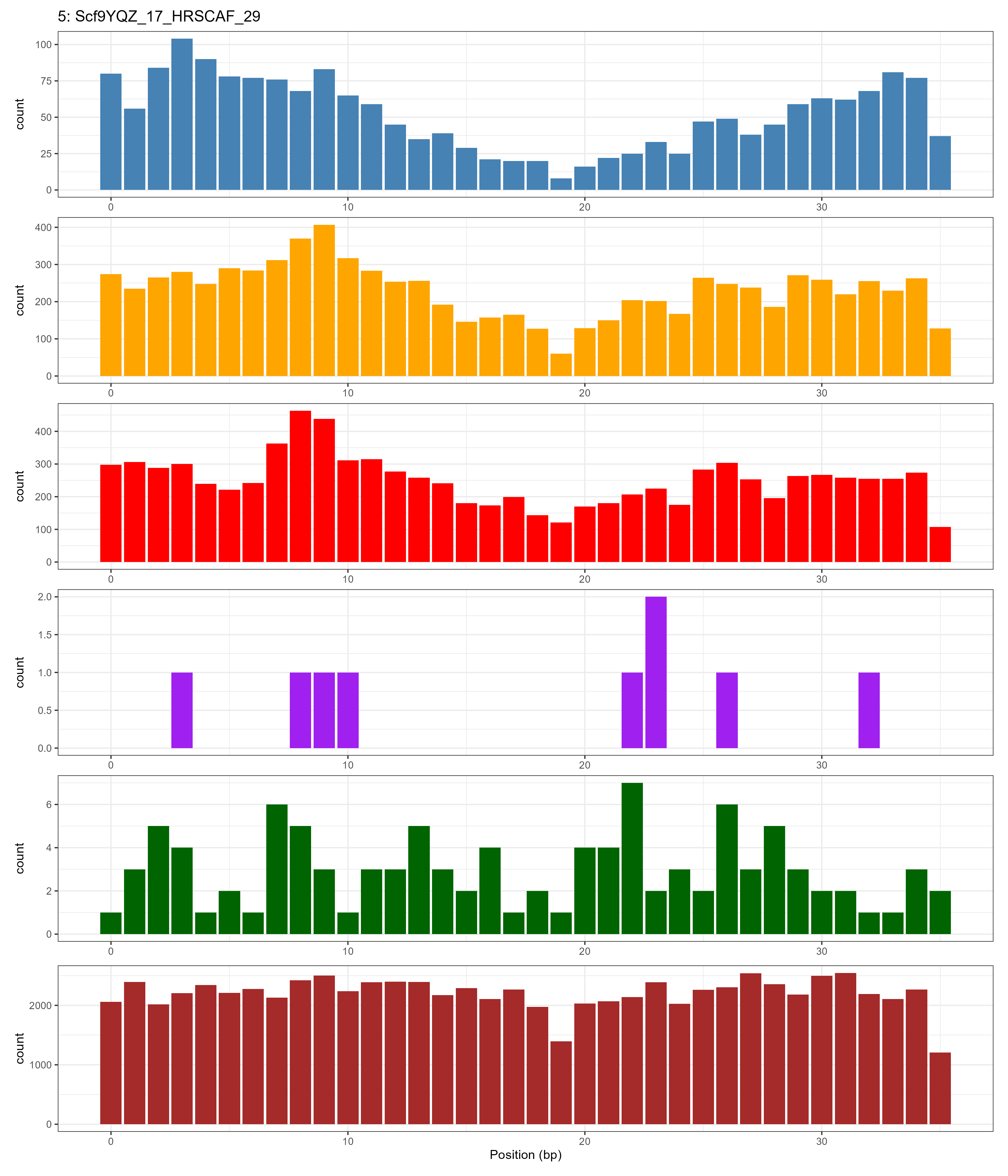

### Chromosome_6.png

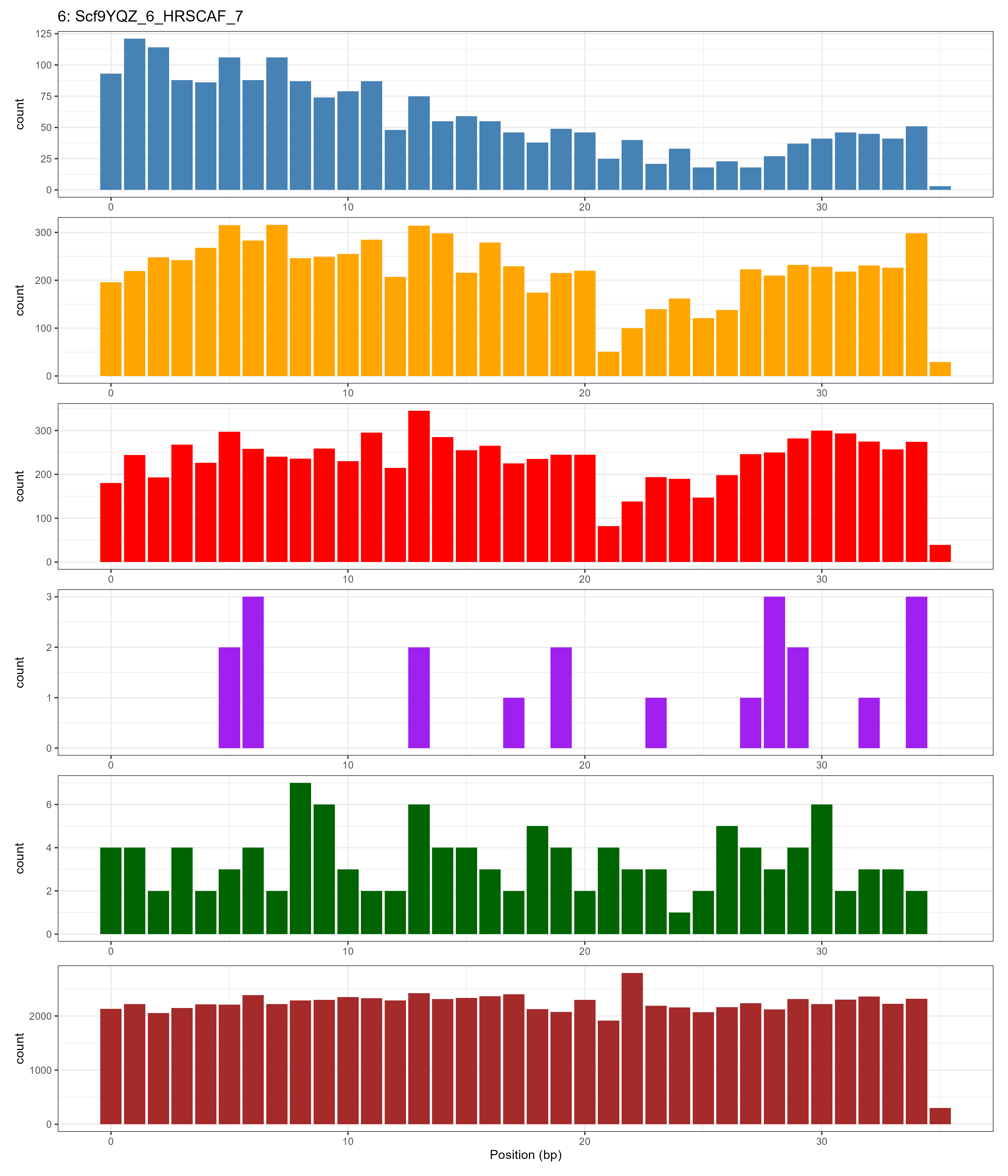

### Chromosome_7.png

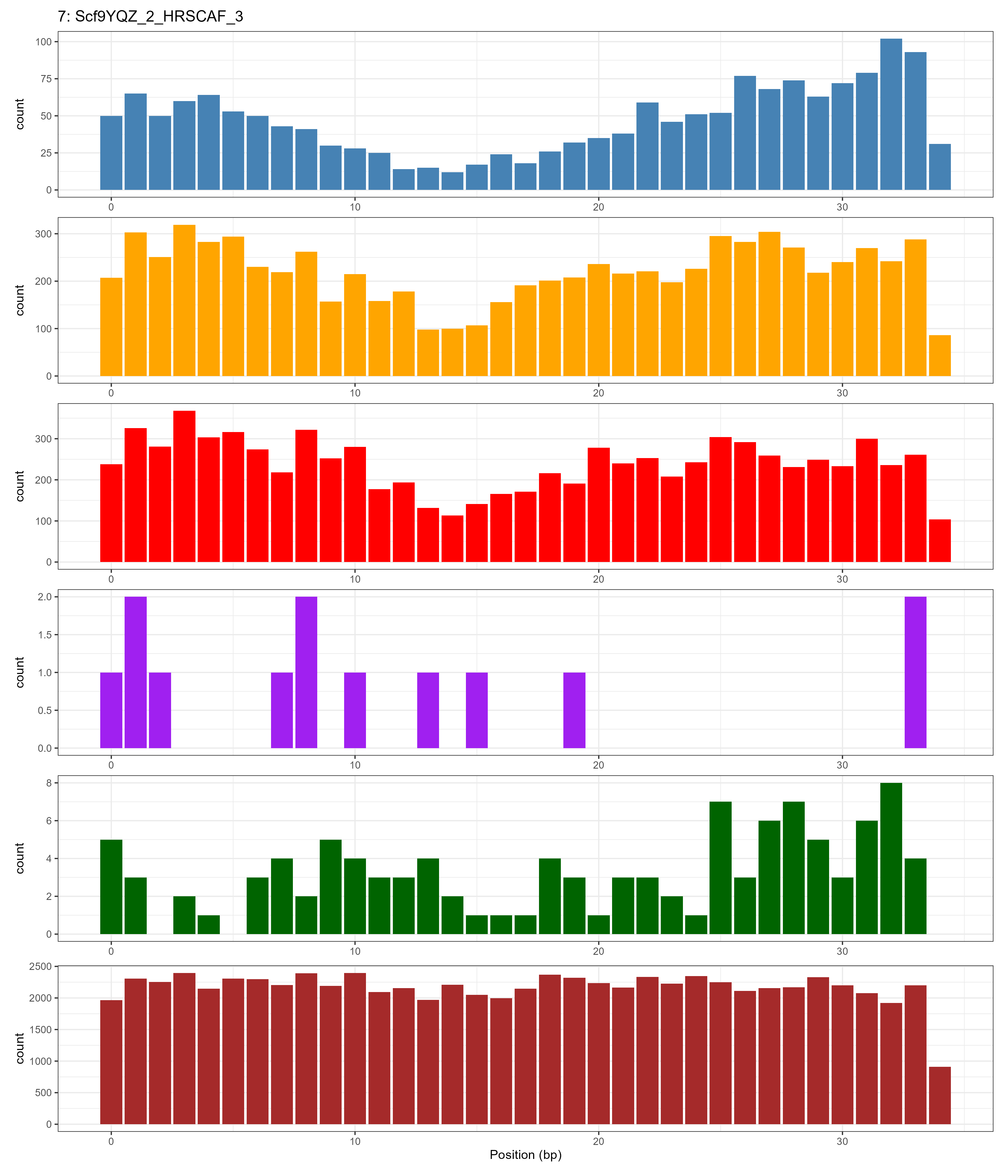

### Chromosome_8.png

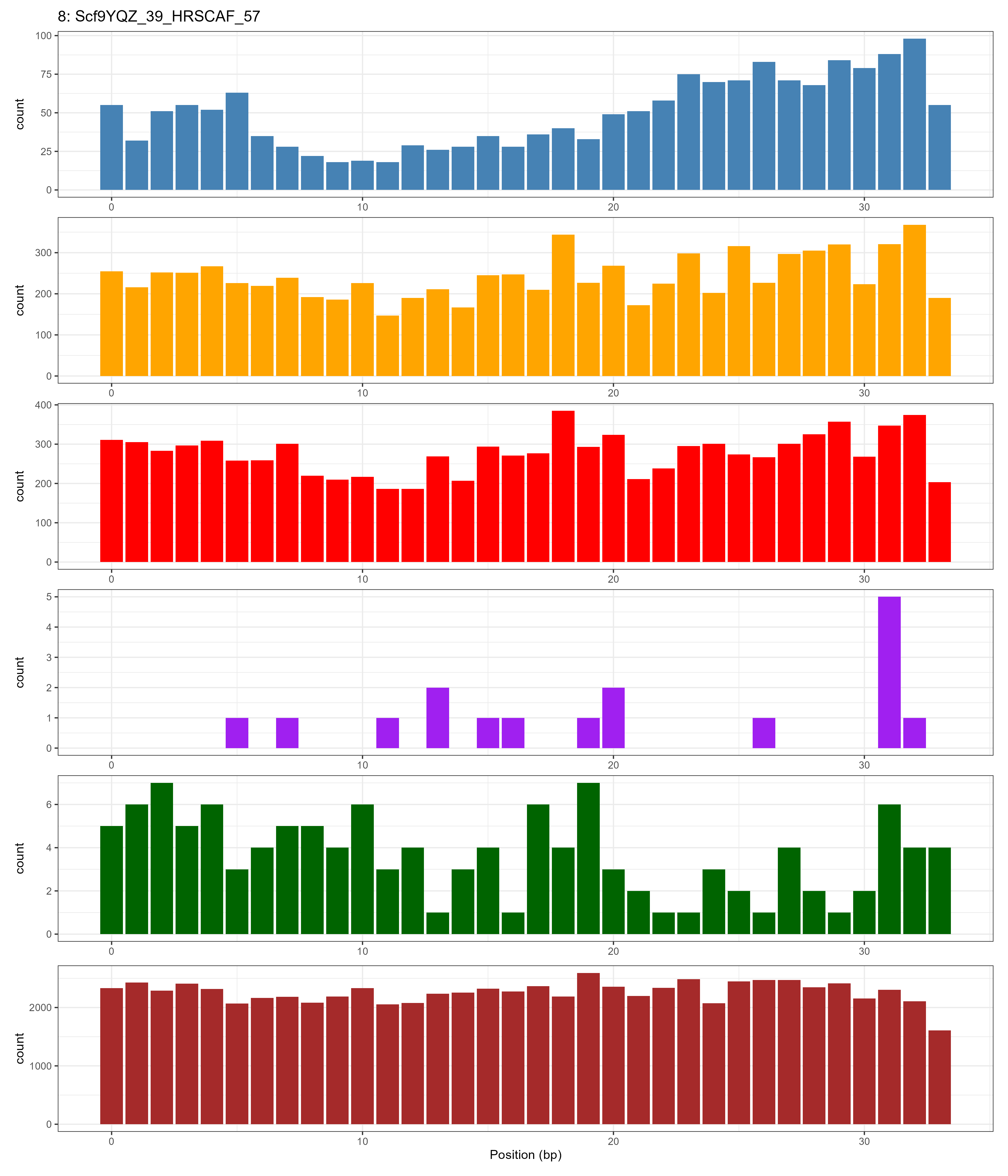

### Chromosome_9.png

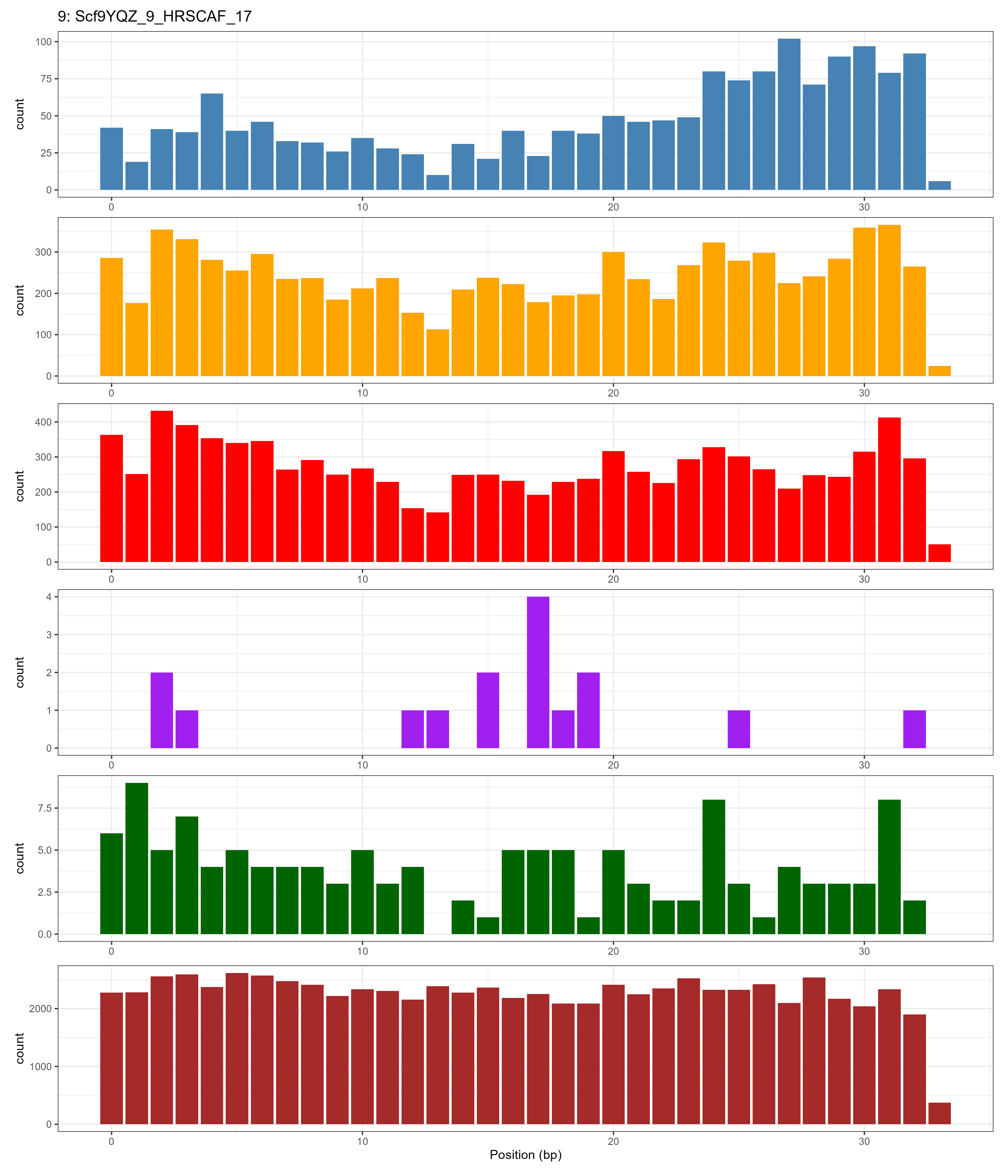

### Chromosome_10.png

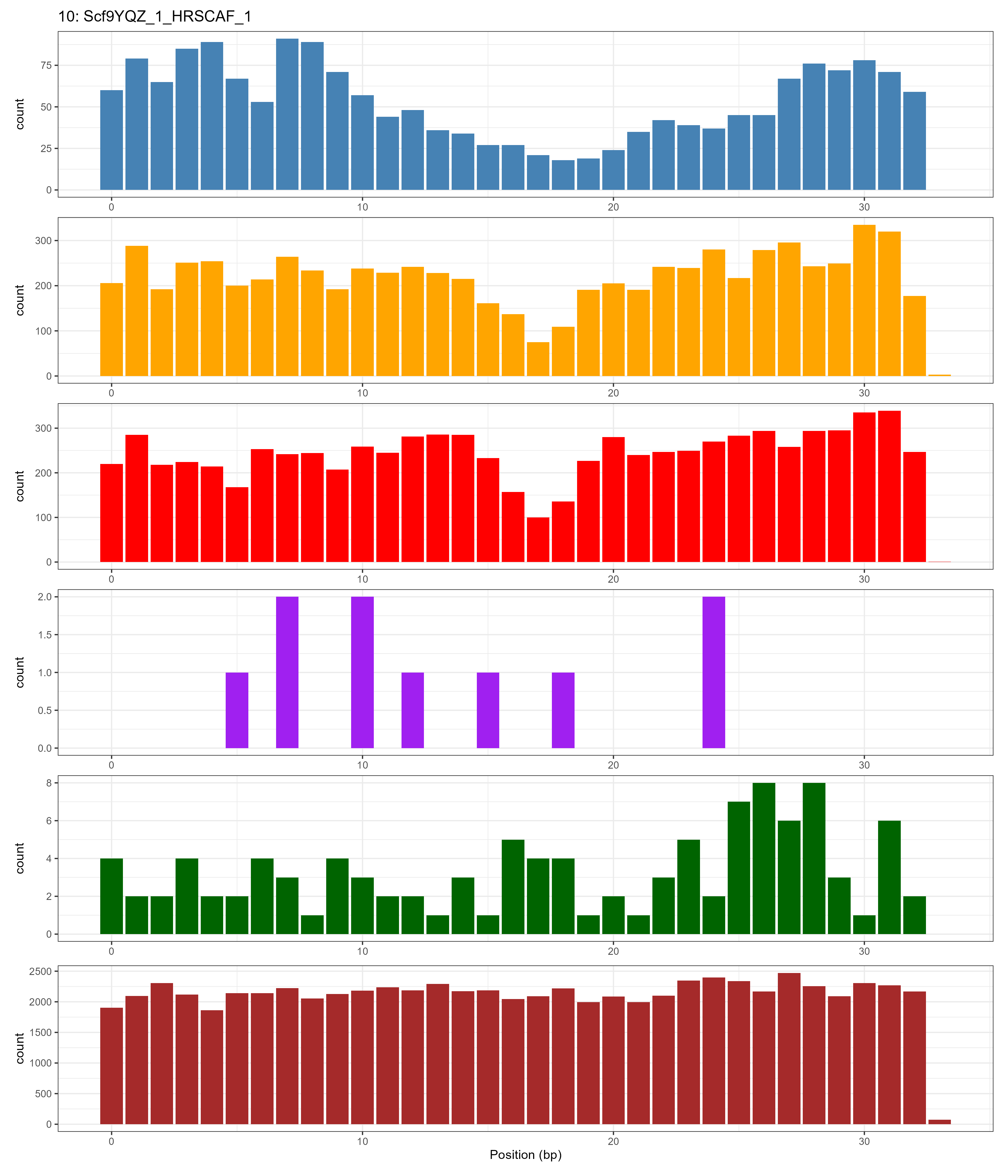

### Chromosome_11.png

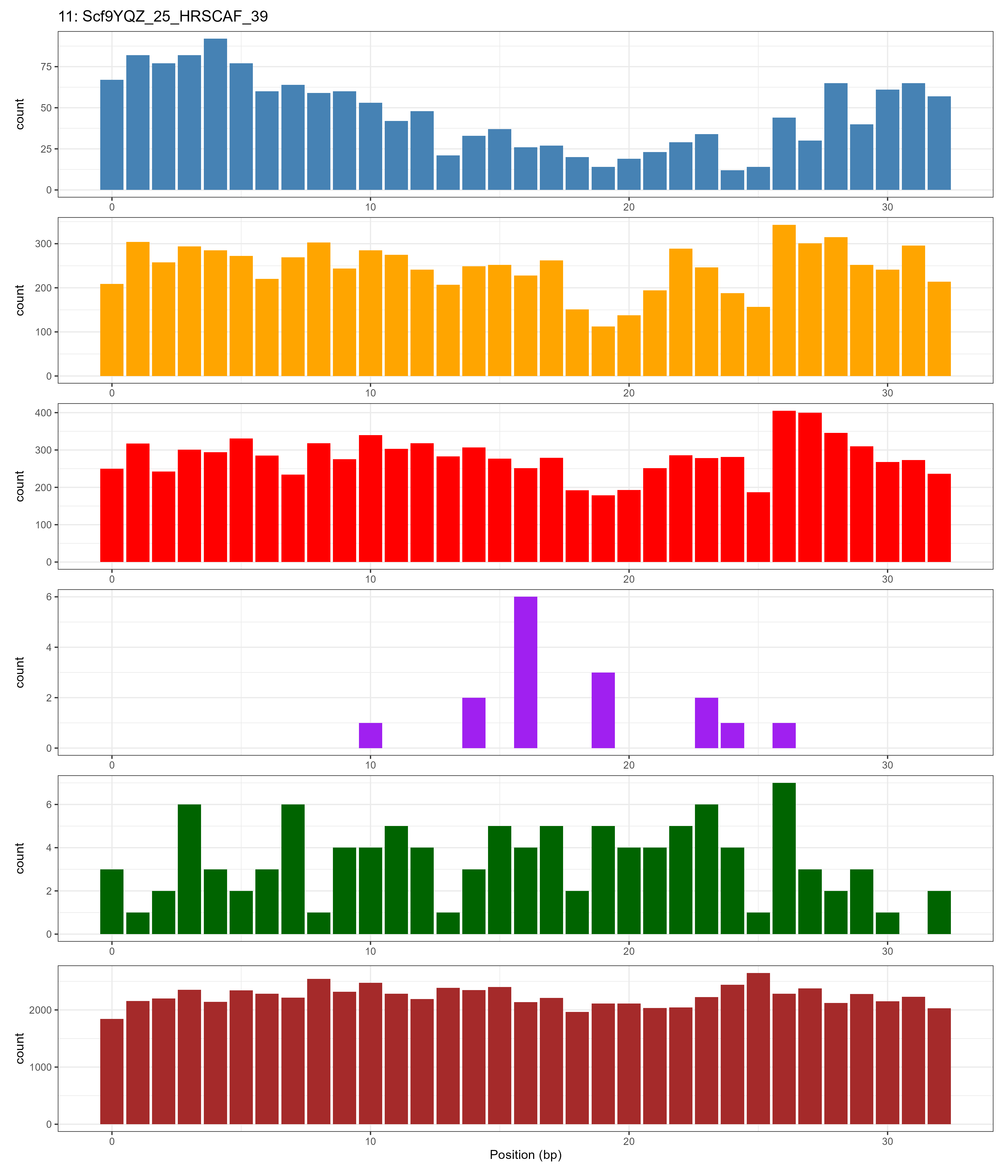

### Chromosome_12.png

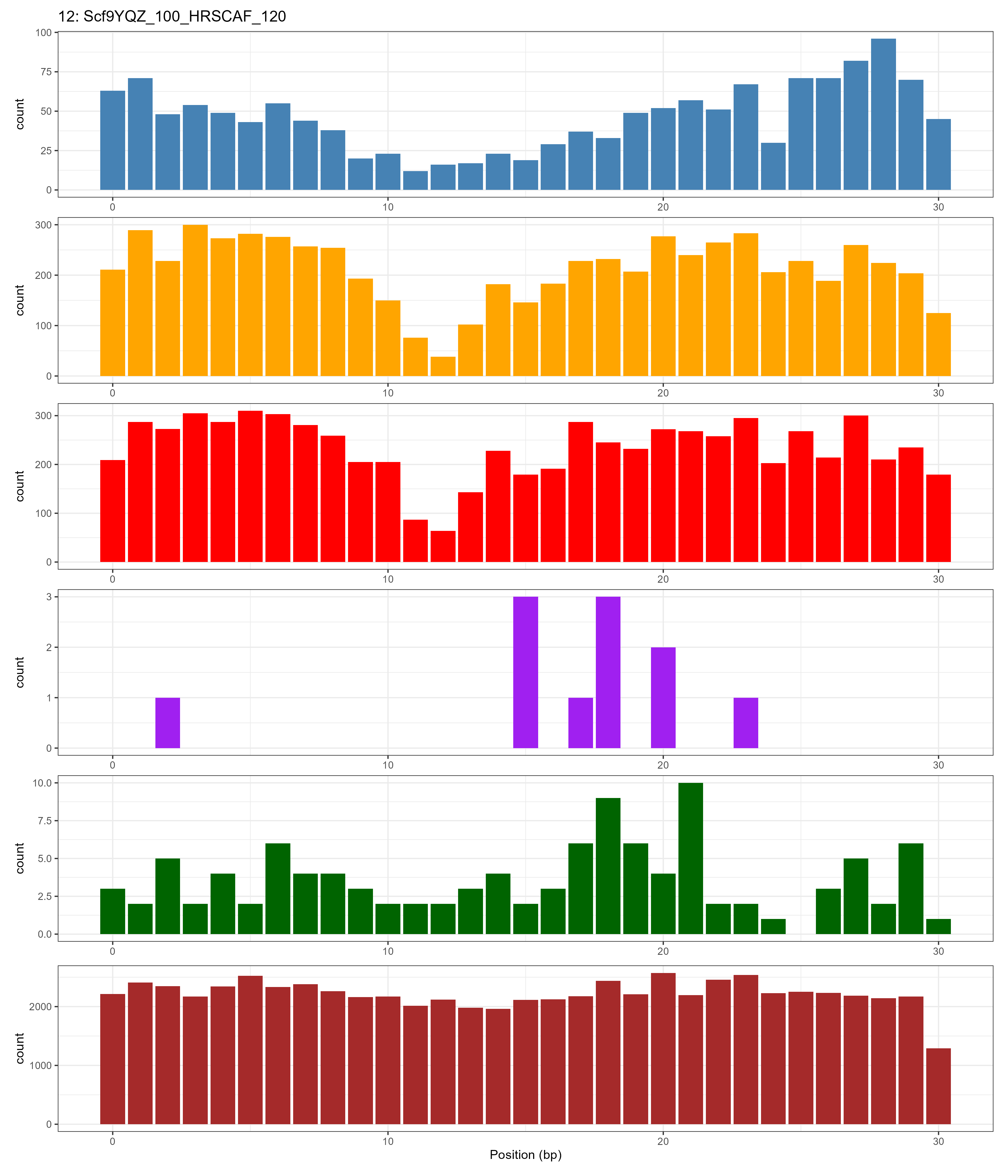

### Chromosome_13.png

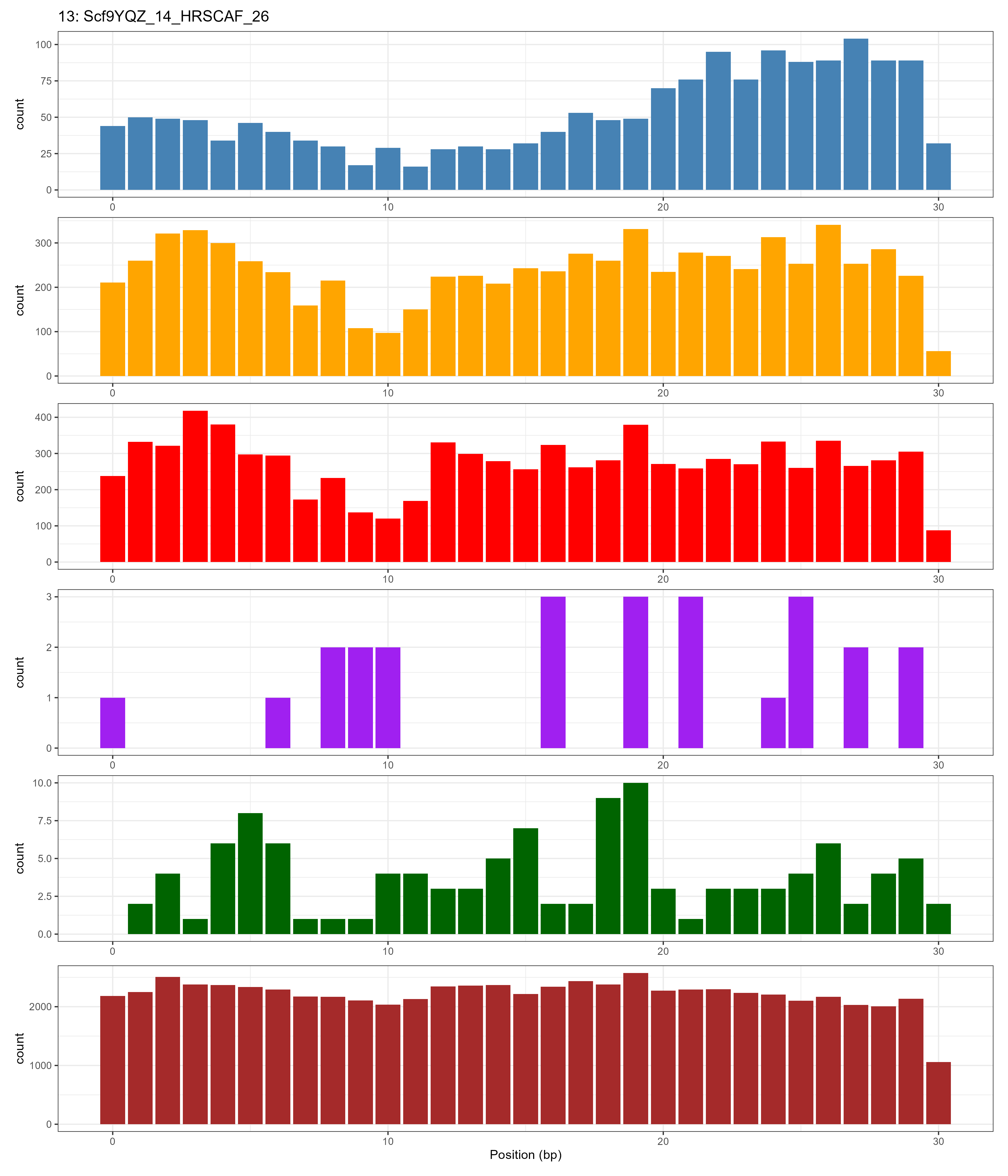

### Chromosome_14.png

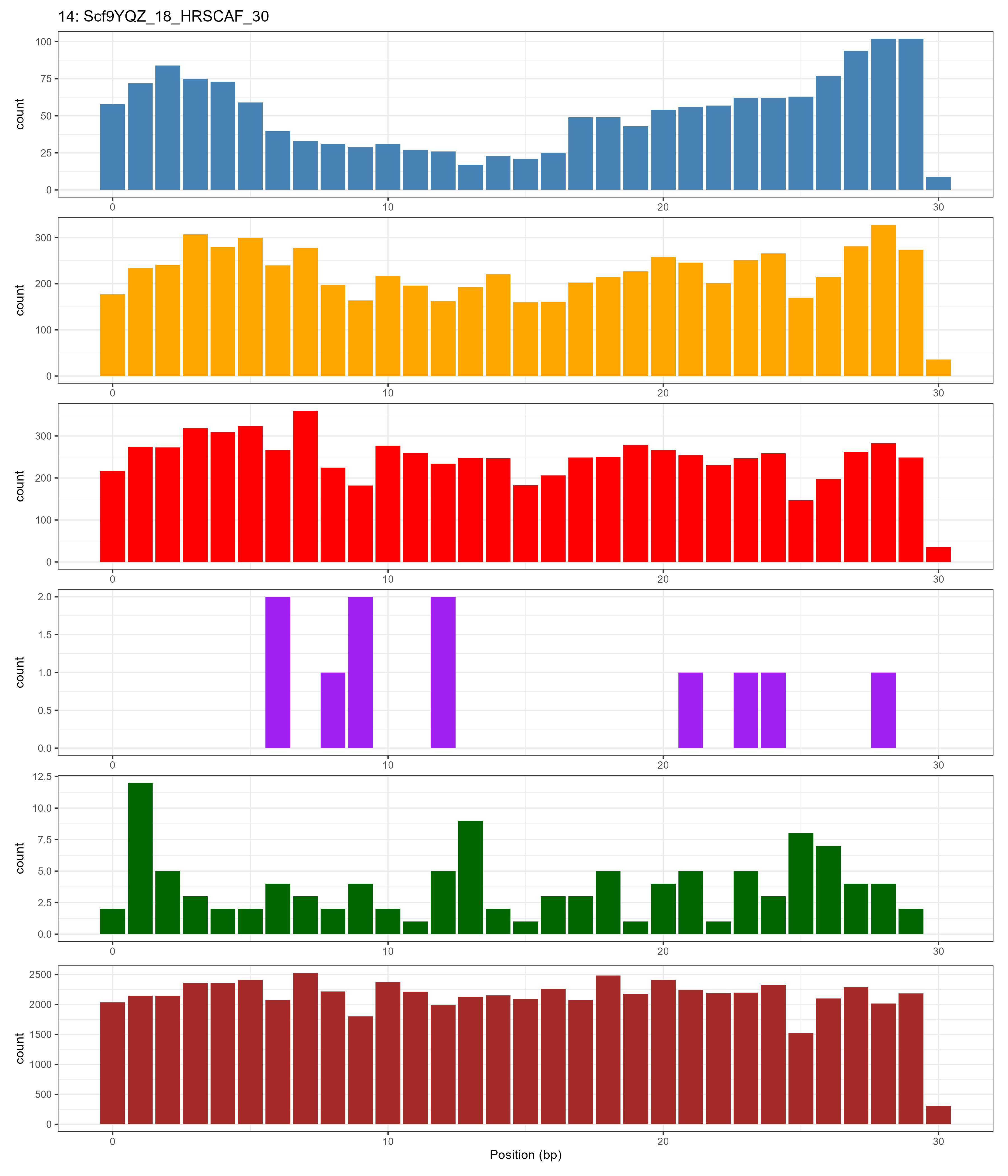

### Chromosome_15.png

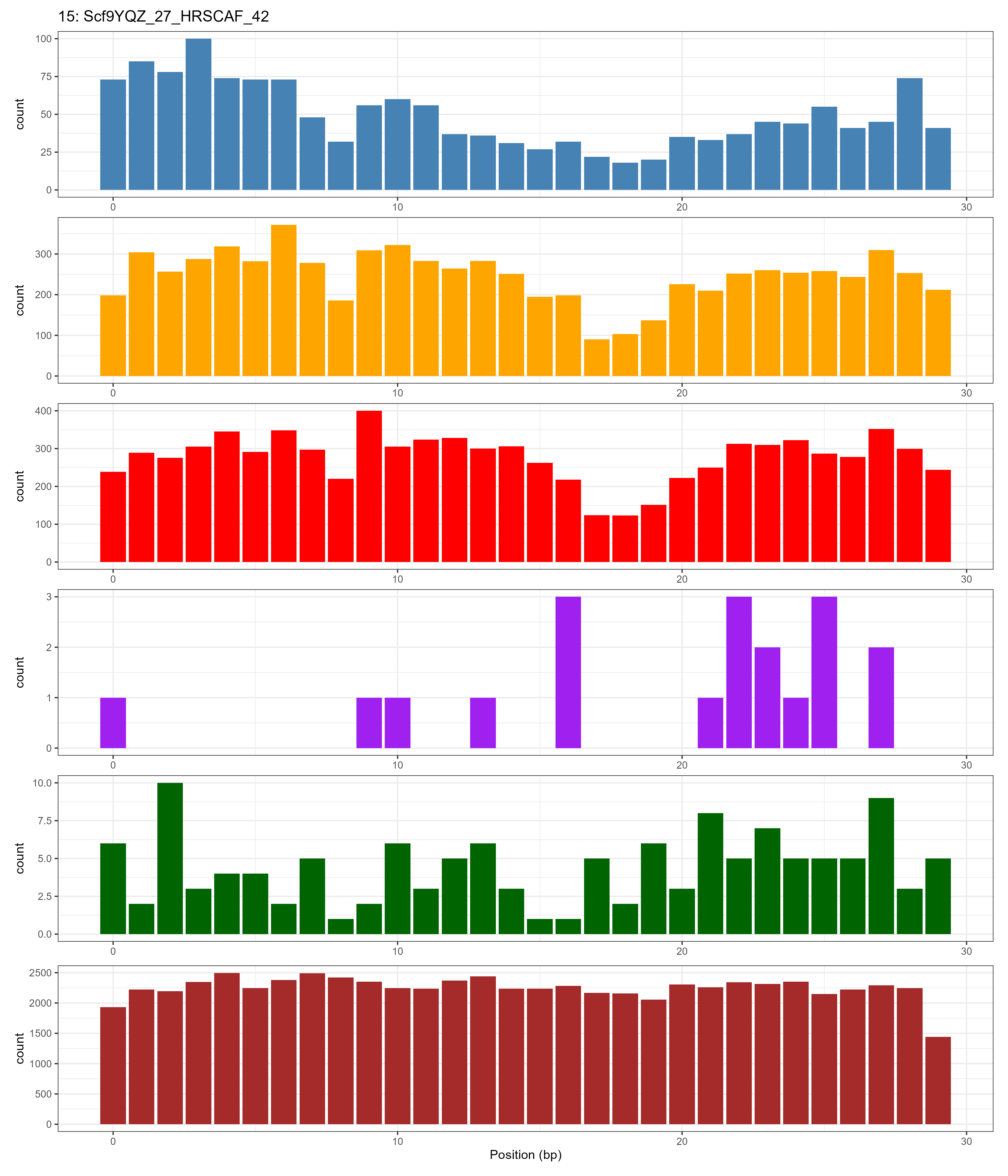

### Chromosome_16.png

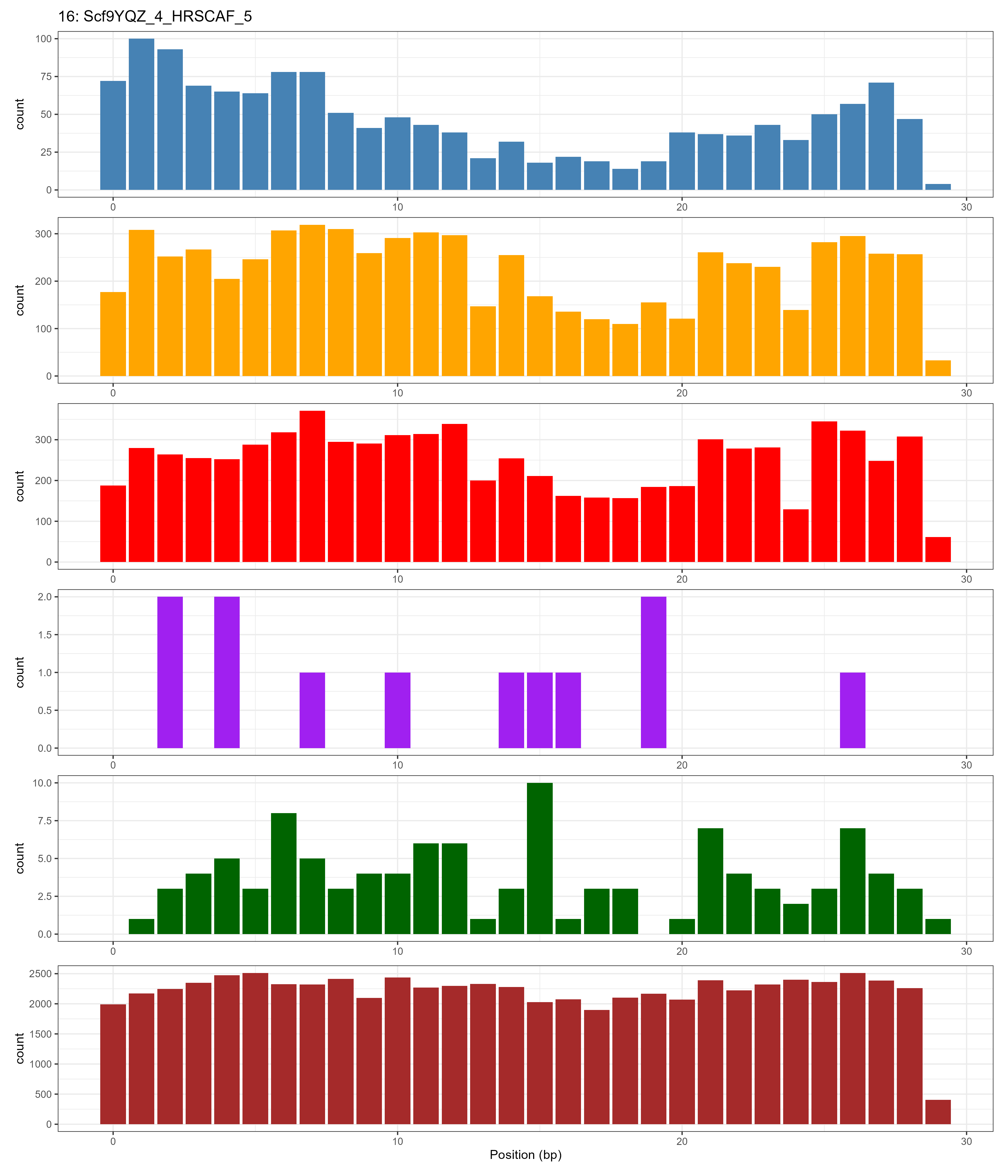

### Chromosome_17.png

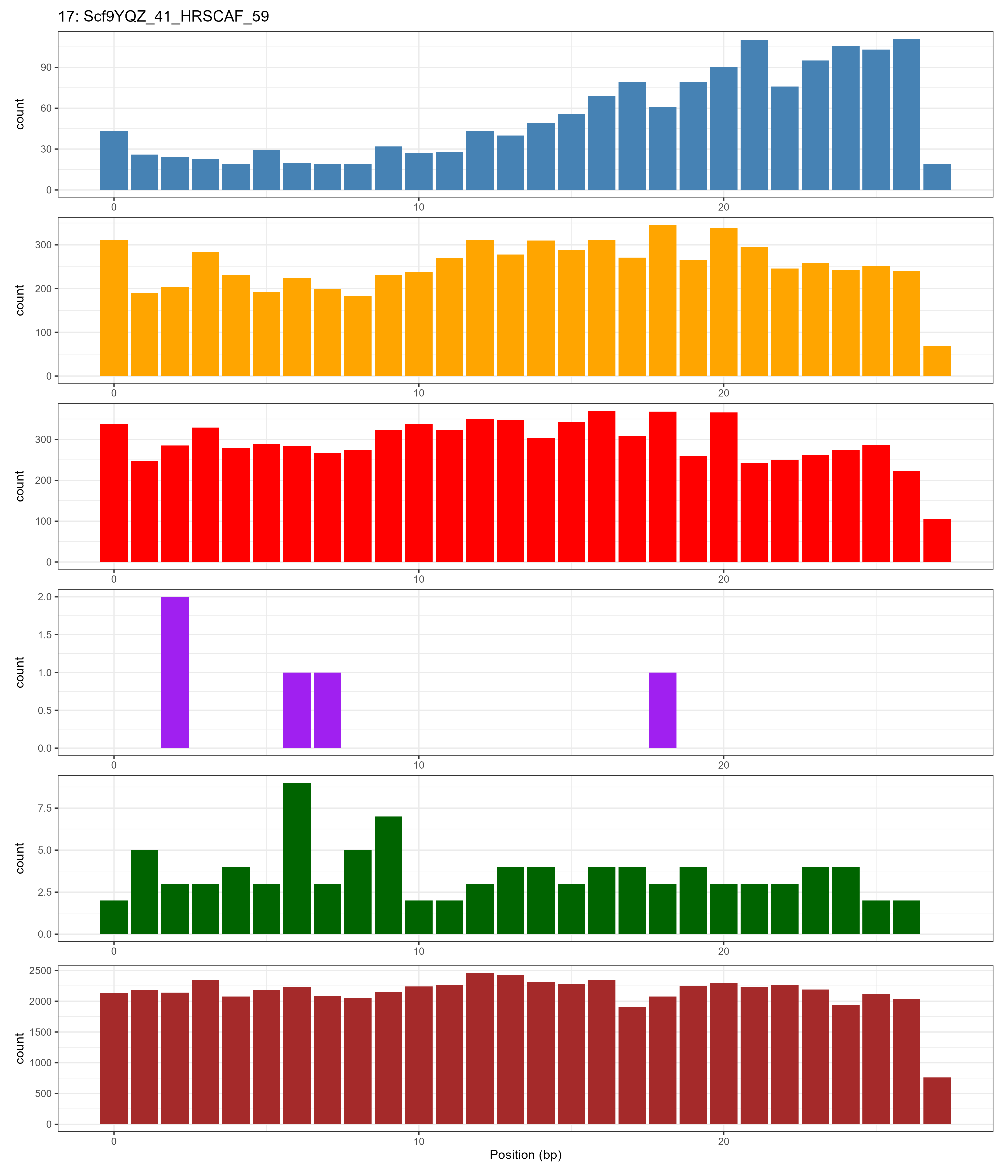

### Chromosome_18.png

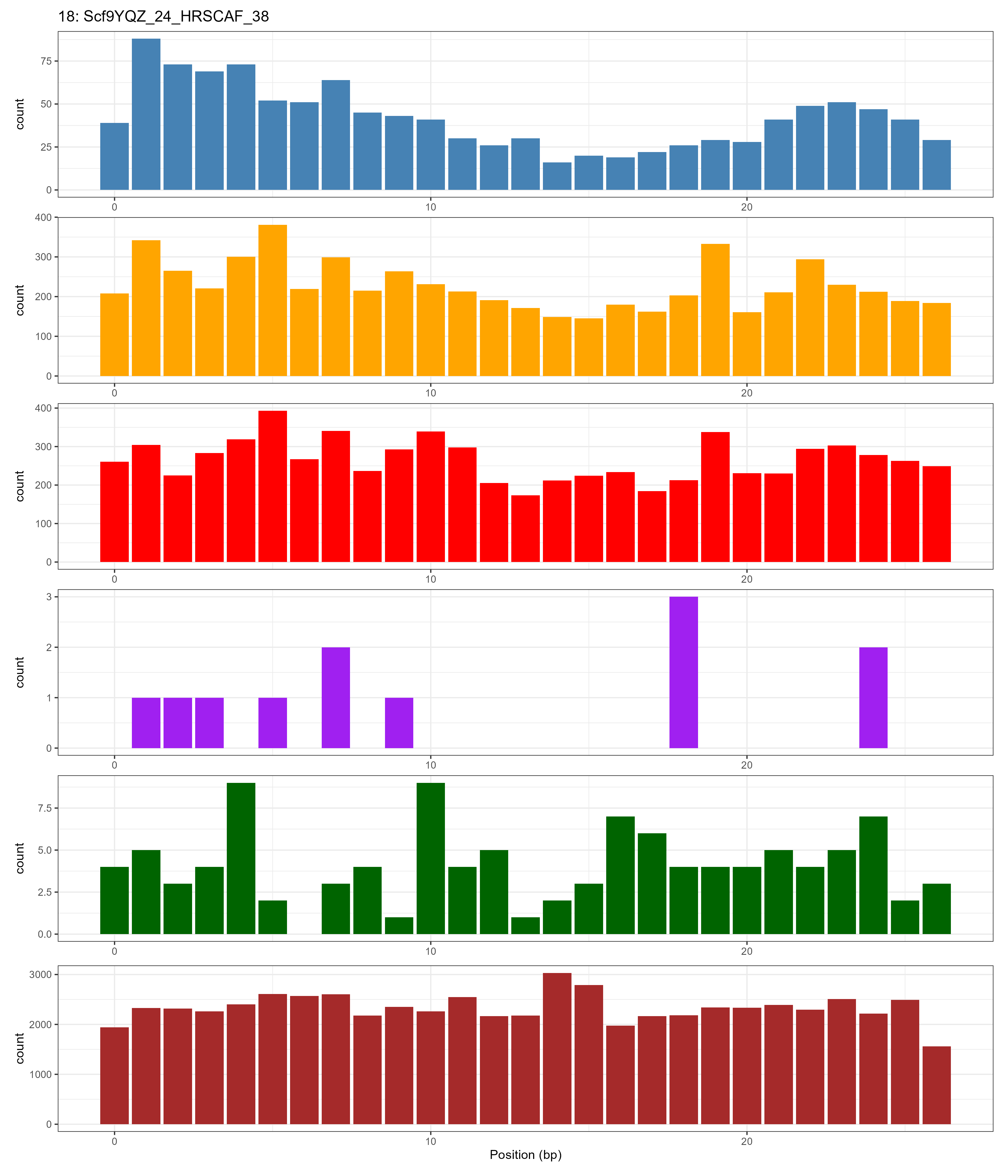

### Chromosome_19.png

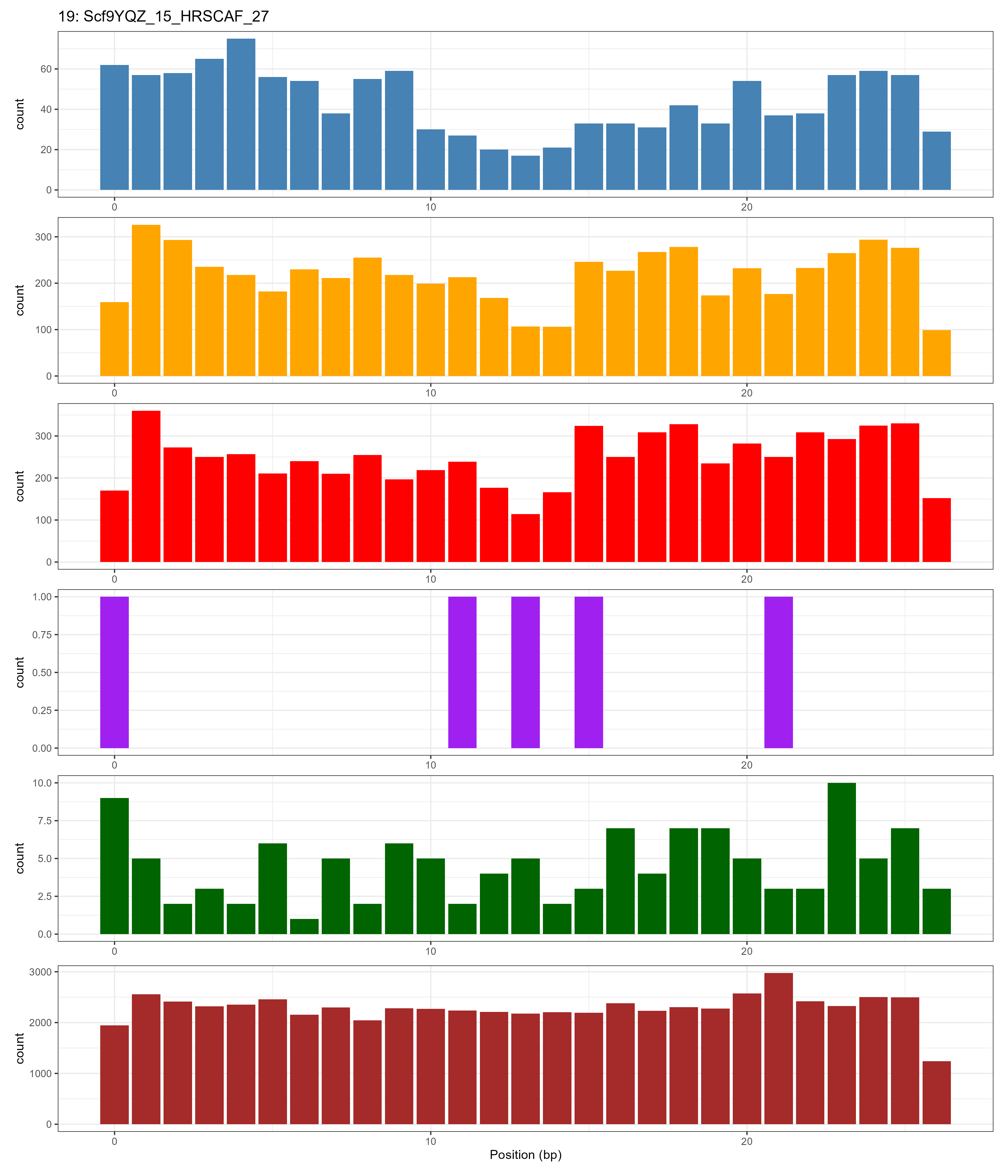

### Chromosome_20.png

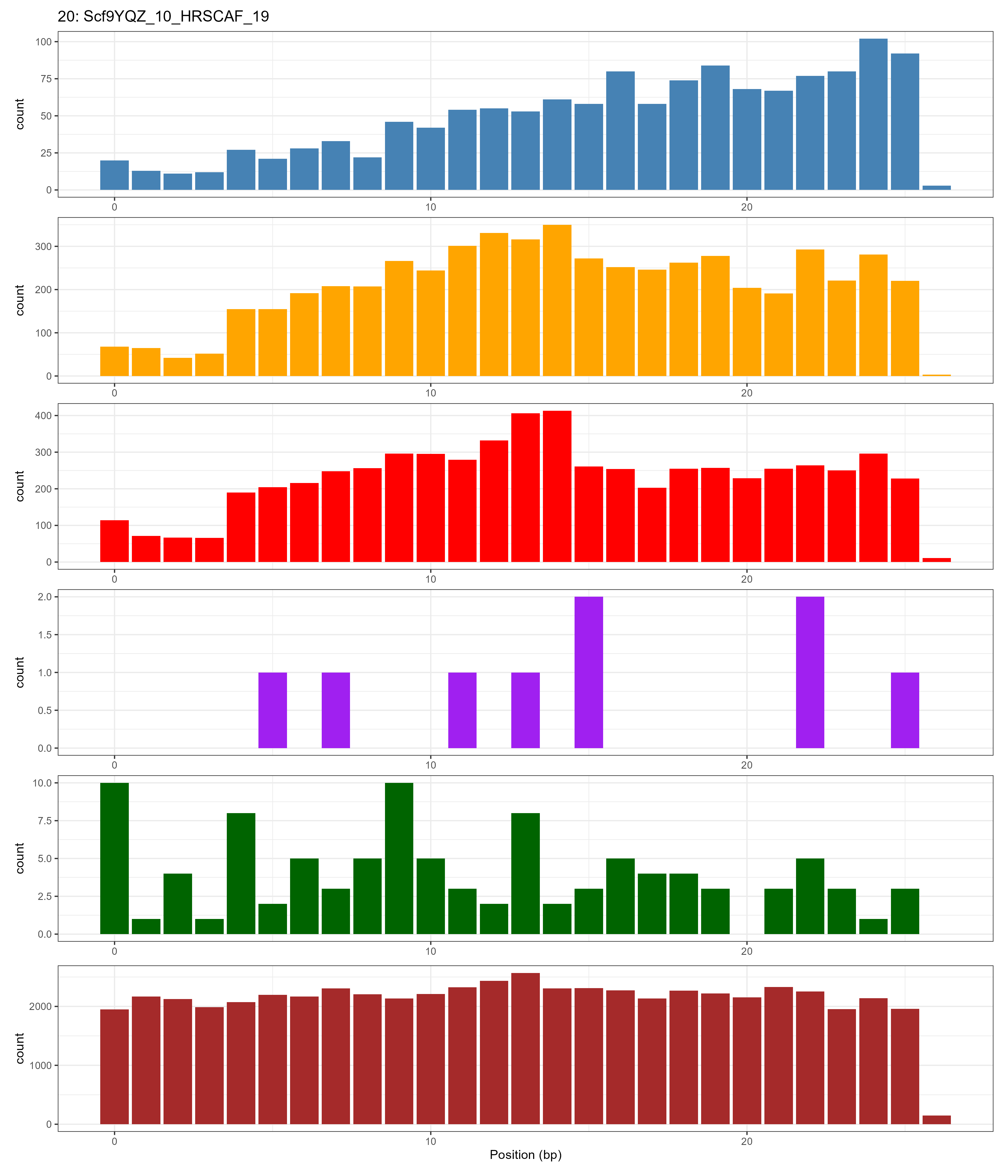

### Chromosome_21.png

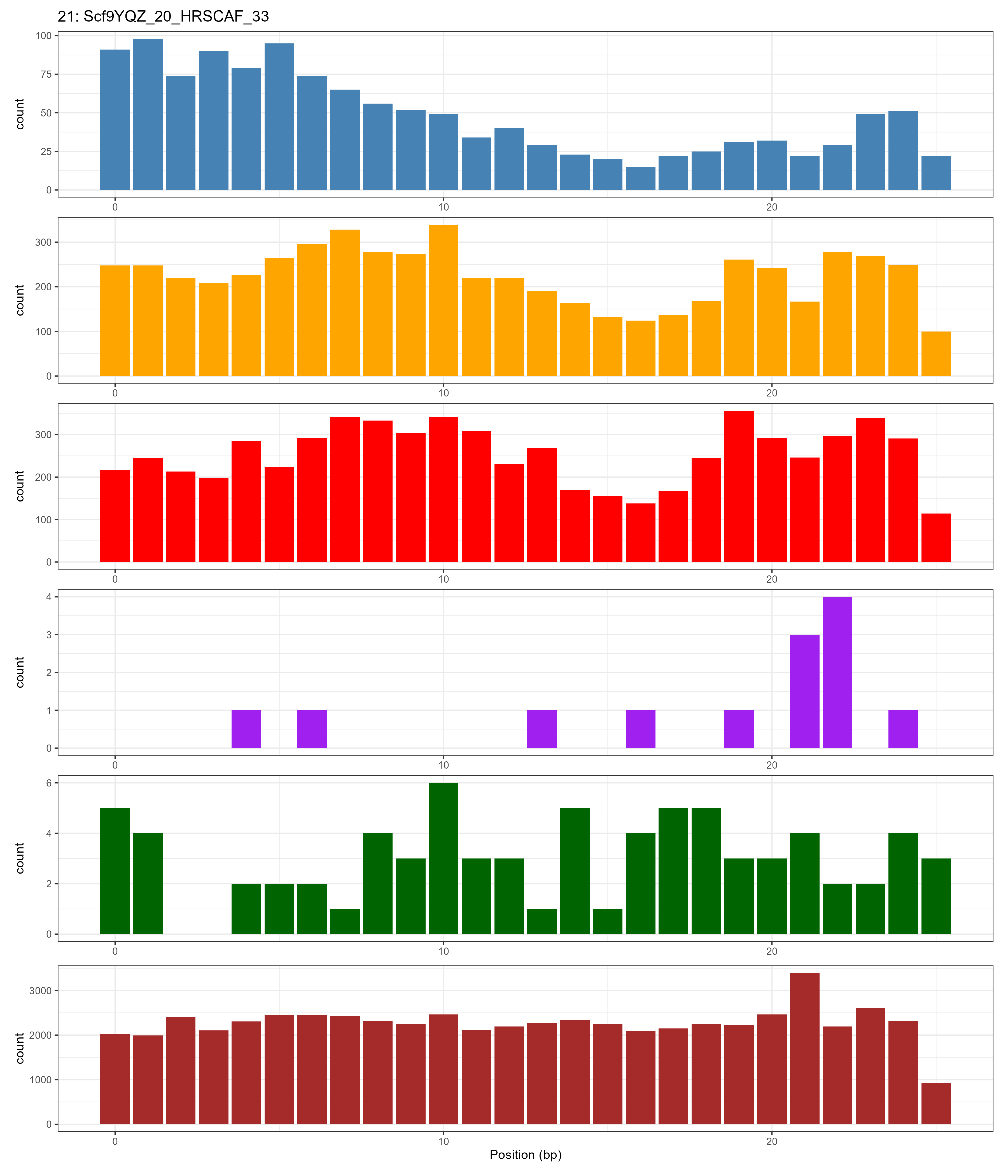

### Chromosome_22.png

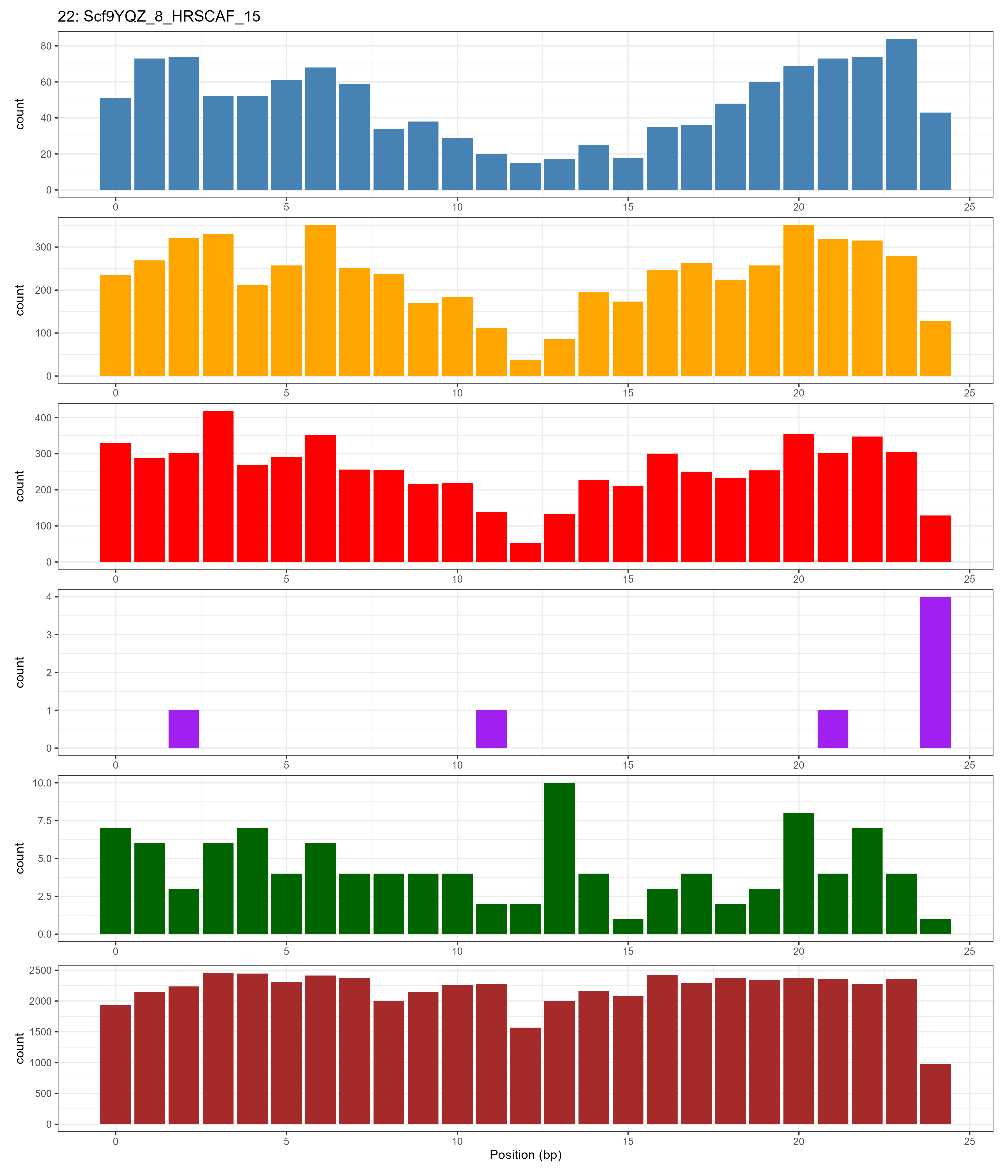

### Chromosome_23.png

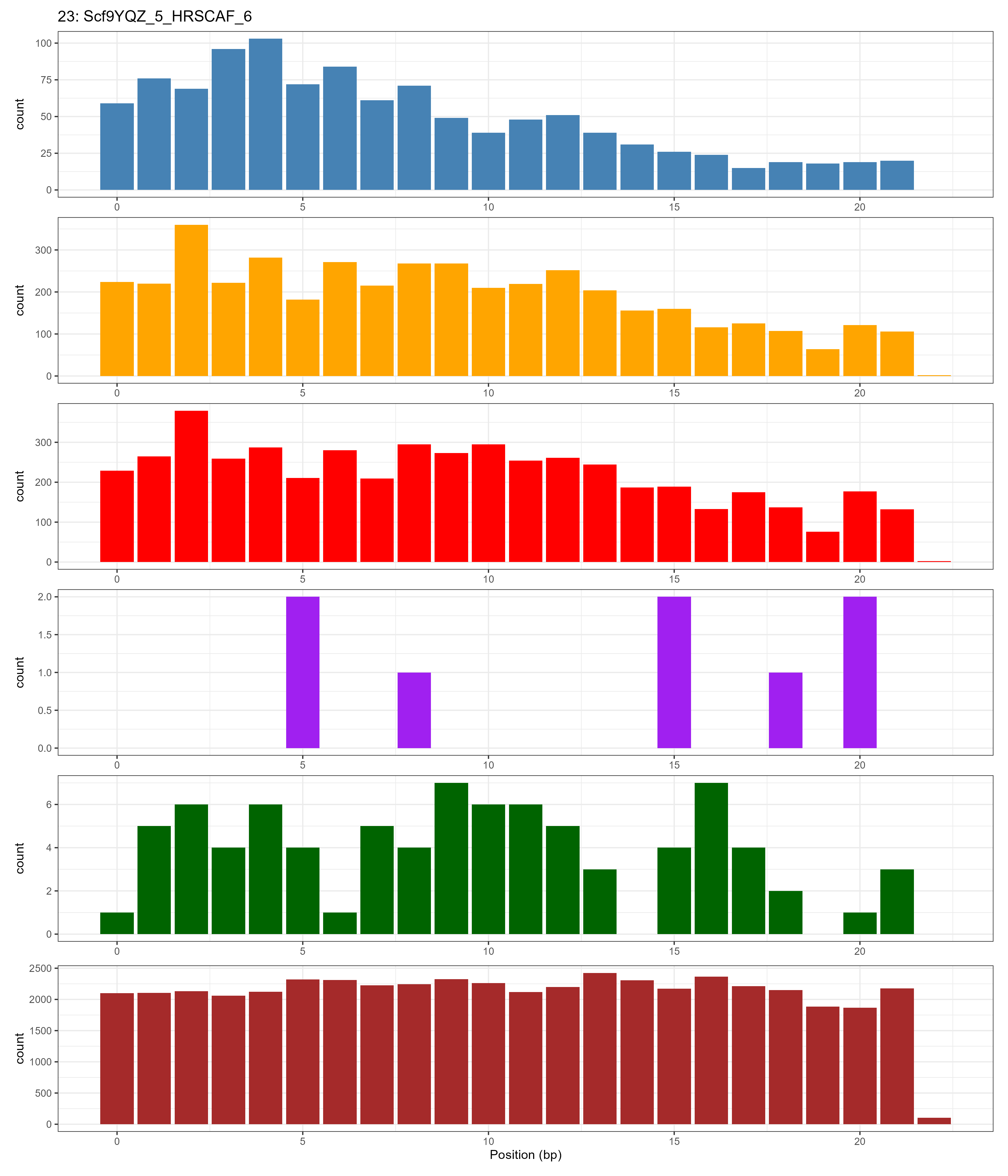
